# From Genes to Patterns: A Framework for Modeling the Emergence of Embryonic Development from Transcriptional Regulation

**DOI:** 10.1101/2024.09.19.613926

**Authors:** Jimena Garcia-Guillen, Ezzat El-Sherif

## Abstract

Understanding embryonic patterning, the process by which groups of cells are partitioned into distinct identities defined by gene expression, is a central challenge in developmental biology. This complex phenomenon is driven by precise spatial and temporal regulation of gene expression across many cells, resulting in the emergence of highly organized tissue structures. While similar emergent behavior is well understood in other fields, such as statistical mechanics, the regulation of gene expression in development remains less clear, particularly regarding how molecular-level gene interactions lead to the large-scale patterns observed in embryos. In this study, we present a modeling framework that bridges the gap between molecular gene regulation and tissue-level embryonic patterning. Beginning with basic chemical reaction models of transcription at the single-gene level, we progress to model gene regulatory networks (GRNs) that mediate specific cellular functions. We then introduce phenomenological models of pattern formation, including the French Flag and Temporal Patterning/Speed Regulation models, and integrate them with molecular/GRN realizations. To facilitate understanding and application of our models, we accompany our mathematical framework with computer simulations, providing intuitive and simple code for each model. A key feature of our framework is the explicit articulation of underlying assumptions at each level of the model, from transcriptional regulation to tissue patterning. By making these assumptions clear, we provide a foundation for future experimental and theoretical work to critically examine and challenge them, thereby improving the accuracy and relevance of gene regulatory models in developmental biology. As a case study, we explore how different strategies for integrating enhancer activity affect the robustness and evolvability of gene expression patterns in embryonic development. Our simulations suggest that a two-step regulation strategy, enhancer activation followed by competitive integration at the promoter, ensures more standardized integration of new enhancers into developmental GRNs, highlighting the adaptability of eukaryotic transcription. These findings provide new insights into the transcriptional mechanisms underlying embryonic patterning, offering a framework for future experimental and theoretical investigations.

## 1 Introduction

A fundamental challenge in developmental biology is understanding how a group of cells is partitioned into distinct identities, where each group is defined by the expression of one or a few specific genes (1–4). This process of cell diversification is further subdivided recursively, creating increasingly specialized subgroups, ultimately forming the complex and highly organized structure of the embryo with its diverse cell types. This phenomenon is known as embryonic patterning (2–9). Embryonic patterning relies on the precise regulation of gene expression—spatially and temporally—ensuring the right genes are activated in the right cells at the right time (1,2,10). This regulation controls how cells acquire their identities and organize into complex structures. Embryonic patterning, therefore, is an emergent phenomenon, resulting from the intricate regulation of many interacting genes across numerous cells. This concept of emergence parallels statistical mechanics, where complex behaviors in macroscopic systems, like gases or liquids, arise from interactions among many microscopic particles (11). Although the underlying laws for these particles, such as Newton’s laws or quantum mechanics, are relatively simple, their collective behavior leads to emergent large-scale phenomena. Similarly, in embryonic development, gene interactions across many cells produce the complex patterns guiding embryonic development (12,13). However, unlike well-understood models in statistical mechanics, our knowledge of gene regulation remains incomplete, particularly regarding how individual genes and their interactions drive these developmental patterns.

In emergent phenomena, not all features of the underlying microscopic elements are critical to the behavior of the larger macroscopic system. Similarly, in embryonic development, not all details of transcription at the single-gene level, which are often intricate (14), are essential contributors to the emergent patterns of gene expression that guide development. Therefore, a key step in understanding how gene regulation mediates embryonic patterning is to develop abstract, simplified models (15,16) that focus on the essential mechanisms at the single-gene level while still capturing the emergent gene expression patterns. However, we currently lack such models.

On one hand, reductionist approaches that delve into the molecular details of transcriptional machinery have uncovered vast amounts of information (10,17–19), but they often fail to clarify how these details contribute to the overarching process of pattern formation. On the other hand, existing models of gene regulation in embryonic pattern formation often rely on simplified systems, such as those found in bacteria (15,16,20–23). While these models provide insights, they fall short in capturing the much greater complexity of transcription in eukaryotes, especially in metazoans (24–27). This raises the question of which aspects of eukaryotic transcription are essential for mediating pattern formation. Identifying these key aspects, even at a theoretical level, would enable more targeted experimental work aimed at understanding the transcriptional mechanisms that drive embryonic patterning. To address this challenge, we need to establish a modeling framework that links gene regulation at the molecular level to the higher-level process of pattern formation across embryonic tissues. Such a framework should not simply aim to reproduce experimental data, but rather expose the underlying assumptions about how gene regulation occurs, providing new perspectives on the problem. Moreover, to enhance understanding, it is crucial to supplement these models with computer simulations, offering intuitive and straightforward code that researchers can easily use and modify.

In this paper, we propose such a framework by developing simple toy models of transcription and gene regulatory interactions at the single-gene level. Building on these models, we construct basic pattern formation models that generate realistic yet simplified gene expression patterns. Even if some of these models are based on outdated experiments in bacteria, this effort will clarify the assumptions driving our understanding of gene regulation, highlight areas for future investigation, and point to key differences between bacterial and eukaryotic transcription. By making these assumptions explicit, we aim to shed light on the connection between the molecular processes of transcription and the large-scale phenomenon of embryonic pattern formation, opening new avenues for exploration in developmental biology and developmental genetics. To enhance accessibility and encourage further research, we provide accompanying computer simulations with intuitive and simple code for each model.

We begin our modeling effort by focusing on the transcription process of single genes, using basic chemical reaction models. From there, we model how multiple genes interact to form gene regulatory networks (GRNs) that mediate specific functions within single cells. Next, we introduce phenomenological models of embryonic pattern formation, including various versions of the French Flag model and the Temporal Patterning/Speed Regulation models. These models are then extended with molecular or GRN realizations, representing a comprehensive modeling approach that connects transcriptional-level processes to tissue-level pattern formation. As a case study, we then explore the impact of different strategies for integrating the activities of multiple enhancers regulating a single gene on the robustness and evolvability of embryonic patterning. Our simulations suggest that the two-step gene regulation strategy observed in eukaryotes—first enhancer activation, followed by competitive integration of enhancer activities at the promoter—provides a more standardized approach for incorporating newly evolved enhancers into developmental GRNs. This insight emphasizes the evolutionary adaptability of eukaryotic transcriptional regulation and its role in shaping robust embryonic development.

## 2 Materials and methods

### 2.1 Modeling the regulation of a single gene

We begin our modeling of transcription during embryonic patterning by focusing on the kinetics of a single gene regulated by a single activator. Transcription initiation is typically controlled by a non-coding DNA sequence associated with the gene. In prokaryotes and simple eukaryotes, this regulatory region is a short DNA sequence located directly upstream of the coding sequence, near or overlapping the promoter, sometimes called the “operator”. Transcription factors (TFs) that regulate the gene bind to the operator, either aiding in the recruitment of RNA polymerase (in the case of activator TFs) or blocking it (in the case of repressor TFs), thereby controlling transcription (28).

In higher eukaryotes, transcription initiation is more complex (14) and often occurs in two stages. Regulatory TFs first bind to a non-coding region associated with the gene, known as an enhancer (10), which may be distant from the promoter or even located far from the gene itself. These enhancers influence the recruitment of general TFs to the promoter, where transcription is initiated. For simplicity, we will start our effort of modeling gene regulation by modeling the binding of a single activator to a single regulatory region. This region may be considered a functional combination of both the promoter and an enhancer, abstracting away the details of their interaction. Later in the paper, we will extend this model to explore the case of interactions between one or more enhancers and the gene promoter, capturing the more complex regulatory dynamics of transcription.

In this simple model of gene activation, an activator TF, denoted as X, binds to the regulatory region of a gene Y (Figure 1A). We represent the unbound regulatory region as *D*_0_and the region bound to X as *D*_1_. Transcription factors bind and unbind the regulatory region stochastically, with binding and unbinding rates denoted as *k*_1_ and *k*_−1_, respectively (Figure 1A), described by the equation:

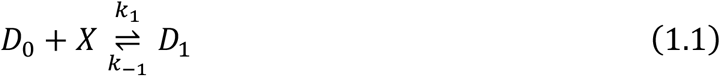

When the regulatory region is unbound (*D*_0_), transcription is inactive, whereas when X is bound (*D*_1_), the gene *Y* is transcribed at a rate *k*_*t*_, producing transcripts of *Y*, as described by the equation:

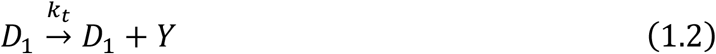

Transcripts are degraded at a rate *λ*, as modeled by the equation:

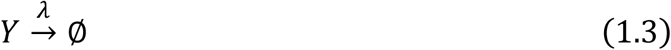

For simplicity, we assume that no transcription occurs in the absence of activator binding (i.e., no basal transcription).

The transcriptional activation of gene Y by TF X, described by Eqs. 1.1–1.3, can be translated into differential equations using the mass action law, as follows (For a detailed explanation of this conversion process, refer to Text S1, Figure S1, and Computer Simulations 1 and 2 (Text S2)).

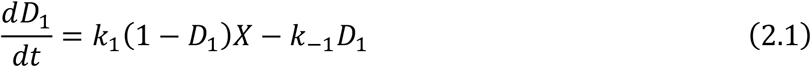

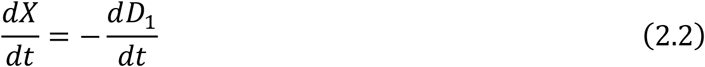

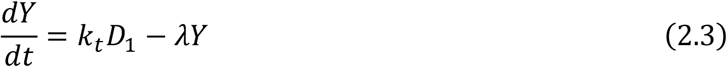

Here, *D*_1_ represents the fraction of DNA bound by X (with *D*_1_ + *D*_0_= 1). These three differential equations can be solved to determine (*t*), the concentration of gene Y transcripts over time (Fig. S2-A; Computer Simulation 3 in Text S2). However, because TF binding to DNA occurs on the millisecond timescale, while transcription proceeds on the order of seconds, we can apply the quasi-steady-state assumption. This assumption suggests that the steady-state values of X and *D*_1_ are reached much faster than the steady state of Y:

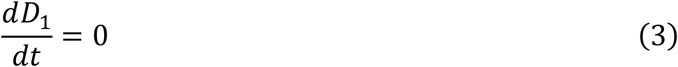

By substituting Eq. 3 into Eqs. 2.1–2.3, we reduce the system to a single equation, which takes the form of the Michaelis-Menten equation:

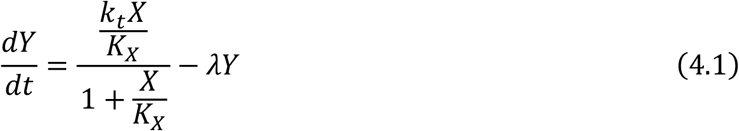

Here, *K*_*X*_ is the dissociation constant of TF X, representing the rate at which TF X detaches from DNA relative to its binding rate, and serves as a measure of the binding strength of X to DNA (lower values of *K*_*X*_ indicate stronger binding):

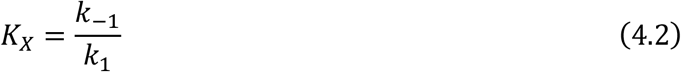

As demonstrated by Computer Simulations 3, and 4 (Text S2), the Michaelis-Menten equation (Eq. 4.1) yields results consistent with the three original equations when *k*_1_, *k*_−1_ ≫ *k*_*t*_ (compare the solution of the full equations describing gene activation, Eqs 2.1-2.3, vs Michaelis-Menten equation, Eq. 4.1; Figure S2-B and S2-C, respectively). To calculate the steady-state concentration of Y as a function of changing X, we set the transcription rate 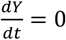, yielding the steady-state solution *Y*_*ss*_:

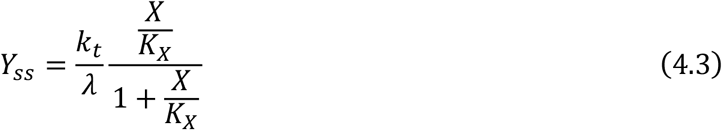

This results in a monotonic increase in steady-state transcription as the concentration of TF X increases, with saturation occurring at a maximum transcription rate of 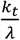 as X exceeds the dissociation constant *K*_*X*_ (see Computer Simulation 5 (Text S2), and simulation results in Figure 1B).

**Figure 1.**
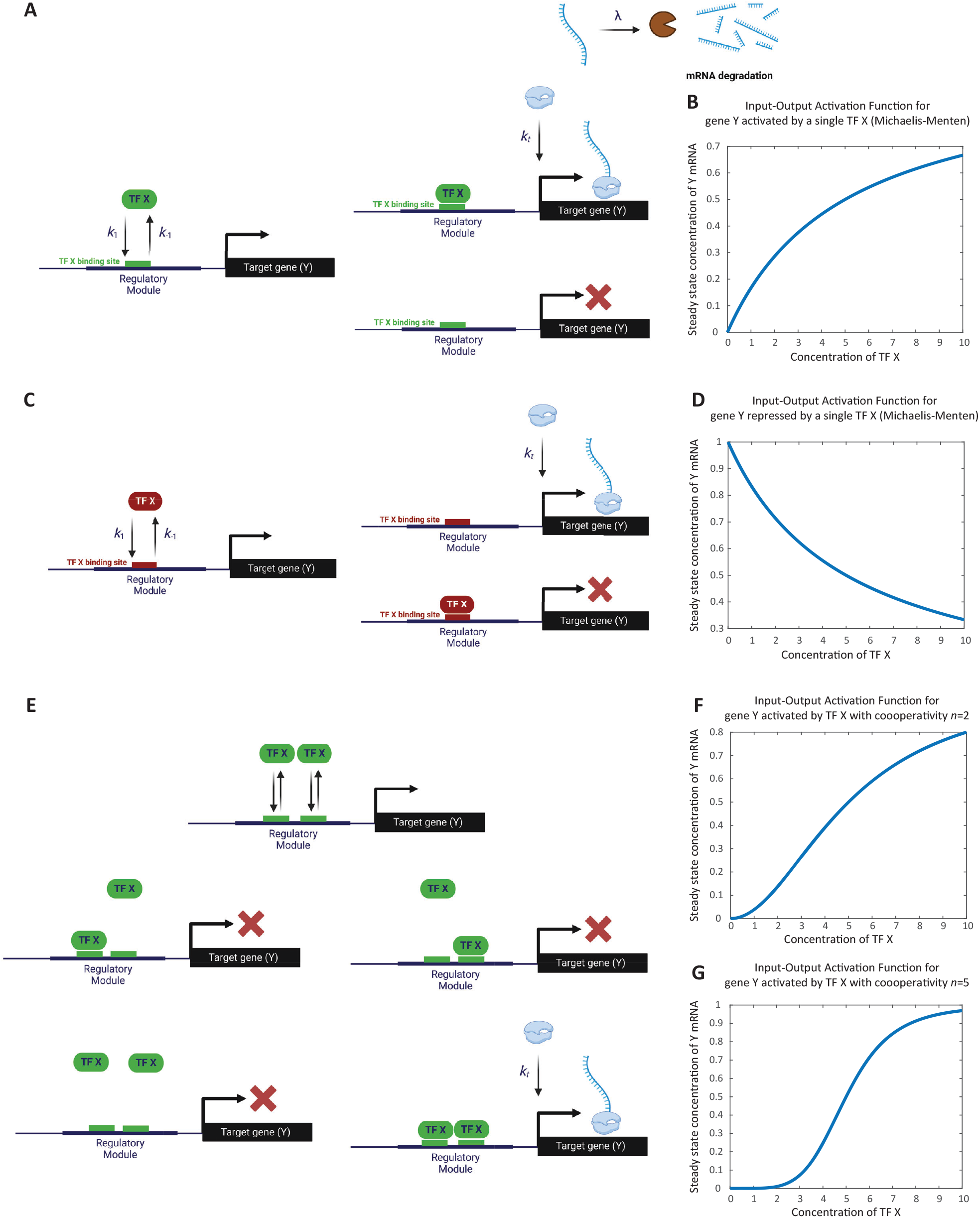
Modeling the regulation of a single gene. **(A)** Schematic representation of gene activation by a single activator transcription factor. The transcription factor X binds to the unbound regulatory region (D_0_) of gene Y, transitioning it to the bound state (D1). Binding and unbinding occur at rates k1 and k_−_1, respectively. Transcripts are degraded at rate λ. As an illustration, transcript degradation is shown here but will be omitted in subsequent schematics. This model demonstrates the fundamental biochemical reactions involved in transcription initiation. **(B)** Simulation of steady-state transcription levels as a function of activator concentration. The graph depicts the steady-state concentration of gene Y transcripts (Y*s*) increasing monotonically with activator X concentration. Saturation is observed when the concentration of X surpasses the dissociation constant (K_×_), resulting in a maximum transcription rate of k_t_/λ. This simulation corresponds to Computer Simulation 5 (Text S2). **(C)** Schematic representation of gene repression by a single repressor transcription factor. Repressor X binds to the regulatory region of gene Y, inhibiting transcription. The unbound state D_0_ transitions to the bound state D_1_ upon binding of X, with rates k_1_ and k_−1_ indicating binding and unbinding dynamics, respectively. **(D)** Simulation of steady-state transcription levels as a function of repressor concentration. The graph depicts the steady-state concentration of gene Y transcripts (Y*s*) decreasing as the concentration of repressor X increases. Maximum transcription occurs when X is zero, diminishing progressively with higher X concentrations. This simulation corresponds to Computer Simulation 6 (Text S2). **(E)** Modeling cooperativity in gene activation. Schematic representation of the cooperative binding of n copies of activator X to the regulatory region of gene Y. All n copies must bind simultaneously to activate transcription, representing extreme cooperativity. **(F)** Effect of cooperative binding on gene activation (*n* = 2). The simulation demonstrates how cooperative binding of two activator molecules leads to a switch-like increase in gene expression. The response curve shows a sharper transition compared to non-cooperative binding. This simulation corresponds to Computer Simulation 7 with *n*=2 (Text S2). **(G)** Effect of cooperative binding on gene activation (*n* = 5). The simulation illustrates that with higher cooperativity (*n* = 5), the gene’s response to activator concentration becomes more digital, exhibiting an even sharper switch-like behavior. This simulation corresponds to Computer Simulation 7 with *n*=5 (Text S2).

The proposed model of gene activation significantly oversimplifies the transcription process, particularly in eukaryotes, by neglecting critical aspects such as chromatin remodeling, nucleosome clearance, transcription pausing and release, transcriptional bursting, and other important details (14,29–34). Additionally, the quasi-steady-state (or equilibrium) assumption, along with the simplification of TF binding and unbinding as fast dynamics, overlooks how these factors might influence the slower overall dynamics of transcription (14). While this model’s simplicity has made it useful for gaining insights into gene regulation during development, assuming that slow dynamic processes have no effect on gene regulation may be misguided. As we will discuss in relation to combinatorial gene regulation, slow dynamics can indeed impact the overall transcriptional logic (35).

#### Thermodynamic State Ensemble (TSE) modelling

As shown above, deriving the Michaelis-Menten relationship from reaction equations can be quite tedious, particularly for more complex scenarios involving multiple TFs binding to the cis-regulatory region. A more streamlined approach for modeling transcriptional regulation that produces comparable results is the “Thermodynamic State Ensemble” (TSE) modeling (21,22,36). In this method, changes in transcript levels are influenced by the transcription rate (*T*_*y*_) and transcript degradation:

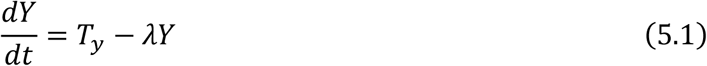

The transcription rate, *T*_*y*_, is expressed as the proportion of TF, DNA, and RNA polymerase configurations conducive to transcription relative to all possible configurations. Implicit in this approach the assumption that all processes involved in transcription are at equilibrium, and so it shares the quasi-steady state assumption as our modeling of gene activation presented above. Below, we outline how to use the TSE strategy to model gene regulation; for a more in-depth and rigorous explanation, readers are referred to the detailed exposition in (36).

As an example, we will use the TSE strategy to re-derive the rate equation governing the binding of a single activator X to the cis-regulatory region of gene Y. In this case, the denominator of *T*_*y*_ accounts for two possible states: unbound DNA (represented by 1) and DNA bound by X (represented by 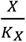, where *K*_*X*_ is the dissociation constant of X). The numerator includes only the states conducive to transcription—DNA bound by X, represented by 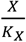. The overall expression is then multiplied by the transcription rate *k*_*t*_, which occurs once RNA polymerase binds to the DNA. This yields:

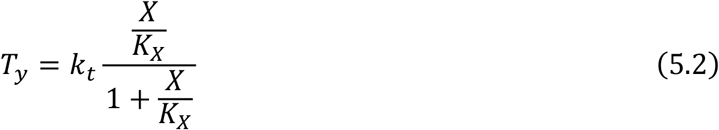

Substituting this into the differential equation for transcript dynamics gives the familiar Michaelis-Menten expression for a single activator (Eq. 4.1):

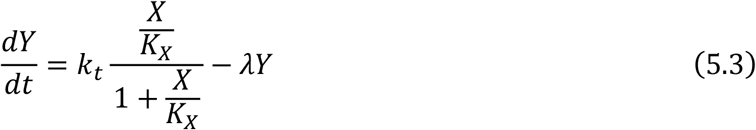

Next, we apply the TSE strategy to model gene regulation by a single repressor X (Figure 1C). In this case, the denominator of *T*_*y*_ remains similar to the activation case, but the numerator reflects only the state conducive to transcription—unbound DNA, represented by 1 (i.e., transcription occurs in the absence of repressor X). This yields the following equation for the transcription rate:

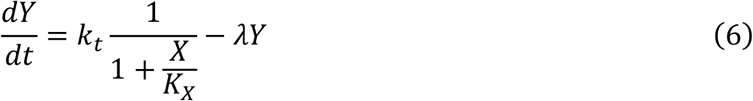

When plotting the steady-state value of Y as a function of X, the transcription rate reaches a maximum at zero concentration of the repressor X, then progressively decreases as X concentration increases (see Computer Simulation 6 (Text S2), which results are depicted in Figure 1D).

#### Modeling cooperativity

Often, multiple copies of the same TF are required to bind cooperatively in order to effectively recruit RNA polymerases. In this section, we will use the TSE approach to model the cooperative binding of *n* copies of activator TF X for the activation of gene Y. We will assume a case of extreme cooperativity, meaning that all *n* copies of X must bind simultaneously to the regulatory region of Y, whereas fewer than *n* copies are unable to bind.

In this scenario, the denominator of *T*_*x*_ represents all possible binding events, which include unbound DNA and the binding of *n* copies of X to the cis-regulatory elements. The numerator, however, includes only the state conducive to gene activation, which is the simultaneous binding of *n* copies of X to the regulatory region of Y (see Figure 1E). This yields the rate equation:

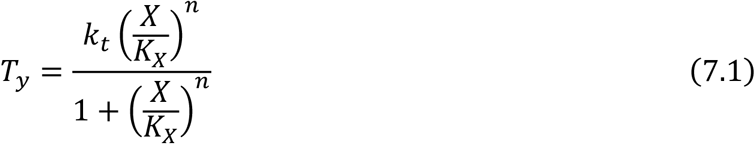

This type of equation is commonly referred to as the Hill equation.

Cooperativity introduces a nonlinear, switch-like behavior in the gene’s response to activator binding. The higher the value of *n*, the more pronounced this switch-like behavior becomes, with a sharper transition from low to high input. As *n* increases further, the gene’s response approaches digital behavior, as demonstrated by comparing the cases of *n* = 2 and *n* = 5 in Figures 1F and 1G, respectively (see Computer Simulation 7 (Text S2)). The point at which the gene switches from low to high output is determined by the dissociation constant *K*_*X*_, which characterizes the binding affinity of TF X (as shown in Eq. 7.1).

Similarly, the Hill equation for the cooperative binding of *n* repressors follows a comparable form:

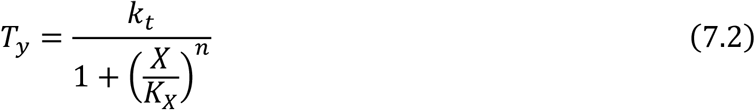

In both cases, the cooperative binding of TFs leads to a sharp, switch-like regulation of gene expression, reflecting the impact of cooperativity on transcriptional control.

#### Modeling multiple TFs binding cis-regulatory module

So far, we have considered the binding of one or multiple copies of the same transcription factor (either activator or repressor) to the regulatory region of a gene. In this section, we will extend this by considering the binding of multiple different TFs. Specifically, we will examine the binding of two TFs, A and B, to the regulatory region of gene Y. This can be extended to cases involving a greater number of different TFs with straightforward modifications.

The effect of binding multiple TFs depends on the logic that governs their regulation of the gene, often referred to as the “transcription logic” of the regulatory module. Below, we will explore several common transcriptional logic models, though more elaborate scenarios are certainly possible.

##### Two ORed Activators

In this model, the binding of either A, B, or both activates the gene (Figure 2A). Using the TSE approach, and assuming there is no cooperative binding between A and B (i.e., the binding of A does not influence the binding of B and vice versa), we obtain the following equation:

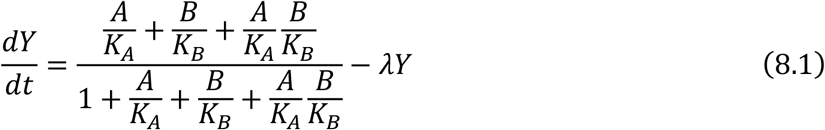

In cases where A and B bind cooperatively, with cooperativity values *n*_*A*_ and *n*_*B*_, we derive the following Hill equation:

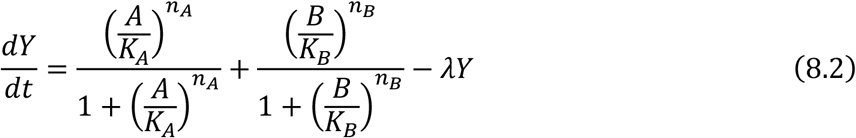

As shown in Figure 2B, simulation result of this relationship matches the OR logic as expected (see Computer Simulation 8 (Text S2)). Moving forward, we will focus on the binding of single copies of A and B, as the extension to cooperative binding is straightforward.

**Figure 2.**
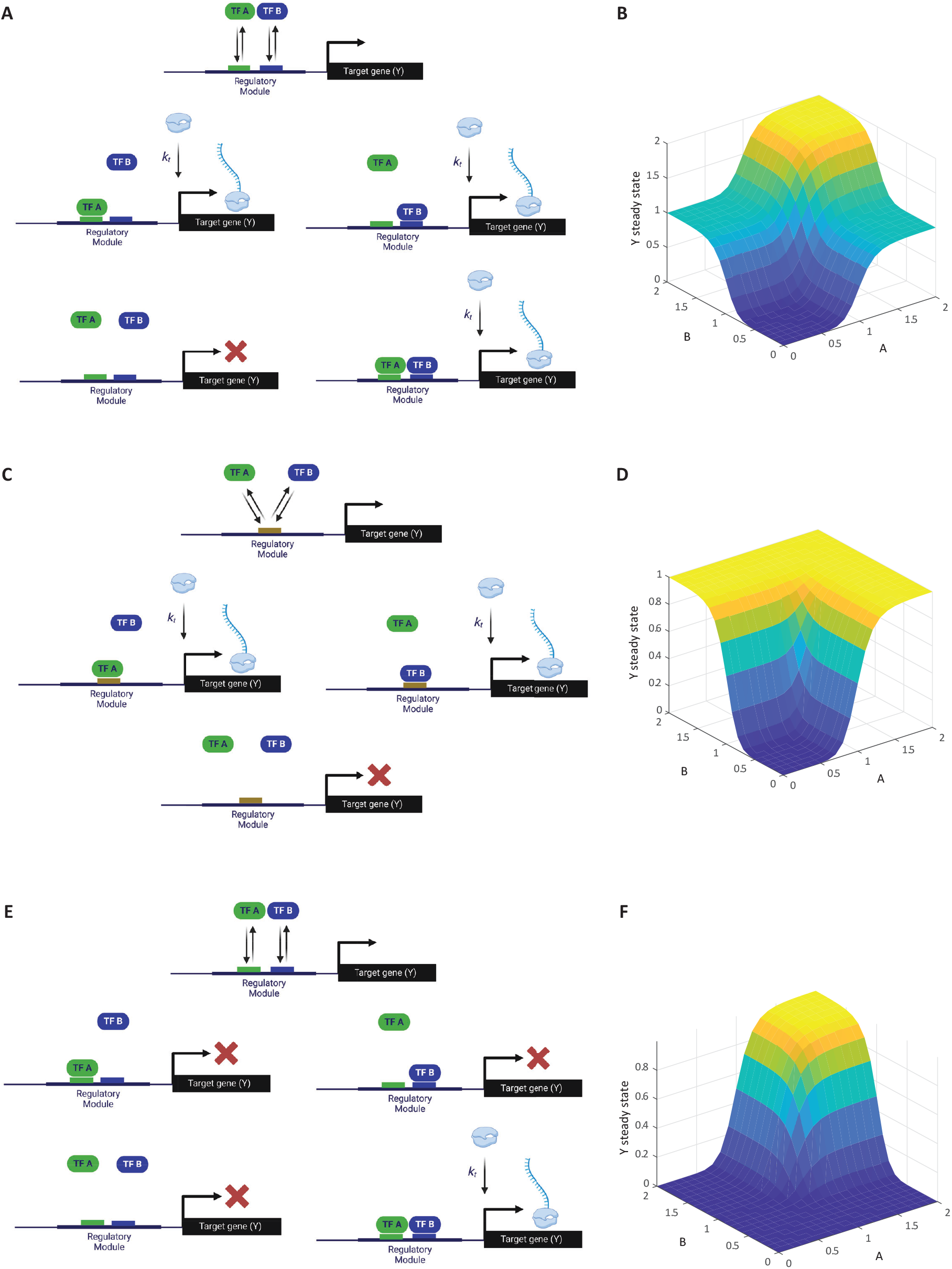
Modeling multiple transcription factors binding to a cis-regulatory module. **(A)** Schematic of gene activation by two ORed activators. Transcription factors A and B independently bind to the regulatory region of gene Y. The presence of either A or B is sufficient to activate transcription, illustrating an OR logic gate. **(B)** Simulation of transcription levels with two ORed activators. The graph shows how the steady-state concentration of gene Y transcripts varies with different concentrations of A and B, reflecting the OR logic. This simulation corresponds to Computer Simulation 8 (Text S2). **(C)** Schematic of gene activation by two competitive ORed activators. Activators A and B compete for the same binding site on gene Y’s regulatory region. Binding of either A or B activates transcription, but they cannot bind simultaneously due to competition. **(D)** Simulation of transcription levels with competitive ORed activators. The graph illustrates how the competition between A and B affects gene Y’s expression, with transcription levels depending on the relative concentrations of A and B. This simulation corresponds to Computer Simulation 9 (Text S2). **(E)** Schematic of gene activation by two ANDed activators. Both transcription factors A and B must simultaneously bind to gene Y’s regulatory region to initiate transcription, representing an AND logic gate. **(F)** Simulation of transcription levels with ANDed activators. The graph demonstrates the switch-like activation of gene Y only when both A and B are present, showcasing the cooperative effect of AND logic on gene expression. This simulation corresponds to Computer Simulation 10 (Text S2).

##### Two Competitive ORed Activators

Here, we consider the case where both A and B are activators that bind the same site on the regulatory region of gene Y, leading to competitive binding (Figure 2C). In this case, the denominator includes unbound DNA, DNA-bound A, and DNA-bound B. However, the simultaneous binding of both A and B is excluded because they compete for the same binding site and cannot bind at the same time. The numerator includes the states where A or B binds independently, resulting in the reaction equation:

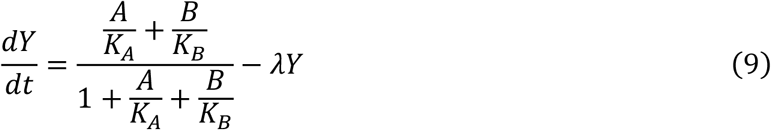

As expected, simulation result of the Hill-function extension of this relationship matches the competitive OR logic as described above (Figure 2D; see Computer Simulation 9 (Text S2)).

#### Two ANDed Activators

In this model, both A and B must bind simultaneously to activate gene expression (Figure 2E). This logic results in the following equation:

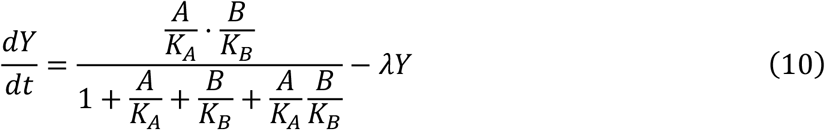

As expected, simulation result of the Hill-function extension of this relationship matches the competitive AND logic (Figure 2F; see Computer Simulation 10 (Text S2)).

##### Two ANDed Repressors

So far, we have considered activators, but the extension to models involving both activators and repressors is straightforward. As an example, we will examine the case of two ANDed repressors. In this case, only unbound DNA allows for transcription, and the transcription rate *T*_*x*_ is the product of the individual repression effects of A and B:

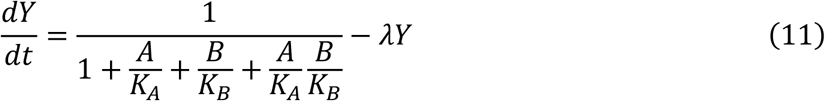

The transcriptional logics presented above represent relatively simple cases. However, more complex scenarios could be modeled within the TSE framework by incorporating cooperative binding between different TFs (24). However, the modeling approach presented here shares the same limitations as our earlier approach using rate equations (Eqs. 1.1-1.3 and 2.1-2.3), as it neglects many details of the transcription cycle in eukaryotes. Additionally, TSE models assume that binding and unbinding events are at equilibrium, similar to the quasi-steady state assumption used in the rate equations (Eqs. 3 and 4.1-4.3). However, the rate equation framework (Eqs. 1.1-1.3 and 2.1-2.3) is less constrained by assumptions and can, in principle, accommodate more transcriptional details. Employing rate equations to capture these complexities has proven effective in providing insights into gene regulation in eukaryotes, revealing a range of potential mechanisms for achieving combinatorial transcriptional logic beyond those typically assumed within TSE models (35). Nevertheless, for simplicity, we will continue with the TSE modeling results in this study.

### 2.2 Modeling gene regulatory networks

So far, we have considered the regulation of single genes by one or more TFs. During development, terminal and housekeeping genes may be passively regulated by other transcription factors. However, many TFs regulate other TFs, forming gene regulatory networks (GRNs) (Figure 3A) (13). In this section, we will explore examples of GRNs that act within a single cell and how to model them. Later, we will extend this to GRNs that function across multiple cells within embryonic tissues.

**Figure 3.**
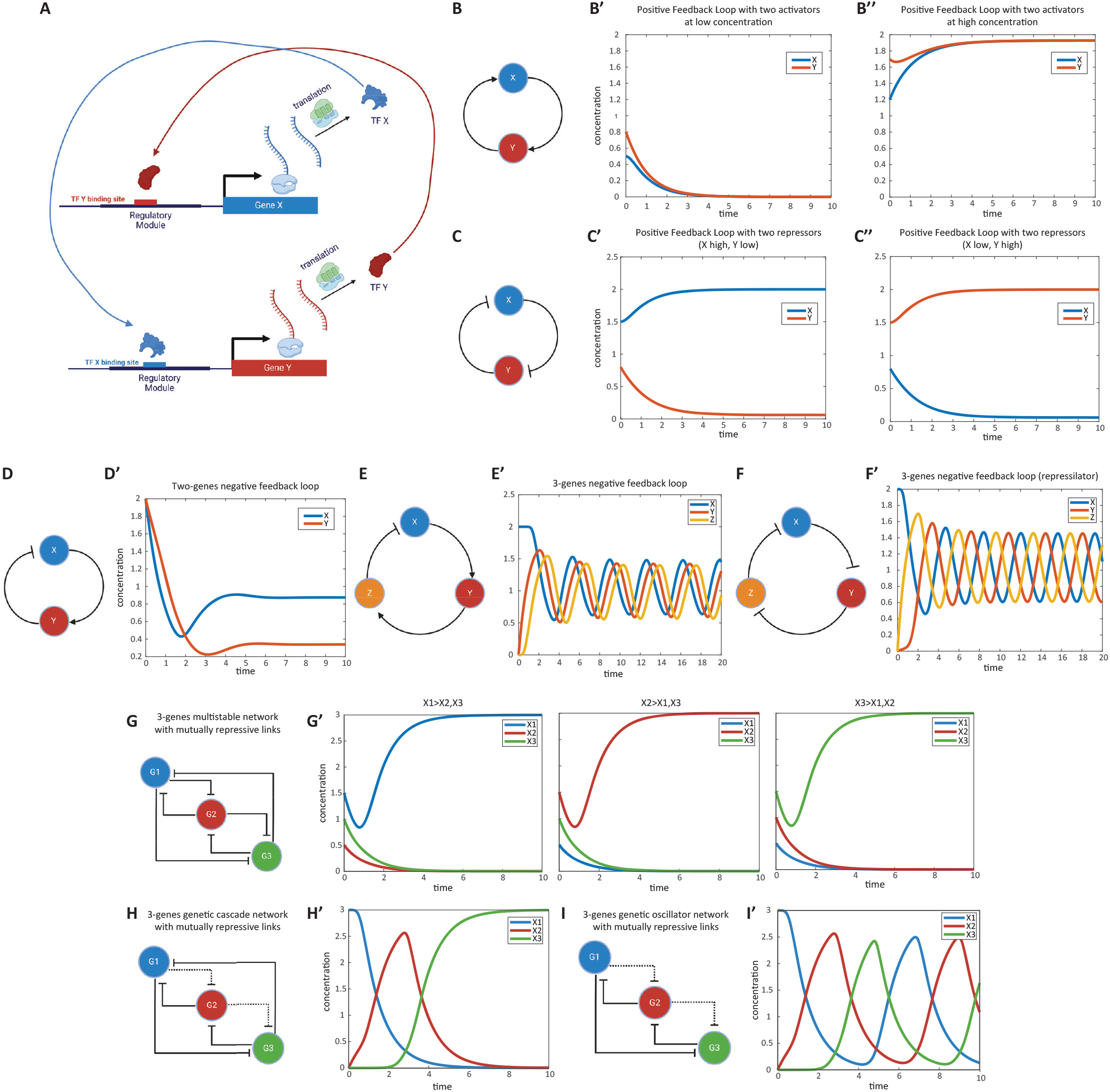
Modeling gene regulatory networks (GRNs). **(A)** Schematic representation of a gene regulatory network (GRN). Multiple genes interact to regulate each other’s expression, forming complex networks that mediate specific cellular functions. Shown is a positive feedback loop between two genes as an example. **(B)** Positive feedback loop between two activator genes. Genes X and Y activate each other’s expression. This mutual activation can lead to bistable states where both genes are either on or off. **(B’)** Simulation of gene expression states in the positive feedback loop. The graph shows stable states achieved in the system, with both genes being not expressed or highly expressed, depending on initial conditions. This simulation corresponds to Computer Simulation 11 (Text S2). **(C)** Positive feedback loop between two repressor genes. Genes X and Y repress each other’s expression. This mutual repression results in a system where one gene is active while the other is repressed. **(C’)** Simulation of gene expression states in the mutual repression loop. The graph demonstrates the system stabilizing with one gene active and the other inactive, based on initial expression levels. This simulation corresponds to Computer Simulation 12 (Text S2). **(D)** Negative feedback loop between an activator and a repressor gene. Gene X activates gene Y, while gene Y represses gene X. This configuration promotes homeostasis, maintaining stable gene expression levels. **(D’)** Simulation of gene expression in the negative feedback loop. The graph shows how the system maintains stable expression levels of genes X and Y over time. This simulation corresponds to Computer Simulation 13 (Text S2). **(E)** Oscillatory GRN with two activators and one repressor. Gene X activates gene Y, gene Y activates gene Z, and gene Z represses gene X. This network can produce oscillations in gene expression over time. **(E’)** Simulation of oscillatory behavior in the gene network. The graph displays cyclic fluctuations in the expression levels of genes X, Y, and Z, characteristic of an oscillatory system. This simulation corresponds to Computer Simulation 14 (Text S2). **(F)** Repressilator network composed of three repressor genes. Genes X, Y, and Z repress one another in a cyclic manner, forming an oscillatory system known as a repressilator. **(F’)** Simulation of oscillations in the repressilator network. The graph shows the periodic expression patterns of genes X, Y, and Z over time. This simulation corresponds to Computer Simulation 15 (Text S2). **(G)** Multi-stable gene regulatory network with mutually repressing genes. Multiple genes repress each other strongly, leading to stable states where only one gene is expressed while the others are repressed. **(G’)** Simulation of gene expression states in the multi-stable network. The graph shows that depending on initial conditions, one gene remains active while others are inactive. This simulation corresponds to Computer Simulation 16 (Text S2). **(H)** Genetic cascade illustrating sequential gene activation. Each gene in the cascade represses the previous gene and weakly represses the next one. Activation of the first gene triggers a domino effect of gene expression. **(H’)** Simulation of gene expression in the genetic cascade. The graph depicts the sequential activation and repression of genes over time, following the cascade logic. This simulation corresponds to Computer Simulation 17 (Text S2). **(I)** Oscillatory gene network in which a series of genes repress each other in sequence without the last gene repressing the first, resulting in continuous oscillations of gene expression. **(I’)** Simulation of oscillations in the gene network. The graph shows the cyclic expression patterns of the genes over time. This simulation corresponds to Computer Simulation 18 (Text S2).

#### Positive Feedback Loop

We begin by considering feedback loops involving two genes. If activation is positive and repression is negative, a positive feedback loop results when the product of all regulatory links’ signs is positive (15,16,37). Positive feedback loops can function as toggle switches or memory devices—if one of the genes is activated, it remains active.

A positive feedback loop can be realized as either two genes activating each other (Eqs. 12.1, 12.2; Figure 3B) or two genes repressing each other (Eqs. 13.1, 13.2; Figure 3C). In a loop of two activators, both genes can be either high (“on”) or low (“off”) (Figures 3B’, 3B”; see Computer Simulation 11 (Text S2)). In a loop of two repressors, the GRN stabilizes in a state where one gene is active and the other is repressed (Figures 3C’, 3C”; see Computer Simulation 12 (Text S2)). In these equations, *K*_*XY*_ denotes the dissociation constant of TF Y binding to the regulatory region of gene X.

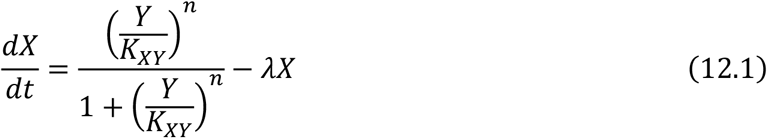

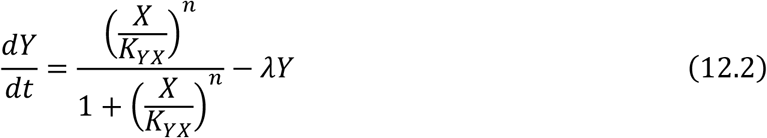

For a repression-based positive feedback loop, the equations become:

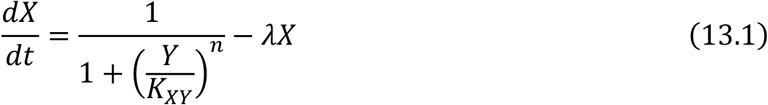

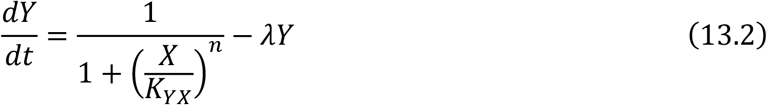

#### Negative Feedback Loop

A two-gene negative feedback loop (Figure 3D) can help maintain homeostasis, ensuring that gene expression remains stable around set values. For instance, the GRN modeled by Eqs. 14.1–14.2 maintains gene Y’s transcriptional activity around the activation threshold for TF X binding to its regulatory region (*K*_*YX*_), and gene X’s activity around the deactivation threshold for TF Y binding to its regulatory region (*K*_*XY*_) (Figure 3D’; see Computer Simulation 13 (Text S2)).

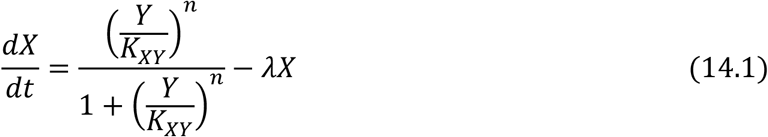

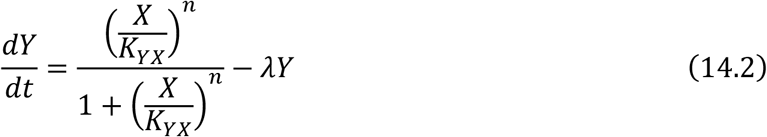

### Oscillatory GRNs

While two-gene negative feedback loops typically maintain homeostasis, more complex three-gene negative feedback loops can result in oscillations, depending on network parameters. For example, a GRN with two activators and one repressor (Figure 3E, E’; see Computer Simulation 14 (Text S1)) can behave as an oscillator (Eqs. 15.1–15.3). A GRN composed of three repressors, also known as a repressilator (38), can also function as an oscillator (Figure 3F, F’; see Computer Simulation 15 (Text S1)).

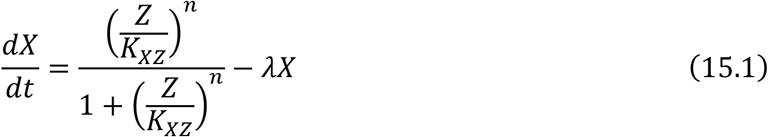

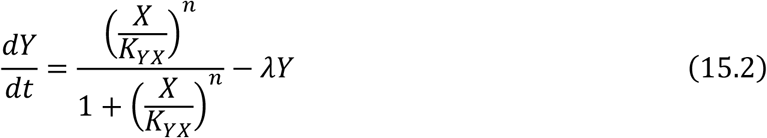

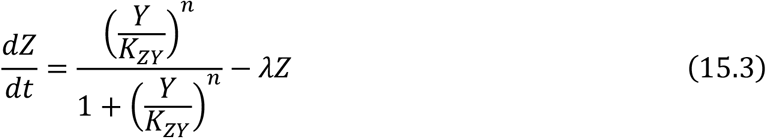

#### Developmentally inspired GRNs

So far, we have presented example GRNs with specific wiring capable of executing various functions within the cell. The structures we’ve discussed are not the only possible configurations, as many different GRN architectures with varying parameter values can accomplish the same function. However, GRNs involved in development, particularly those involved in embryonic pattern formation, have distinct characteristics. One notable feature is the minimal reliance on activators; most TFs involved in embryonic patterning are repressors, where activation is achieved through the repression of a repressor, a process called de-repression (39,40). For example, most of the patterning genes involved in the patterning of the AP and DV axes in the early *Drosophila* embryo are transcriptional repressors (41–43).

We will now explore some common GRN architectures that perform specific tasks during embryonic pattern formation, relying on repression and de-repression to mediate regulatory relationships between genes. Another important feature of these GRNs is the extensive cross-regulation between the genes, where oftentimes all constituent genes repress each other. Despite this complexity, variations in the strength of repression among the genes can produce meaningful and distinct functions, as illustrated by the GRNs discussed below.

#### Multi-stable GRNs

A multi-stable GRN (40) functions similarly to the positive feedback loop or genetic toggle switch discussed earlier. However, developmental GRNs typically consist of more than two genes. The simplest extension of the toggle switch to more than two genes is a multi-stable GRN, composed of M mutually repressing genes. This repression is characterized by small dissociation constants (*θ*_*s*_), indicating strong repression (Eq. 16; Fig. 3G). In this configuration, depending on the initial conditions, one gene remains active while the others are inactive (Fig. 3G’; see Computer Simulation 16 (Text S2)). Other variations of this realization are possible, where subsets of genes are co-expressed due to weakened repressive links between them (44).

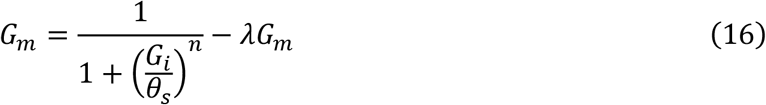

#### Genetic Cascade

Another developmentally inspired GRN is the genetic cascade, where a sequence of genes is activated one after another. A simple realization of this cascade involves one gene activating the next, which in turn represses the previous gene, and so on. In a GRN based on de-repression logic (39,44,45), all genes are naturally active but kept repressed by other genes in the cascade (with small dissociation constants, *θ*_*s*_, indicating strong repression; Eqs. 17.1, 17.2). The only exception is that a gene weakly represses the gene preceding it in the cascade (with larger dissociation constants, *θ*_*w*_, indicating weaker repression) (Fig. 3H). When the cascade is initialized by activating the first gene, a domino effect occurs where the next gene represses the previous one, leading to the de-repression of the subsequent gene, and so forth (Fig. 3H’; see Computer Simulation 17 (Text S2)).

For *m* = 1:

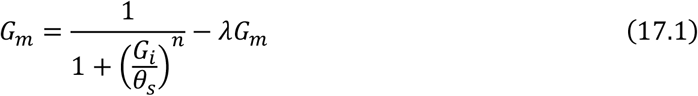

For *m* ≠ 1:

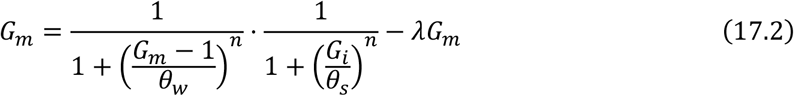

#### Oscillators

The basic logic of an oscillator, which is based on mutually repressive links and de-repressive interactions, is similar to that of a genetic cascade. However, in this case, the last gene in the sequence does not repress the first gene, allowing the system to cycle continuously (Eqs. 18.1, 18.2; Fig. 3I, I’; see Computer Simulation 18 (Text S2)).

For *m* = 1:

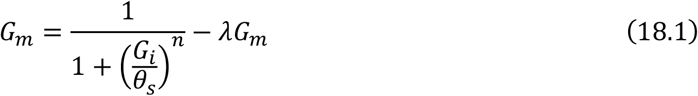

For *m* ≠ 1:

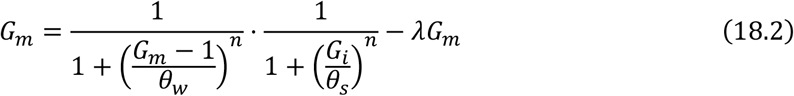

In our modeling framework of GRNs presented above, the cis-regulatory regions of constituent genes primarily encoded AND and/or OR regulatory logics of activators and repressors. However, developmental genes might exhibit more complex regulatory logic (24,46–48), with TFs acting as activators in some conditions and repressors in others (49–51). Given this complexity, one might question the value of the GRN models presented here—as well as many GRN implementations found in the literature— especially that their functionality often relies heavily on parameters that are experimentally challenging to determine. Some modeling approaches sidestep explicit GRN interactions altogether, focusing instead on the overall dynamic behavior of the network using simple equations to characterize overall functionality, like homeostasis or oscillations (52–54), an approach that is often called “geometric” modeling. While such approaches have their merits, GRN modeling remains valuable despite its unverified assumptions. Although the detailed wiring may not be entirely accurate, the overall regulatory logic of a GRN can capture essential system dynamics. For instance, as discussed, developmental genetic cascades can follow different overall regulatory logics—either a relay logic, where each gene activates the next, or a logic that depends of de-repressions, where down-regulation of one gene releases the next gene within the cascade. These two realizations may perform similarly under normal conditions but differ in robustness and responses to genetic perturbations (39). Pure dynamical modeling cannot capture these differences, whereas traditional GRN models, despite their limitations, offer insights into critical aspects of the network’s regulatory logic.

### 2.3 Cellular differentiation of GRNs: Space enters the scene

So far, we have discussed GRNs that function within a single cell. In development, although cells within a tissue share the same genes and GRNs (due to having the same genome), they are regulated differently, allowing each cell to express a unique set of TFs and follow distinct developmental trajectories. This manifests as gene expression patterns across the tissue, where different subsets of cells express different genes.

While such expression patterns could, in theory, take any form, they usually adopt specific, stereotyped patterns. Two major classes of patterns typically observed during early developmental stages are periodic and non-periodic patterns (Fig. 4A, A’; shown for simplicity as a one-dimensional row of cells) (1,42). Periodic patterns mediate the division of tissue into serial structures, such as demarcating the segmented structures along the anterior-posterior (AP) axis in arthropods and vertebrates. Non-periodic patterns, which are more common, mediate the division of tissue into distinct cell fates (regionalization). This differential expression of genes across a tissue results from regulating the same GRN differently in various cells (Figure 4B).

**Figure 4.**
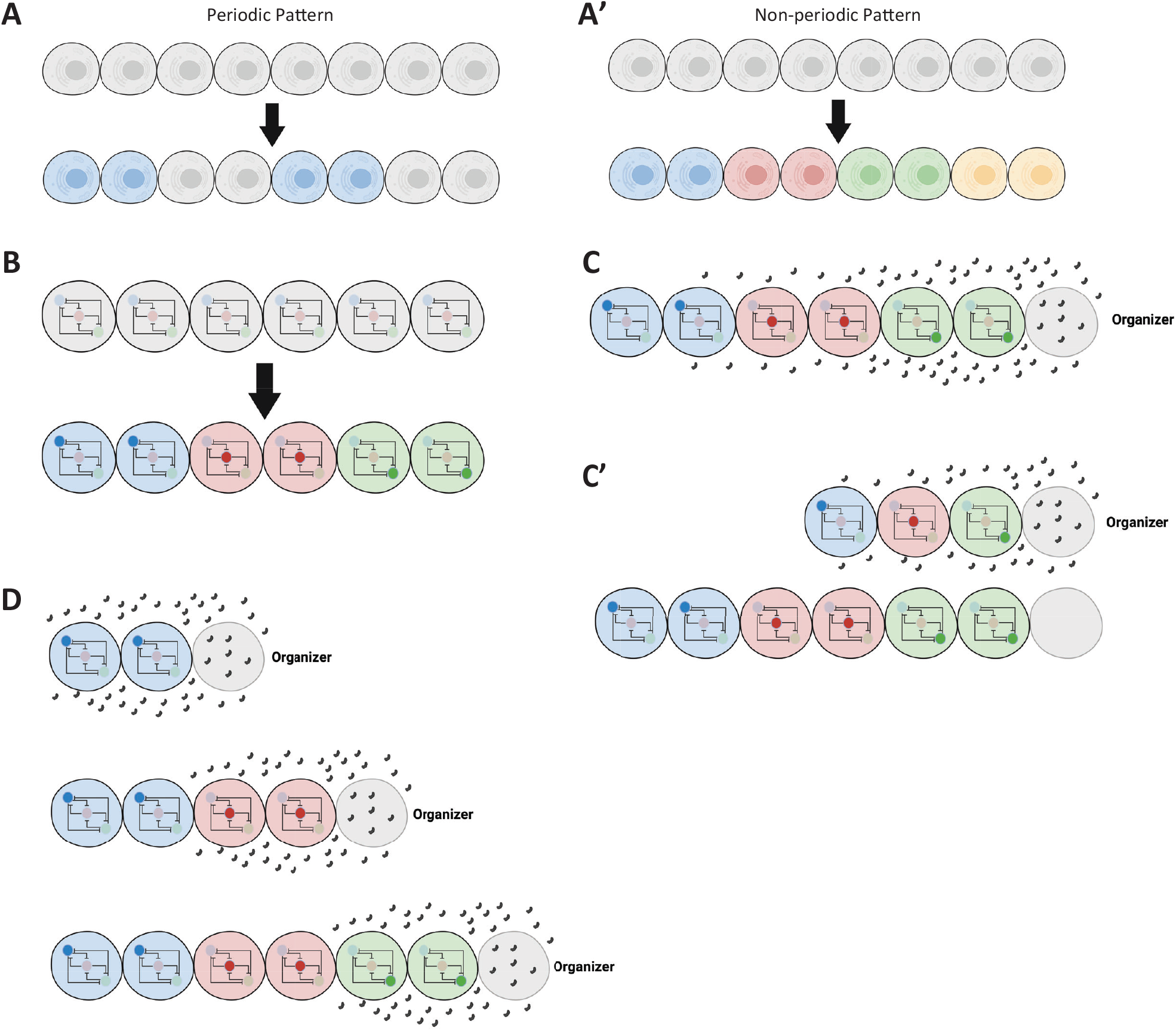
Introduction of spatial patterns in gene regulatory networks. **(A, A’)** Illustration of non-periodic and periodic patterns in a tissue. The schematic shows a linear arrangement of cells exhibiting non-periodic (distinct regions) and periodic (repeating units) gene expression patterns. **(B)** Conceptual representation of cells within a tissue sharing the same GRN but being regulated differently. Different regulatory inputs lead to distinct expression patterns despite identical GRNs in each cell. **(C)** Formation of a morphogen gradient by an organizer. An organizer at one end of the tissue secretes a ligand that diffuses to form a gradient, providing positional information to cells. **(C’)** Patterning followed by tissue elongation. The diagram illustrates how a tissue is patterned by a morphogen gradient before undergoing elongation, resulting in the expansion of patterned regions. **(D)** Concurrent tissue elongation and patterning with a retracting gradient. A moving morphogen gradient, exposing cells to changing morphogen concentrations over time, patterns the tissue as it elongates.

To regulate cells differently across a tissue, a morphogen gradient is often employed (9,55,56). An organizer (usually positioned at one extremity of the tissue to be patterned) secretes a ligand that diffuses and forms a morphogen gradient (Figure 4C). Different concentrations of the gradient set distinct developmental paths for cells along the tissue. However, tissue elongation—whether due to growth or convergent extension—often accompanies embryonic patterning. In some cases, tissue elongation occurs after patterning (Figure 4C’), while in others, it takes place concurrently with patterning.

Patterning a tissue during elongation is mechanistically distinct from patterning a non-elongating tissue (or a tissue patterned after elongation), as it often involves a dynamically retracting morphogen gradient. As a result, cells are exposed to changing concentrations of the morphogen over time (Figure 4D). In summary, embryonic patterning can be categorized based on the nature of the pattern (periodic vs. non-periodic) and whether the tissue is elongating or non-elongating (42). These developmental processes are explored further in the Results section.

The mechanisms that drive morphogen gradient formation range from passive diffusion to highly regulated ligand transport across a tissue, which are covered extensively in other reviews (55,57–60) and are beyond the scope of this study. Here, we use a simplified formulation for the gradient, independent of the mechanism that generates it. For the case of a non-retracting gradient in a non-elongating tissue, we use the following sigmoid function, where *x*_0_specifies the inflection point of the gradient, and *k* denotes the gradient slope:

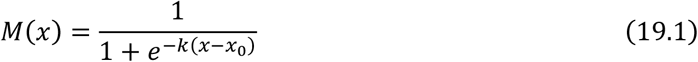

For the case of a retracting gradient, we employ the following moving sigmoid function, where *v* is the velocity of gradient retraction:

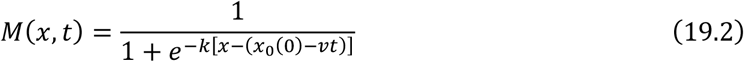

## 3 Results

In this section, we will discuss two of the most prominent models of embryonic pattern formation: the French Flag model (2,3) and the Temporal Patterning (or Speed Regulation) model (42,44,61–66). These models are phenomenological in nature, focusing on descriptive aspects and leaving out the molecular details of their genetic or molecular realization. We will explore four variations of each model, addressing the four common scenarios of embryonic patterning: (1) a non-periodic pattern in a non-elongating tissue, (2) a non-periodic pattern in an elongating tissue, (3) a periodic pattern in a non-elongating tissue, and (4) a periodic pattern in an elongating tissue.

Following this, we will apply the gene regulation models previously discussed to investigate different molecular and genetic realizations of these two phenomenological models of embryonic pattern formation. Although the reaction-diffusion model is another significant framework for understanding pattern formation, little is known about its regulation at the molecular level, and it will not be covered in this study.

### 3.1 The French Flag Model

The core mechanism of the French Flag model (2,3,42) is that different concentrations of a morphogen gradient activate different genes or cellular states (Fig. 5A). This mechanism is ideal for mediating non-periodic patterns in non-elongating tissue (Fig. 5A, left panel), where certain ranges of the morphogen gradient activate specific genes. However, a more challenging scenario arises when applying the French Flag model to pattern an elongating tissue during elongation. The model assumes a stable gradient to set precise boundaries between gene expression domains. To address this, a retracting gradient (M’; black in Fig. 5A, right panel; Eq. 20.1) activates a gene (M; grey in Fig. 5A, right panel; Eq. 20.2) with a slow decay rate (ideally *λ* = 0), forming a stable long-range morphogen gradient (grey) from a retracting short-range gradient (black). This stable gradient then mediates the division of the tissue into distinct gene expression domains. Extending the French Flag model to periodic patterns (Fig. 5B) is conceptually straightforward: alternating ranges of morphogen gradient concentrations turn a gene on and off. However, achieving this at the molecular level is more complex and will be discussed later.

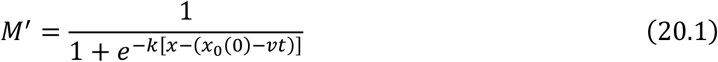

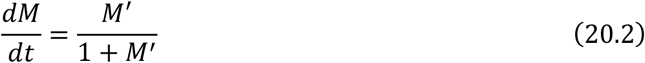

It is important to note that the French Flag model is a purely phenomenological, descriptive model and does not specify how it is realized at the molecular or genetic level. Here, we will explore potential molecular and GRN (gene regulatory network) realizations of this model. Despite the simplicity of the French Flag model, realizing it at the GRN level is challenging, as some genes must be both activated and repressed by the same morphogen gradient. For instance, in Fig. 5A, the red gene is activated by moderately high levels of the morphogen gradient (grey) but repressed at very high concentrations. This implies that the red gene responds to two different thresholds, T1 and T2, of the same morphogen gradient. None of the models discussed so far can accommodate this feature. While this phenomenon has been experimentally observed (50), it has not yet been fully mechanistically elucidated. For now, we will limit ourselves to simpler models where each gene responds to a single threshold of the morphogen.

We will start by reviewing potential GRN realizations of the French Flag model through a basic gene expression pattern where genes are expressed in bands extending to the tissue ends (Fig. 5C, C’). This simple pattern can be realized by adjusting the activation and repression thresholds of patterning genes. For example, in Fig. 5C, the blue gene (G1) is strongly activated by the morphogen gradient (M, grey in Fig. 5C, C’; see Computer Simulation 19 (Text S2)), extending its expression towards the lower end of the gradient. The red gene (G2), on the other hand, is weakly activated by M. This can be achieved by setting G1 to have a low activation threshold (low *K*_*G*1*M*_, indicating strong binding sites for M at G1’s CRM) and G2 to have a high activation threshold (high *K*_*G*2*M*_, indicating weaker binding sites for M at G2’s CRM). The expression domains of the green and orange genes (G3 and G4, respectively) are formed by having them repressed by M—strongly in the case of G3 (low *K*_*G*3*M*_) and weakly in the case of G4 (high *K*_*G*4*M*_) (Eq. 22).

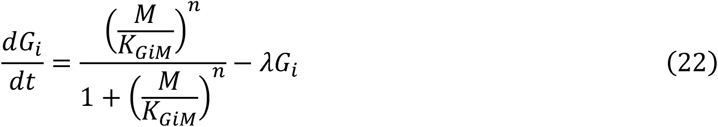

To create gene expression domains that do not extend to the boundaries of the morphogen gradient, two morphogen gradients can be employed. An example of this is shown in Fig. 5D, D’ (see Computer Simulation 20; (Text S2)), where two gradients (M1 and M2, represented by light and dark grey, respectively) together define the boundaries of the blue and red genes (G1 and G2). In this scenario, G1 is strongly repressed by M1 but weakly repressed by M2 (i.e., small *K*_*G*1*M*1_ and large *K*_*G*1*M*2_), while G2 is weakly repressed by M1 and strongly repressed by M2 (i.e., large *K*_*G*2*M*1_ and small *K*_*G*2*M*2_) (Eq. 23). This approach can be easily generalized to multiple genes by adjusting the repression thresholds of the two gradients for each gene.

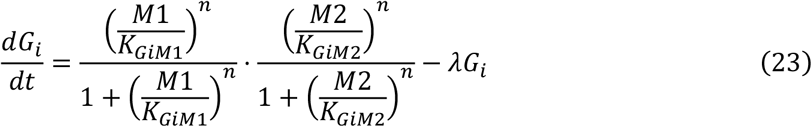

To generate multiple gene expression domains from a single morphogen gradient, cross-regulatory interactions between genes are required. The specific GRN wiring can vary, but one straightforward realization is shown in Fig. 5E. In this configuration, the morphogen gradient M activates patterning genes (G1, G2, and G3) at different thresholds. Without cross-regulatory interactions, this would result in broad expression domains, extending from the high concentration of M to the respective activation thresholds. Repression from other genes is needed to refine these domains. For instance, G1 is repressed by both G2 and G3, restricting its expression to the lower end of the gradient (see Computer Simulation 21 and its simulation in Fig. 5E’; Eq. 24.1-24.3).

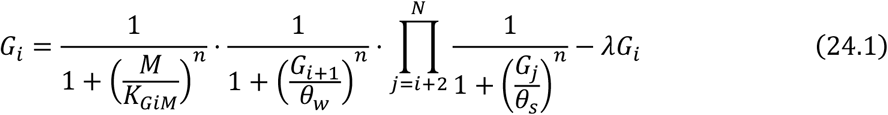

For *i* = *N* − 1:

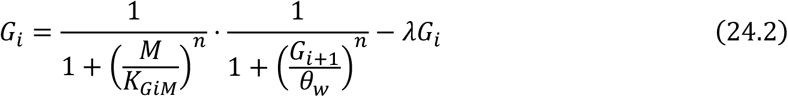

For *i* = *N*:

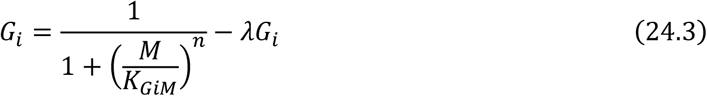

Other GRN configurations are also possible. For instance, in the AC-DC GRN motif (67–69), the morphogen gradient thresholds emerge from cross-regulation between genes, rather than being dictated by the gradient itself.

**Figure 5.**
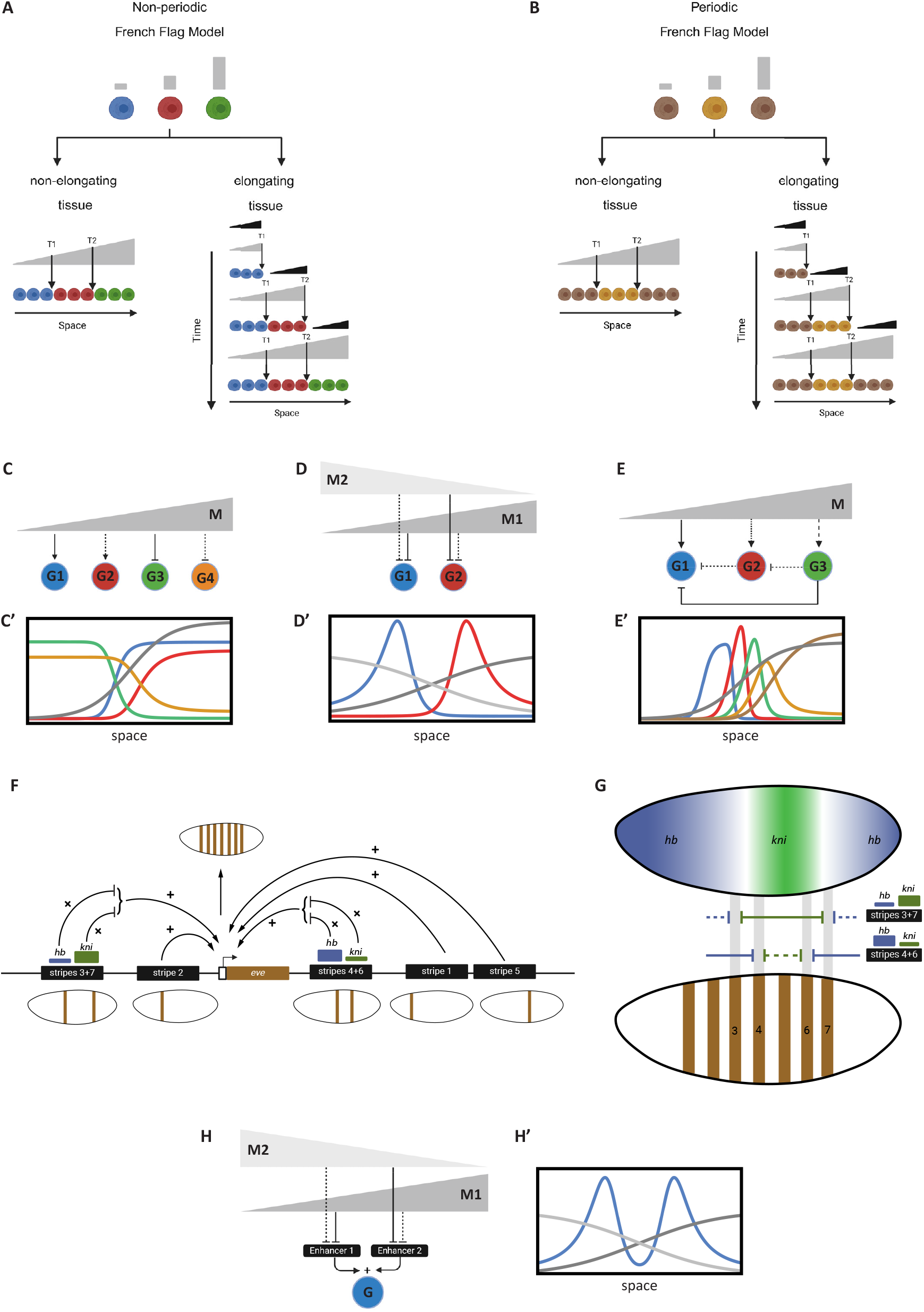
The French Flag Model and its gene regulatory network realizations. **(A)** *French Flag model for non-periodic patterning*. Left panel: A stable morphogen gradient divides the tissue into distinct gene expression domains (blue, red, green). Right panel: In an elongating tissue, a retracting gradient (black) induces a stable gradient with slow decay (gray) to pattern the tissue. **(B)** French Flag model applied to periodic patterning. Alternating concentrations of a morphogen gradient lead to the activation and repression of genes, creating repeating patterns across the tissue. **(C)** GRN realization using activation and repression thresholds. Genes G1-G4 are activated or repressed by different thresholds of the morphogen gradient M (gray), resulting in specific expression domains. **(C’)** Simulation of GRN realization shown in (C). This simulation corresponds to Computer Simulation 19 (Text S2). **(D)** Gene expression boundaries defined by two morphogen gradients. Gradients M1 (dark gray) and M2 (light gray) overlap to establish precise expression domains for genes G1 (blue) and G2 (red). **(D’)** Simulation of GRN realization shown in (D). This simulation corresponds to Computer Simulation 20 (Text S2). **(E)** GRN configuration involving cross-regulatory interactions. The morphogen gradient M activates genes G1, G2, and G3 at different thresholds, while cross-repressions between them refine their expression domains. **(E’)** Simulation of GRN realization shown in (E) This simulation corresponds to Computer Simulation 21 (Text S2). **(F)** The periodic expression of the gene eve in Drosophila is the result of the additive effect of multiple stripe-specific enhancers, each mediating one or two stripes. The logic of each enhancer is mediated by the multiplicative effect of several TF factors. Shown is the regulatory logic of enhancer 3+7 (mediating the expression of eve stripes 3 and 7), and enhancer 4+6 (mediating the expression of eve stripes 4 and 6). Both enhancers mediate the formation of stripes by integrating inputs from gap gene morphogens (mainly *hb* and *kni*). **(G)** A closer look at the regulatory logic of the eve 3+7 and 4+6 enhancers in *Drosophila* given the gene expression patterns of gap genes *hb* and *kni*. Harboring strong binding sites (thick line) for kni (green) but weak bind sites (thin line) for hb (blue), enhancer 3+7 place stripes 3 and 7 away from kni expression and close to abutting hb expressions. Harboring weak binding sites for kni but strong bind sites for hb, enhancer 4+6 place stripes 4 and 6 closer to central kni expression and away from abutting hb expressions. **(H)** *Additive effect of separate enhancers on gene expression*. Gene G is expressed in two domains, each regulated by a distinct enhancer (E1 and E2). The overall expression pattern is the sum of both enhancer activities. The same basic principle can be extended to form periodic pattern of arbitrary repetitions. **(H’)** Simulation of GRN realization shown in (H). This simulation corresponds to Computer Simulation 22 (Text S2).

So far, we have discussed the formation of non-periodic patterns, noting that their GRN realization is more challenging than the simplicity of the phenomenological French Flag model suggests. This difficulty primarily stems from the challenge of making a single gene respond to two different thresholds of the same morphogen gradient, which we addressed using either two morphogen gradients or cross-regulatory interactions. Applying the French Flag model to periodic patterns is even more complex, as genes must respond to multiple thresholds, with the morphogen gradient alternately activating and repressing the same gene. One solution is similar to the non-periodic case: decomposing the periodic pattern into several non-periodic patterns, each mediated separately by two morphogen gradients. The final periodic pattern is the sum of all sub-patterns.

This method of generating periodic patterns has been extensively studied in the context of AP patterning in the early *Drosophila* embryo (Fig. 5F, G), providing insights into the organization of cis-regulatory modules during development (70). A notable example is the expression of the *even-skipped* (eve) gene, which forms seven distinct stripes in the early *Drosophila* embryo (70,71). Rather than arising from a periodic process like an oscillator or reaction-diffusion mechanism (6,7), this pattern results from decomposing the periodic pattern into several non-periodic sub-patterns, each composed of a single stripe or pair of stripes (41,42,72,73). Each sub-pattern is mediated by a dedicated enhancer, and the final expression of *eve* is the sum of all enhancer activities. The regulatory logic of each enhancer is primarily determined by the multiplicative (AND) interactions of repressing TFs known as gap genes, which form (localized) morphogen gradients in the early *Drosophila* embryo. Fig. 5F, G illustrates the regulatory logic of two enhancers (74): the 3+7 enhancer (which mediates the formation of the 3rd and 7th stripes) and the 4+6 enhancer (which mediates the formation of the 4th and 6th stripes). The positioning of these stripes is determined by the binding strengths of the gap gene morphogens *hunchback* (*hb*) and *knirps* (*kni*) to the 3+7 and 4+6 enhancers along the AP axis.

This example, along with others, demonstrates that an additive OR logic is mediated by using separate enhancers spaced far enough to avoid interference between their activities, while multiplicative AND logic is mediated by placing TF binding sites close to each other within the same enhancer.

A simple example of modeling this mechanism is shown in Fig. 5H, H’, where gene G is expressed in two domains, each mediated by the additive effects of two separate enhancers: Enhancer 1 (E1) and Enhancer 2 (E2). E1 is strongly repressed by morphogen 1 (M1) and weakly repressed by morphogen 2 (M2), while E2 is weakly repressed by M1 and strongly repressed by M2 (see Computer Simulation 22 and its simulation in Fig. 5H’; Eq. 25.1-25.2).

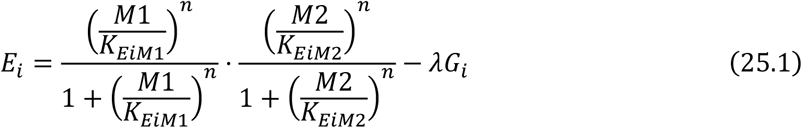

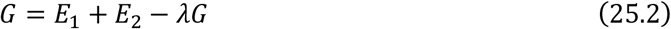

In the GRN realization of the French Flag model presented above, we considered only the case of non-elongating tissues. The extension to the case to elongating tissues, where both M and M’ gradients are used (right panels in Fig. 5A and 5B) is straight forward (61).

### 3.2 Temporal Patterning or the Speed Regulation Model

In temporal patterning, a temporal sequence is translated into a spatial pattern rather than directly mediating a spatial pattern (42,75,76). This mode of patterning, although less intuitive, is a standard mechanism in the development of neurogenesis (77–79) in many animals and at various stages of development. Recent studies have shown it is involved in many other tissues as well, such as segmentation and regionalization of the AP axis in short-germ insects (63,80) and vertebrates (76,81,82), ventral neural tube patterning (66), and limb bud development (83,84). This raises the possibility that temporal patterning may be the default mechanism for embryonic pattern formation. While the reason for this preference is not fully understood, it has been suggested that this mechanism may offer greater robustness compared to alternatives, such as the French Flag model (53,64,67). It has also been hypothesized that since animals evolved from single-celled organisms, many evolutionarily conserved gene regulatory networks (GRNs) are temporal in nature, and multicellularity may have evolved by translating these conserved temporal patterns into spatial ones (79).

The Speed Regulation (SR) model (44) was recently proposed as a unifying mechanism for various types of temporal patterning observed in different tissues. The SR model synthesizes several patterning schemes that all share the core feature of a morphogen gradient modulating the speed of a temporal sequence, whether periodic or non-periodic. In the non-periodic version of the SR model, each cell within a tissue progresses through successive states (depicted in different colors in Fig. 6A), with each state defined by the expression of one or more genes. The speed of transitions between these states is controlled by a molecular factor (referred to as the “speed regulator,” shown in grey at the top of Fig. 6A). At very low or zero concentrations of the speed regulator, the transitions slow down to the point where the states are indefinitely stabilized. When a group of cells experiences a gradient of the speed regulator (Fig. 6A’; left panel: ‘for non-elongating tissues’), all cells transition through successive states, but at progressively slower speeds as they encounter lower concentrations of the gradient. This creates the illusion of cellular states propagating as waves from high to low gradient concentrations. These waves, known as “kinematic” or “pseudo-waves,” do not rely on diffusion or cell-to-cell communication (85,86). We refer to this version of the model as “gradient-based speed regulation,” which is particularly suited for patterning non-elongating tissues (Fig. 6A’, left). The model can also pattern elongating tissues if the gradient retracts as a wavefront, a process we call “wavefront-based speed regulation” (Fig. 6A’, right). The SR model can also generate periodic structures if the sequential gene activation process is driven by a biological clock instead of a sequential transition of states (42). Notably, in the wavefront-based version of SR, if the wavefront is in the form of a tapered gradient (a superposition of the gradient-based and wavefront-based SR models), kinematic waves will propagate from high to low concentrations of the gradient, in the opposite direction of wavefront retraction, as the tissue elongates.

**Figure 6.**
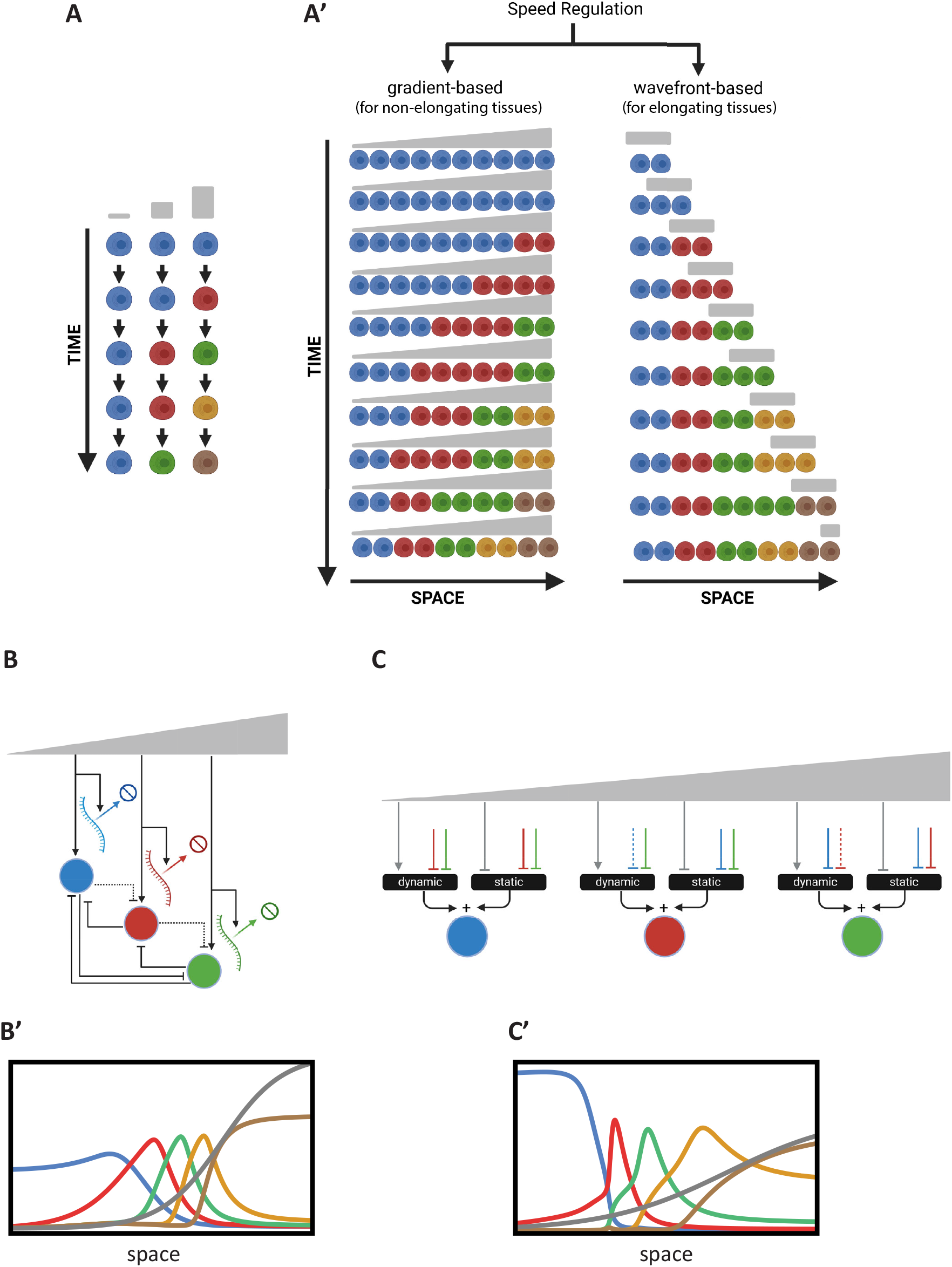
Temporal Patterning or the Speed Regulation Model. **(A)** Schematic of the temporal sequence of cellular states in temporal patterning. Cells progress through successive states (depicted in different colors), with transitions regulated by a speed regulator (gray). **(A’)** Gradient-based and wavefront-based speed regulation in tissue patterning. Left: In a non-elongating tissue, a morphogen gradient modulates the speed of transitions between cellular states, resulting in the induction of non-periodic waves that eventually stabilize into a non-periodic pattern. Right: In an elongating tissue, a retracting wavefront regulates the speed of state transitions, resulting in the formation of a non-periodic pattern. **(B)** Realization of the Speed Regulation model by modulating transcription and decay rates. The morphogen gradient jointly affects the overall transcription and mRNA degradation rates to control the timing of gene expression. **(B’)** Simulation of the GRN realization in (B). This simulation corresponds to Computer Simulation 23 (Text S2). **(C)** Enhancer switching model as a GRN realization of the Speed Regulation model. Each patterning gene is regulated by the additive sum of a dynamic GRN (driving sequential activities) and a static GRN (stabilizing expression) through two separate enhancers. The speed regulator modulates the balance between these GRNs. **(C’)** Simulation of the GRN realization in (C). This simulation corresponds to Computer Simulation 24 (Text S2).

Like the French Flag (FF) model, the SR model is phenomenological and does not propose a molecular mechanism or GRN realization that can transform temporal sequences (or oscillations) into patterns. A straightforward realization of the model is for a morphogen gradient (the speed regulator) to modulate the overall transcription rate, including mRNA decay rates (61) (Fig. 6B) M2 (see Computer Simulation 23 (Text S2) and its simulation in Fig. 6B’).

For *j* = 1:

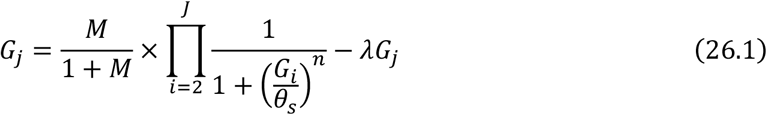

For *j* ≠ 1:

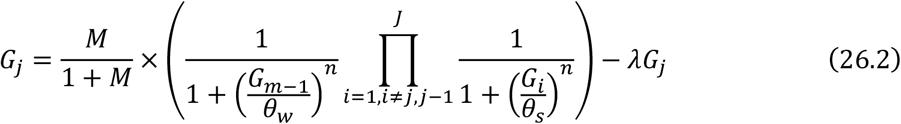

While this approach works well, it requires that the decay rates of all genes involved in patterning be regulated by the morphogen gradient in proportion to the regulation of their transcription. Achieving this at the molecular level may be unfeasible.

An alternative mechanism, recently proposed to modulate the timing of GRNs, is the Enhancer Switching Model (42,44,87). This model posits that each patterning gene is simultaneously wired into two GRNs (Fig. 1C): (i) a dynamic GRN that drives periodic or sequential gene activities, and (ii) a static GRN that stabilizes gene expression patterns. The concentration of the speed regulator (shown in grey in Fig. 1C) activates the dynamic GRN while repressing the static GRN, thus balancing the contribution of each GRN to the overall dynamics and consequently regulating the speed of gene expression. At high concentrations of the speed regulator, the dynamic GRN is dominant, facilitating rapid oscillations or sequential gene activities. At low concentrations, the static GRN dominates, resulting in slower oscillations or sequential gene activities. Each gene is connected to these two GRNs through two enhancers: (i) a dynamic enhancer that encodes the wiring of the gene within the dynamic GRN, and (ii) a static enhancer that encodes the wiring of the gene within the static GRN (Fig. 1D) (Eqs. 27.1-27.4; see Computer Simulation 24 (Text S2) and its simulation in Fig. 6C’).

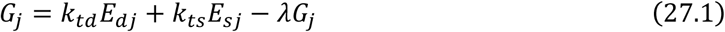

For the dynamic enhancer, *E*_*dj*_ :

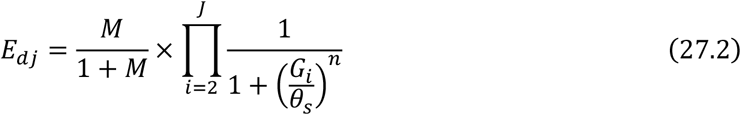

For the static enhancer, *E*_*sj*_ :

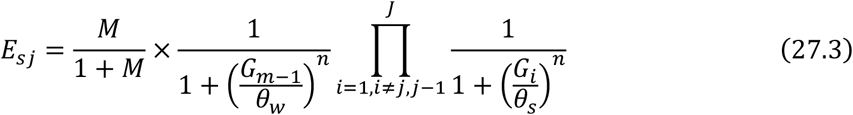

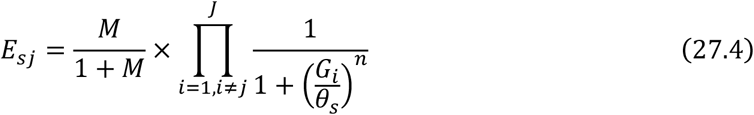

The Enhancer Switching model has been proposed as the molecular mechanism underlying AP patterning in the intermediate-germ insect *Tribolium castaneum*, with partial experimental support (61,87). This model also aligns with recent findings on AP patterning in *Drosophila* (88–91). However, it requires further rigorous validation to confirm its applicability.

### 3.3 Case study: Impact of enhancer integration mechanisms on embryonic patterning

So far, we have discussed various embryonic pattern formation mechanisms using basic models of transcription. To that end, we employed simple toy models at each level of the modeling process. One of the main advantages of this approach is that it allows us to examine the effect of fundamental transcriptional and gene regulatory mechanisms on the overall performance of embryonic patterning systems. In this section, we will explore a specific example to demonstrate this. We will examine how the integration of multiple enhancer activities influences the performance of the enhancer switching model.

In our modeling of the enhancer switching mechanism (Fig. 6C, and Eqs. 27.1–27.4), we used a weighted sum of the two enhancer activities (Eq. 27.1). The physical interpretation of this relationship is that each enhancer can transcribe the gene independently (Fig. 7A). In other models of enhancer function, however, enhancers compete for binding to the promoter, and the enhancer that wins the competition sets the transcriptional state of the promoter (Eq. 28; Fig. 7B) (92). In the first scenario (Fig. 7A; Eq. 27.1), each enhancer determines its own final transcriptional rate (denoted as *k*_*td*_ and *k*_*ts*_ in Eq. 27.1), whereas in the second scenario, the normalized activities of competing enhancers set the transcriptional state of the promoter, but the maximum final transcriptional rate (denoted as *k*_*t*_ in Eq. 28) is set by the promoter itself (Fig. 7B).

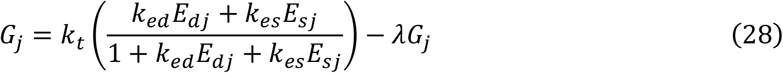

The normalization effect of the enhancer competition scenario may be a crucial mechanism for integrating enhancer activities. The overall transcription level of a gene is a key parameter in any GRN, as the activation thresholds of genes within a GRN depend on the average transcription rates of the genes regulating them. Enhancers are rapidly evolving genetic elements and are the primary drivers of novelty in animal evolution. However, this also introduces a risk, as allowing a newly evolved enhancer to take control of a key transcriptional parameter, such as the overall average transcription rate, could disrupt normal function. Alternatively, in the enhancer competition model, enhancers first compete to activate the promoter, with the promoter acting as the final arbiter for the transcription rate.

**Figure 7.**
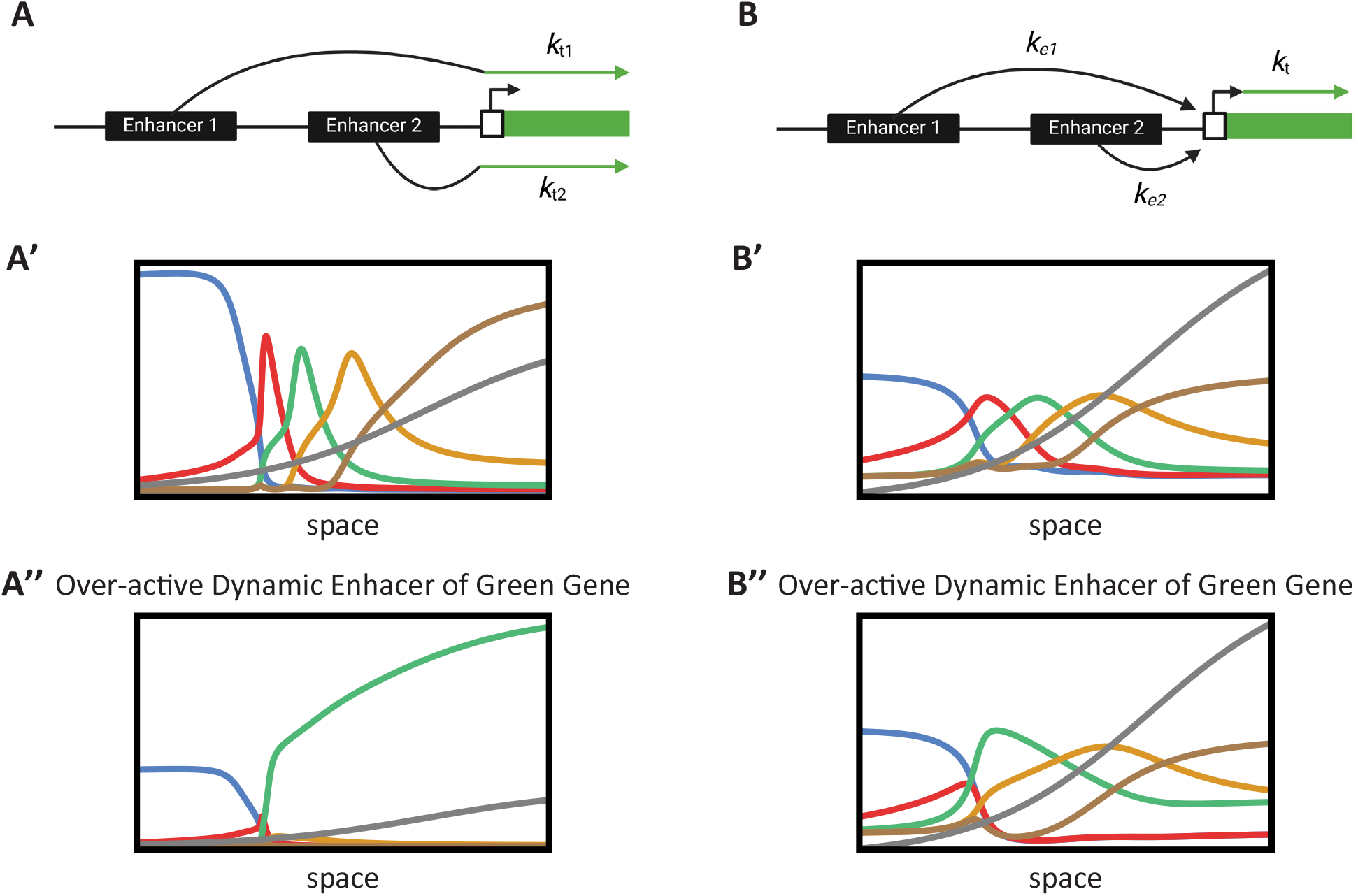
Case study: Impact of enhancer integration mechanisms on embryonic patterning. **(A)** *Additive enhancer integration model*. Multiple enhancers independently contribute to the transcription rate of a gene. The total transcription is the weighted sum of the individual enhancer activities. **(A’)** Simulation of gene expression with additive enhancer integration under normal conditions. The performance of the enhancer switching model is shown, demonstrating effective temporal patterning. This simulation corresponds to Computer Simulation 25 (Text S2) with mut_flag = 0 (wild-type). **(A”)** Impact of an overactive enhancer in the additive model. The simulation illustrates how an overactive enhancer disrupts gene expression, leading to aberrant patterning and loss of GRN function. Impact of an overactive enhancer in the additive model. The simulation illustrates how an overactive enhancer disrupts gene expression, leading to aberrant patterning and loss of GRN function. This simulation corresponds to Computer Simulation 25 (Text S2) with mut_flag = 1 (overactive dynamic enhancer of Gene 3). **(B)** Enhancer competition integration model. Enhancers compete for binding to the promoter of a gene. The promoter’s transcriptional state is determined by the enhancer that successfully binds, normalizing the transcriptional output. **(B’)** Simulation of gene expression with enhancer competition integration under normal conditions. The model shows comparable performance to the additive case, maintaining proper temporal patterning. This simulation corresponds to Computer Simulation 26 (Text S2) with mut_flag = 0 (wild-type). **(B”)** Mitigation of overactive enhancer effects in the competition model. The simulation demonstrates that the competitive mechanism normalizes transcription levels, preserving GRN function despite the presence of an overactive enhancer. This simulation corresponds to Computer Simulation 26 (Text S2) with mut_flag = 1 (overactive dynamic enhancer of Gene 3).

To investigate this further, we simulated our enhancer switching model for both the additive case (Fig. 7A, A’; see Computer Simulation 25 (Text S2)) and the enhancer competition case (Fig. 7B, B’; see Computer Simulation 26 (Text S2)). Both scenarios resulted in comparable performance (Fig. 7A’ and 7B’). We then examined the effect of an overactive enhancer (Fig. 7A” and 7B”). In the additive case, the overactive enhancer disrupted the overall function of the GRN (Fig. 7A”), whereas in the enhancer competition case, the normalization effect mitigated the impact of the overactive enhancer, preserving the overall performance of the GRN (Fig. 7B”).

## 4 Discussion

In this paper, we developed a comprehensive framework to model the emergence of embryonic patterning, linking molecular gene regulation to tissue-level organization. We began by modeling transcription at the single-gene level using basic chemical reaction models and extended this to model GRNs that govern specific cellular functions. We then introduced phenomenological models of embryonic pattern formation, such as the French Flag model and Speed Regulation models, integrating these with molecular and GRN realizations. To facilitate understanding and application of our models, we accompanied our mathematical framework with computer simulations, providing intuitive and simple code for each model. Through our case study on enhancer integration, we demonstrated that a two-step gene regulation strategy—enhancer activation followed by competitive integration at the promoter—ensures robustness and evolvability in gene expression patterns, emphasizing the adaptability of eukaryotic transcriptional regulation.

Modeling gene regulation during embryonic development has traditionally followed two distinct strategies. The first involves using finely tuned GRN models to fit large datasets, which we refer to as the “simulation approach” (45,93,94). The second relies on simplified, abstract models that capture the overall behavior of the observed data, which we call the “toy model approach” (15,44,67,95,96). In our view, the simulation approach is only appropriate when detailed biochemical data are available, which is often not the case. By contrast, while the toy model approach is inherently oversimplified, it offers the advantage of conceptual clarity. However, both approaches share a significant limitation in modeling gene regulation during development: they rely on gene regulatory models that are largely inspired by bacterial systems. These models may not accurately reflect the complexity of eukaryotic transcriptional machinery, particularly in animals. In this paper, we adopted the toy model approach, acknowledging its inherent limitations. However, our aim was to explicitly articulate the various assumptions made when modeling gene regulation, from the molecular scale to the tissue level. This transparency allows us to critically examine and challenge these assumptions throughout our modeling process. The simplicity and low computational cost of the toy model approach—unlike the simulation approach, which requires extensive data collection and optimization—facilitates this exploration. Moreover, we provided accompanying computer simulations with intuitive and simple code, making our models accessible and facilitating their use as educational tools or starting points for further research. In fact, we present our model not as a definitive framework, but as a basis for questioning and refining the assumptions at each step. Our hope is that future work by our group and others will further challenge and improve upon these ideas.

One specific aspect of gene regulation we examine in this paper, which relates to the complex transcriptional machinery of animals, is the impact of multiple enhancer integration on the performance of GRNs during development—an aspect largely overlooked in previous modeling efforts. This represents a small departure from the bacteria-inspired models traditionally used in gene regulation studies, moving instead toward the more intricate transcriptional mechanisms found in animals, particularly during development. However, this is neither the only nor the most critical aspect to reconsider in efforts to model gene regulation more realistically. A widely used assumption, which we also adopted in our modeling, is the simplistic regulatory logic of individual genes within the GRN, typically modeled with basic OR and/or AND relationships. Under this assumption, the complexity of GRN performance stems from the network’s structure rather than from the computational capabilities of individual genes. Yet, it has been observed that single genes can exhibit far more complex regulatory logic (48). For instance, a single TF may act as an activator or a repressor, depending on the concentration of co-regulating TFs (49). These intricate regulatory interactions are likely evolutionarily conserved and could serve as modular building blocks, akin to Lego pieces, which can be used as is or slightly modified to construct more extensive GRNs.

Another key aspect of gene regulation that warrants further investigation using our modeling framework is how the timing of GRNs is modulated. Recent findings suggest that many phenomena in embryonic patterning arise from the modulation of GRN timing by morphogen gradients, supporting a temporal/speed regulation model. In this paper, we modeled this effect using two distinct hypotheses (shown in Fig. 6B and 6C, respectively). However, given the largely unexplored computational complexity of transcriptional machinery in developmental genes, it is conceivable that additional mechanisms for timing regulation exist. Moreover, recent experimental findings suggest other assumptions in current models of gene regulation may need re-evaluation. For example, the effects of transcriptional bursting (19,97) and the formation of TF clusters or condensates (98–100) on GRN performance—particularly in embryonic patterning—remain unclear. These newly discovered phenomena could significantly influence how GRNs function.

With so many uncertainties surrounding current gene regulation and GRN modeling approaches, how can we approach the complex task of modeling embryonic patterning? While the answer is not straightforward, careful and minimal use of simple, transparent models—despite their inaccuracies— can yield partial insights. This is achievable as long as we remain aware of the models’ assumptions and are willing to critically evaluate them. In this paper, our goal has been to accomplish exactly that. To support further exploration, we offered a simple and transparent computational framework for modeling gene regulation during embryonic pattern formation, complete with accessible simulations and intuitive code. This enables researchers to test and expand upon our models, promoting a deeper understanding of the mechanisms underlying gene regulation in development.

## Figure Legends

**Figure S1.**
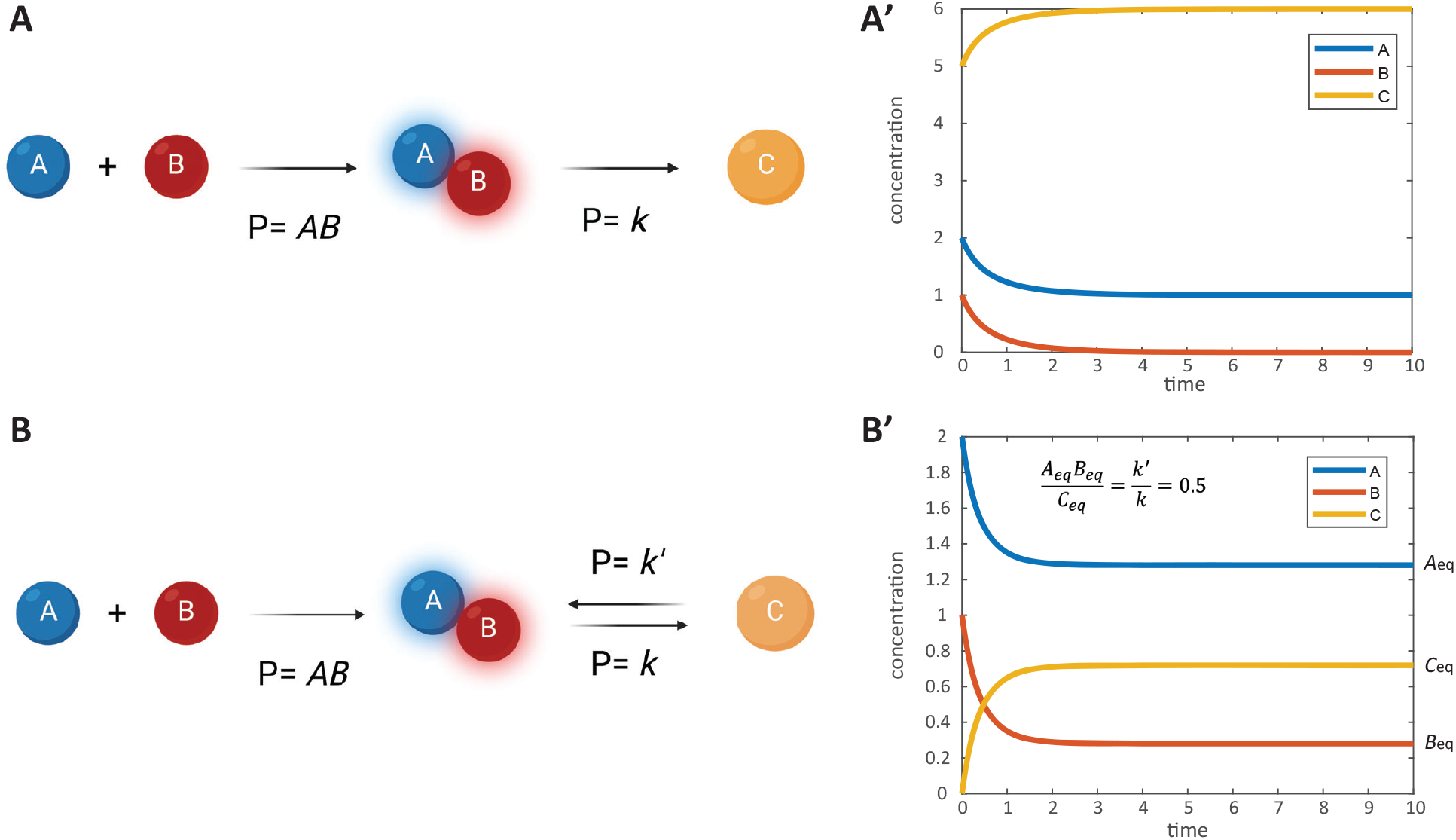
Modeling the kinetics of simple chemical reactions. **(A)** Simulation of the kinetics of a simple chemical reaction where chemicals A and B react to form chemical C. The reaction follows the equation *A* + *B* → *C*, with a reaction rate proportional to the concentrations of A and B. The graph shows the concentrations of A (blue line), B (orange line), and C (green line) over time. As the reaction progresses, the concentrations of reactants A and B decrease while the concentration of product C increases, illustrating the consumption of reactants and formation of product over time. This simulation corresponds to Computer Simulation 1 (Text S1). **(B)** Simulation of a reversible chemical reaction where chemicals A and B react to form chemical C, which can also decompose back into A and B. The reaction follows the equation *A* + *B* ↔ *C*, incorporating both forward and reverse reactions with rate constants *k* and *k*′, respectively. The graph displays the concentrations of A (blue line), B (orange line), and C (green line) over time. The system approaches equilibrium where the rates of the forward and reverse reactions balance, resulting in constant concentrations of A, B, and C. This simulation demonstrates how reversible reactions reach equilibrium within a given timeframe and corresponds to Computer Simulation 2 (Text S2).

**Figure S2.**
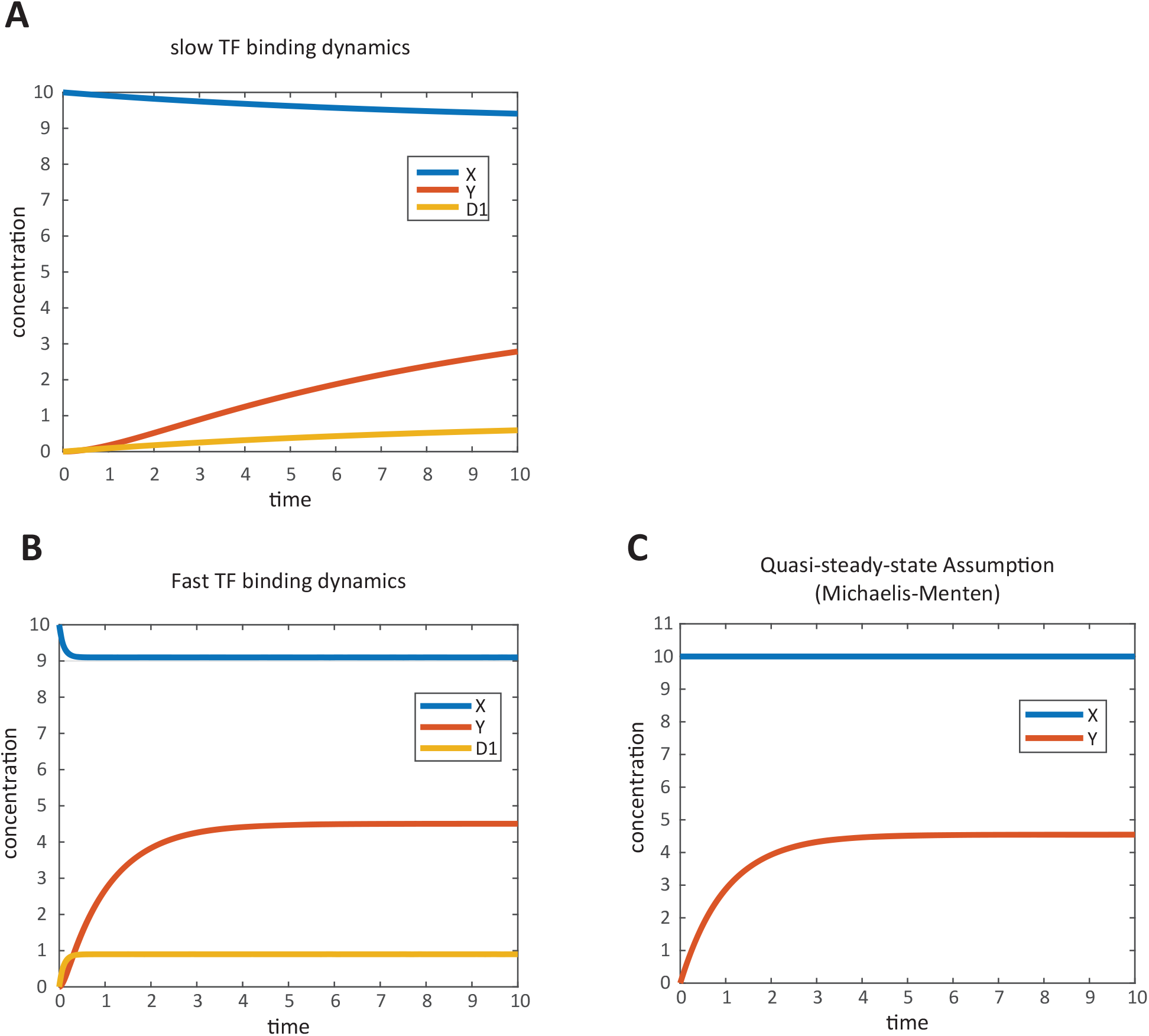
Modeling gene activation dynamics and the effects of transcription factor binding kinetics. **(A)** Simulation of gene activation by a single activator transcription factor (TF) with fast binding dynamics (high *k*_1_ and *k*_−1_). The model illustrates the interaction between TF X, the active DNA state D1, and the mRNA transcript Y. The graph shows the concentrations of X (blue line), D1 (green line), and Y (orange line) over time. Rapid TF binding and unbinding lead to quick fluctuations in the active DNA state and a prompt increase in mRNA production, resulting in a swift response in gene expression levels. This highlights how fast TF-DNA interactions can affect the timing of gene activation. This simulation corresponds to Computer Simulation 3 (Text S2) with high *k*_1_ and *k*_−1_ values. **(B)** Simulation of gene activation by a single activator TF with slow binding dynamics (low *k*_1_ and *k*_−1_). The graph depicts the concentrations of X (blue line), D1 (green line), and Y (orange line) over time. Slower TF binding and unbinding result in delayed formation of the active DNA state and a gradual increase in mRNA levels. This demonstrates that slow TF-DNA binding kinetics can lead to a delayed gene expression response, affecting the timing of downstream cellular processes. This simulation corresponds to Computer Simulation 3 (Text S2) with low *k*_1_ and *k*_−1_ values. **(C)** Simulation of gene activation using Michaelis-Menten kinetics. The model describes how the concentration of mRNA transcript Y changes over time in response to a constant concentration of activator TF X. The graph shows the concentration of Y (orange line) increasing over time and reaching a steady state. The use of Michaelis-Menten kinetics captures the saturation effect where, at high TF concentrations, the rate of mRNA production approaches a maximum due to the limited number of available binding sites on the DNA. This results in a hyperbolic relationship between TF concentration and gene expression level, illustrating the principles of enzyme kinetics applied to transcriptional activation. This simulation corresponds to Computer Simulation 4 (Text S2).

## Text S1

### Modeling chemical reactions

The Law of Mass Action describes the relationship between the concentrations of reactants and the reaction rate in a chemical reaction. According to this law, the reaction rate is directly proportional to the product of the reactants’ concentrations. In this context, the concentration of a species can also be understood as the probability of a molecule encountering another molecule with which it can react.

In the example shown in Fig. S1-A, chemical species A interacts with chemical species B to produce a new species C, with a rate constant *k*. This constant can be interpreted as the probability that, upon collision, species A and B will successfully transform into species C. This relationship is represented by the following chemical equation.

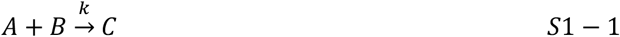

According to the law of mass action, the rate at which this forward reaction occurs can be described by *kAB*, where *k* is the rate constant, and A and B represent the concentrations of the reactants. The rates of change of the concentrations of the reactants and the product can be described by the following system of differential equations:

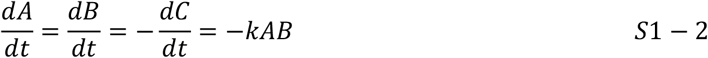

As the reaction proceeds, the concentrations of A and B decrease as they are consumed to form the product C, as shown in Computer Simulation 1 (Text S2) and Fig. S1-A’.

#### Equilibrium

Here we will consider a reversible version of the reaction considered in the example above:

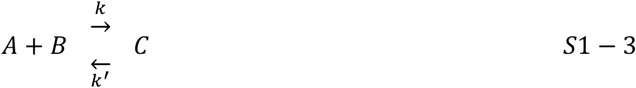

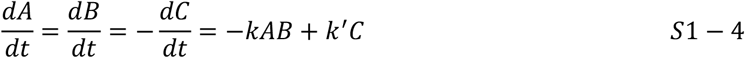

In a reversible reaction, the system will eventually reach equilibrium, where the rates of the forward and reverse reactions are equal. At equilibrium, the concentrations of the reactants and product remain constant because the forward reaction rate equals the reverse reaction rate:

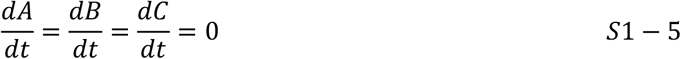

Substituting this into Eq. S1-2, the relative relationship between reactants at equilibrium can be given by Eq. S1-6, as shown in by Computer Simulation 2 (Text S2) and Fig. S1-B’.

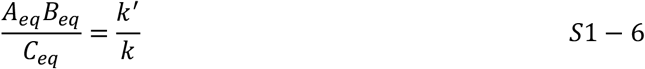

## Text S2

**Computer Simulation 1**. Modeling the Kinetics of a Simple Chemical Reaction Between A and B to Form C.

The results of this computer simulation are shown in Figure S1-A’.

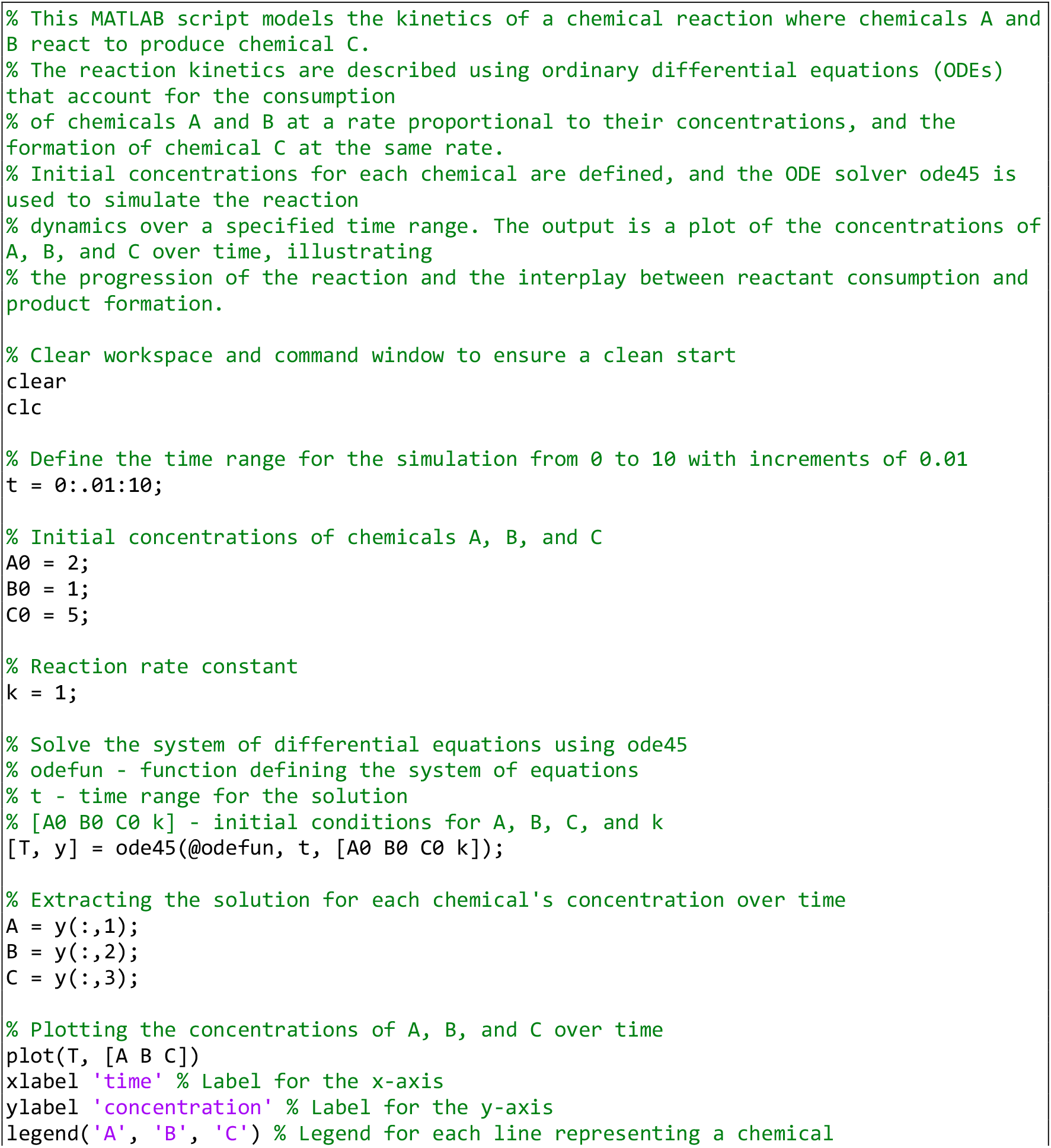
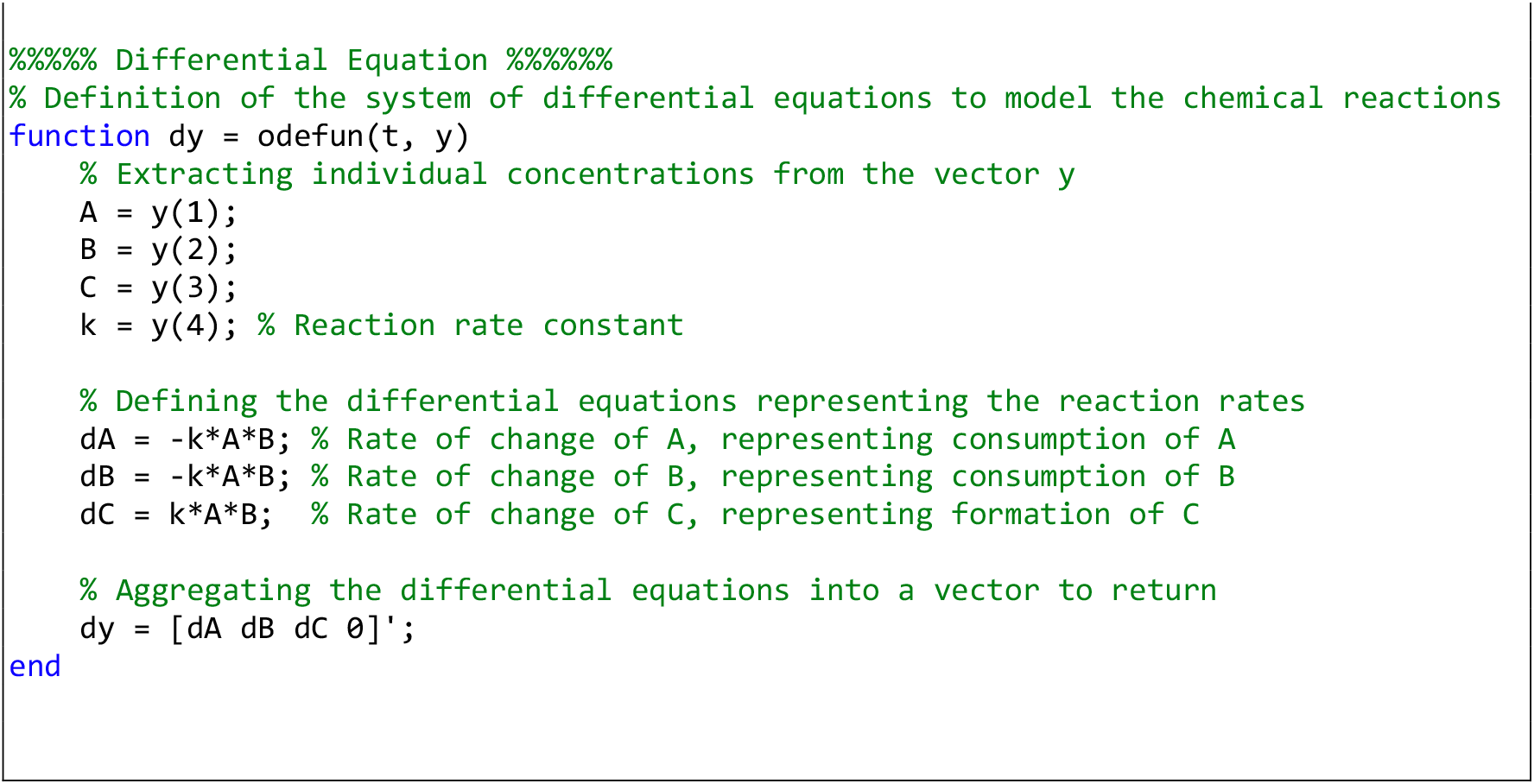

**Computer Simulation 2**. Modeling Reversible Chemical Reaction Dynamics Towards Equilibrium.

The results of this computer simulation are shown in Figure S1-B’.

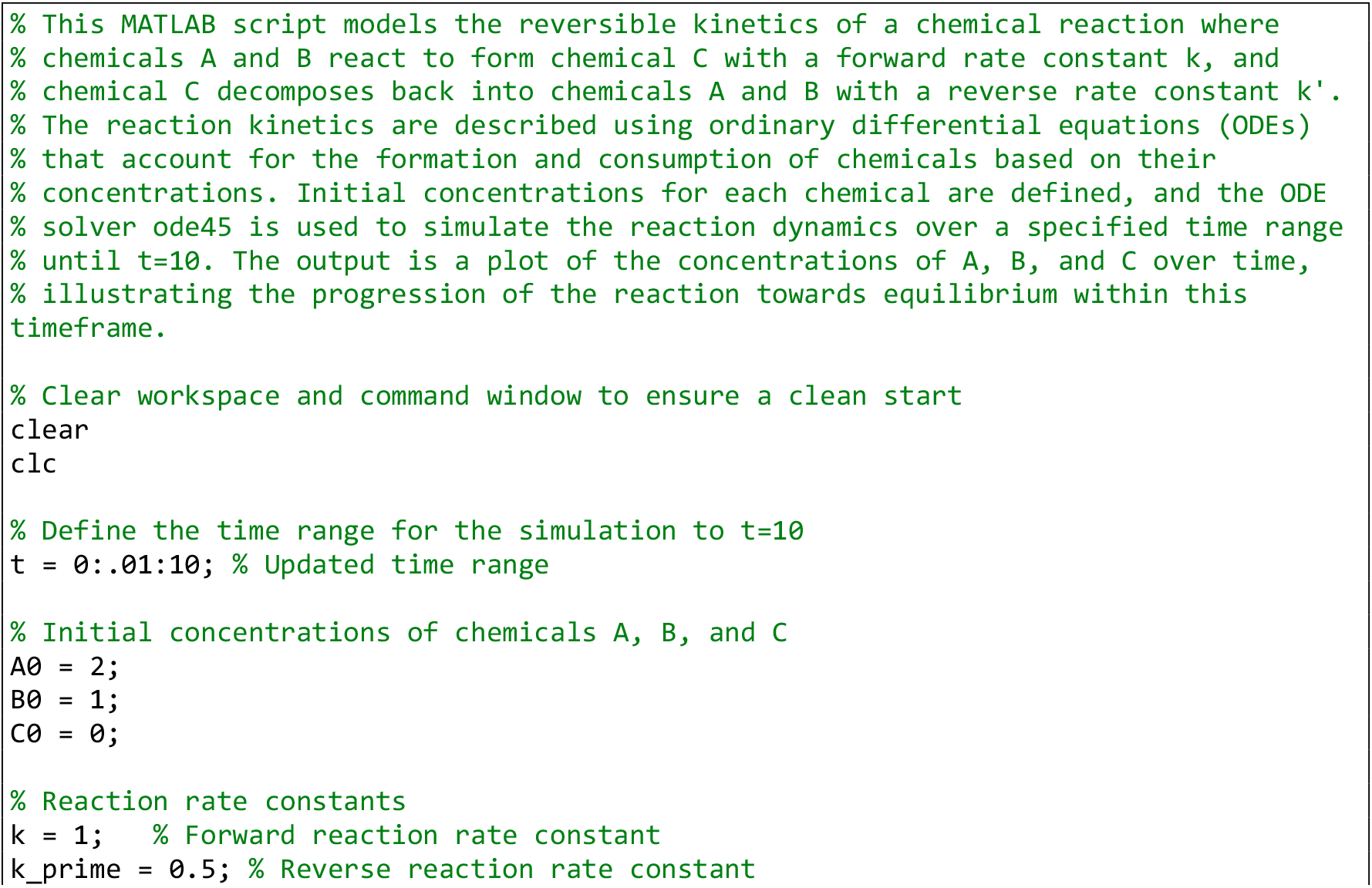
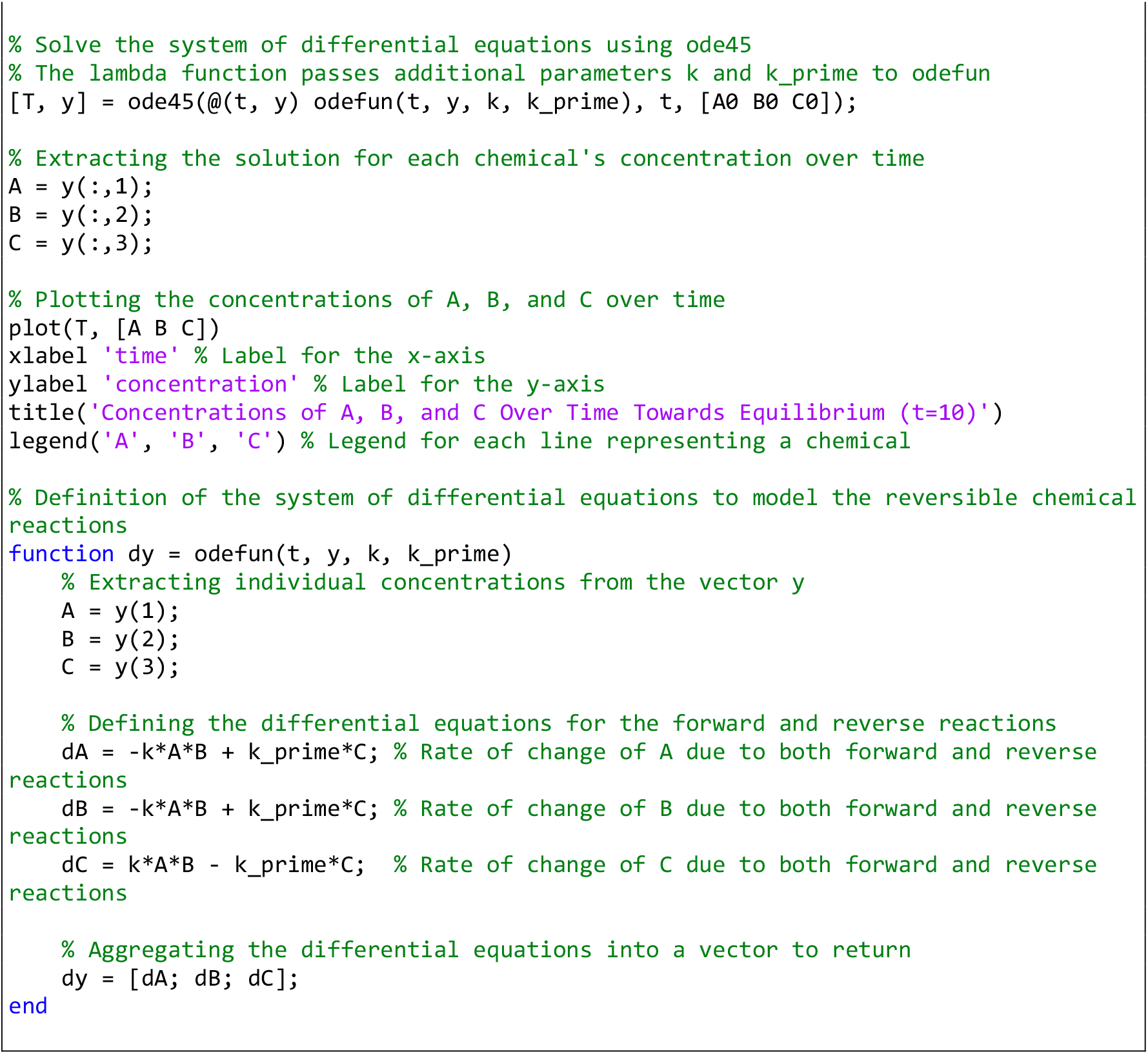

**Computer Simulation 3**. Modeling Gene Activation by a Single Activator with Varying Transcription Factor Binding Dynamics.

The results of this computer simulation are shown in Fig. S2-A (fast TF binding dynamics; high k1 and k_1), and Fig. S2-B (slow TF binding dynamics; low k1 and k_1). Recommended values for binding constants for slow dynamics are k1=k_1=1. Recommended values for binding constants for fast dynamics are k1=k_1=.01.

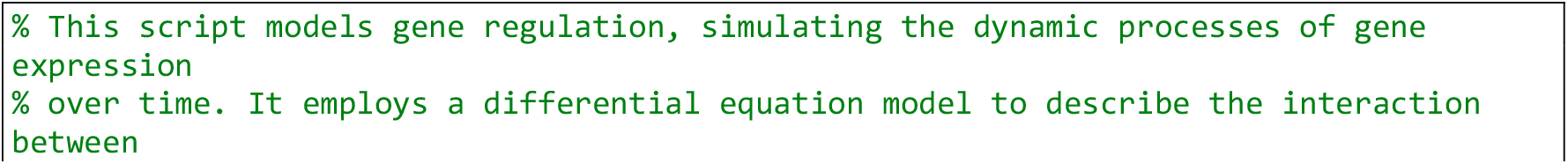
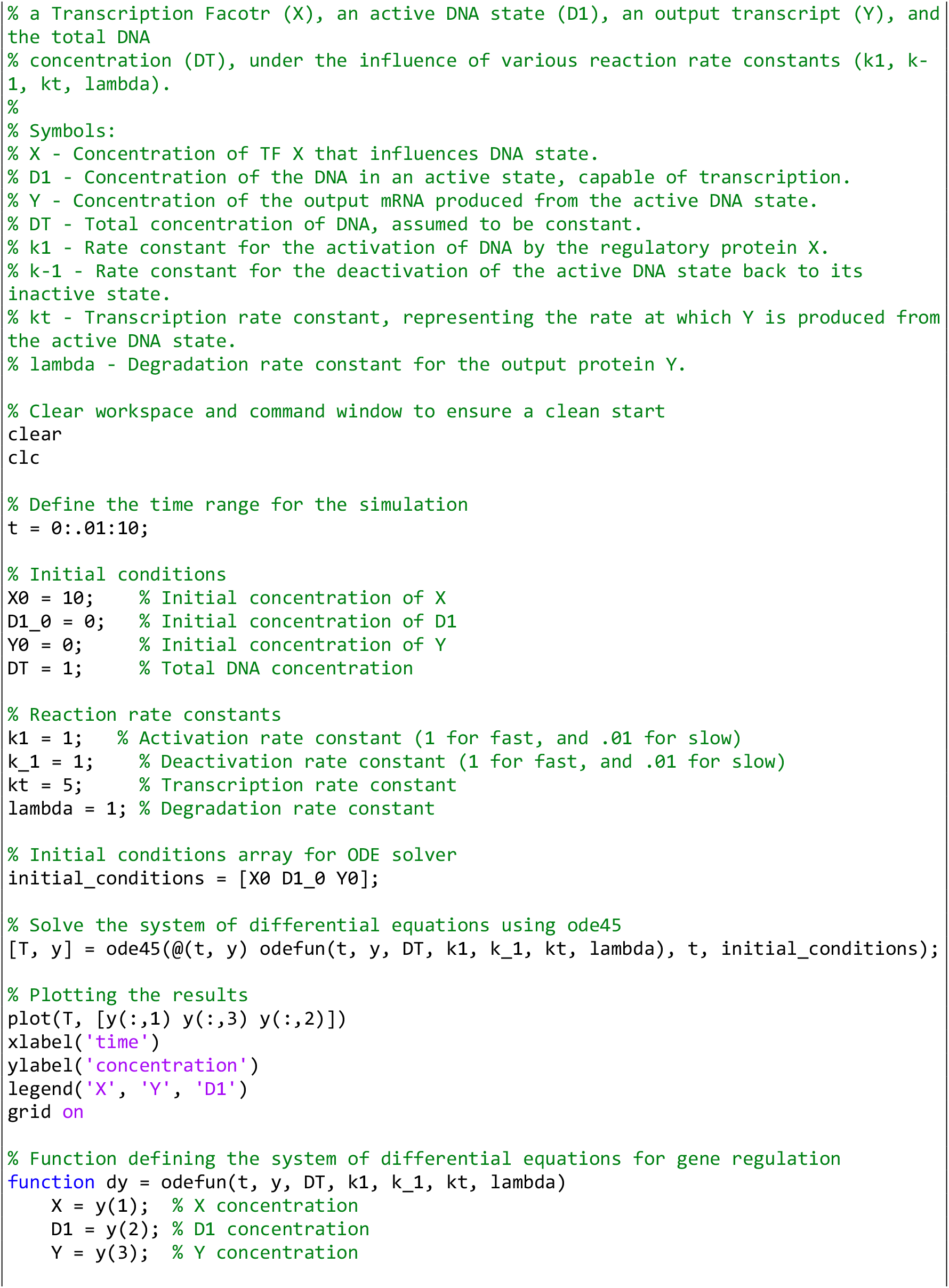
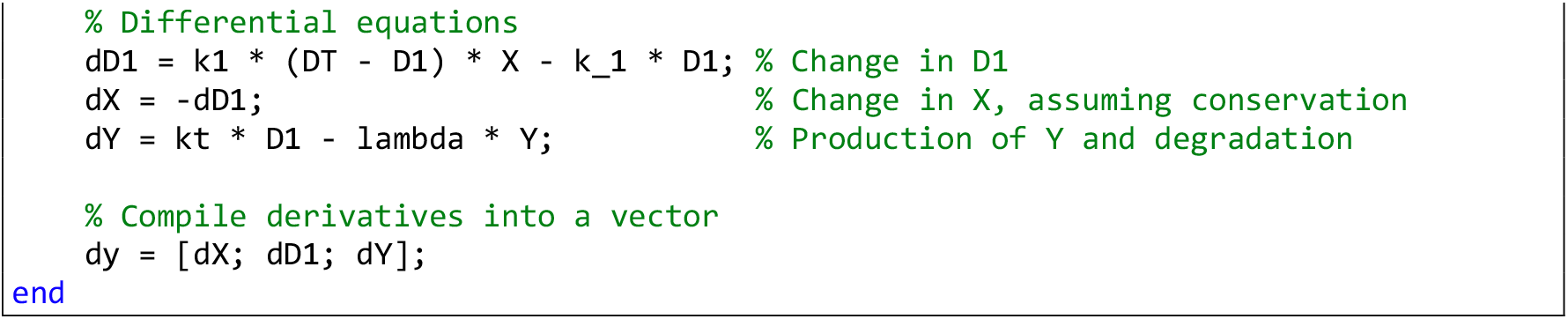

**Computer Simulation 4**. Modeling Gene Activation Using Michaelis-Menten Kinetics.

The results of this computer simulation are shown in Figure S2-C.

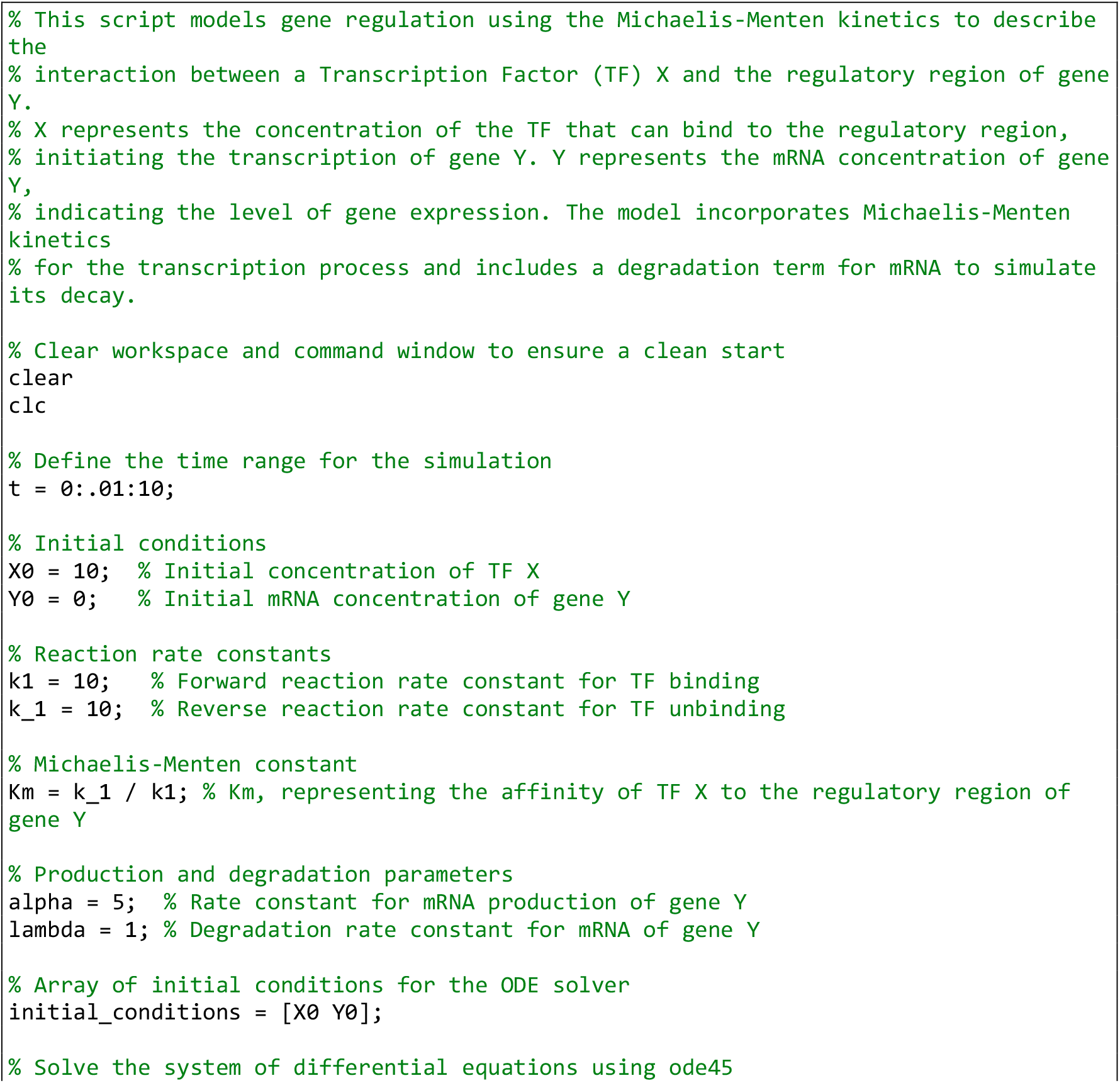
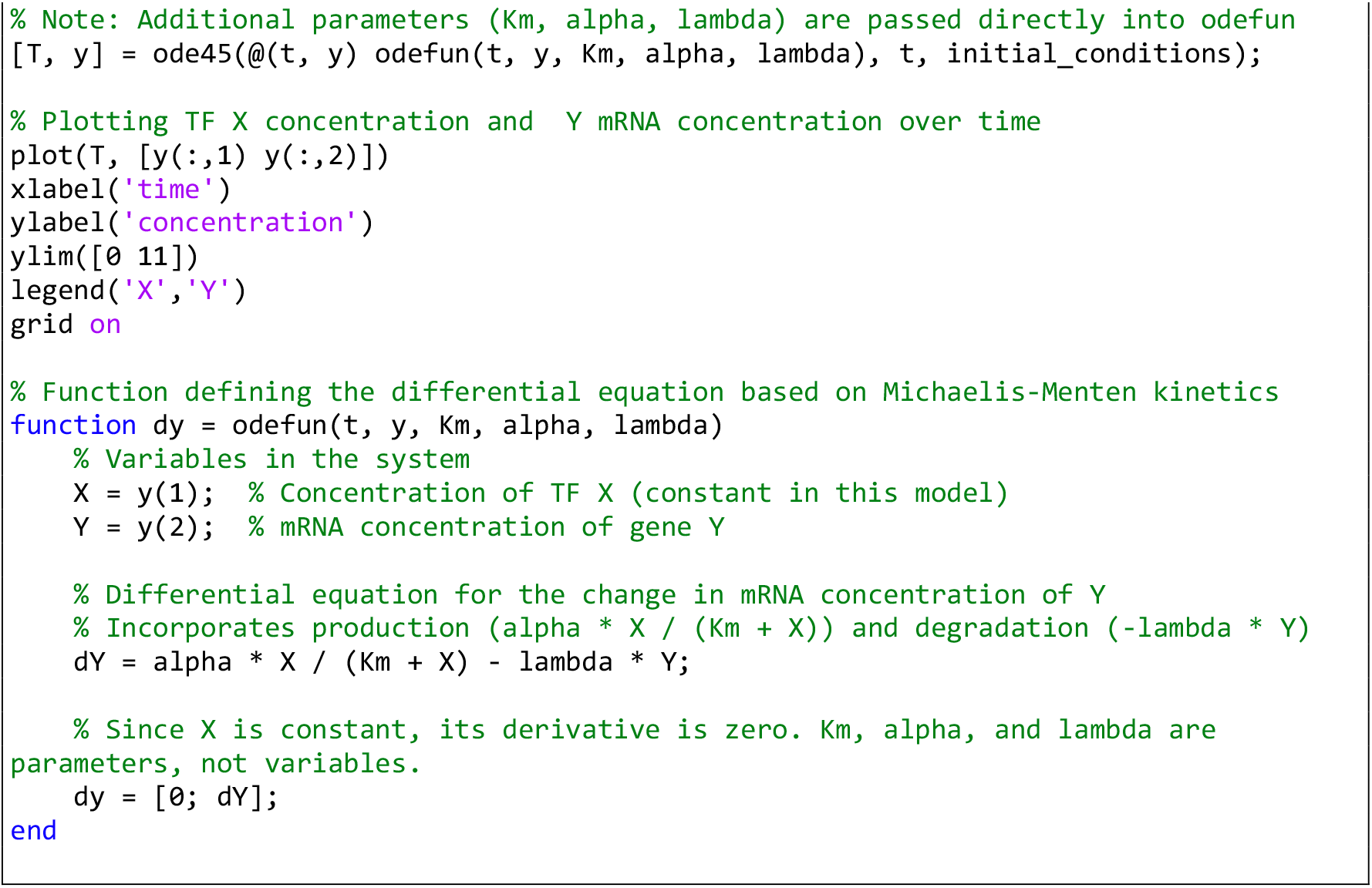

**Computer Simulation 5**. Input-Output Relationship Between Activator Concentration and Gene Expression Using Michaelis-Menten Dynamics.

The results of this computer simulation are shown in Figure 1B.

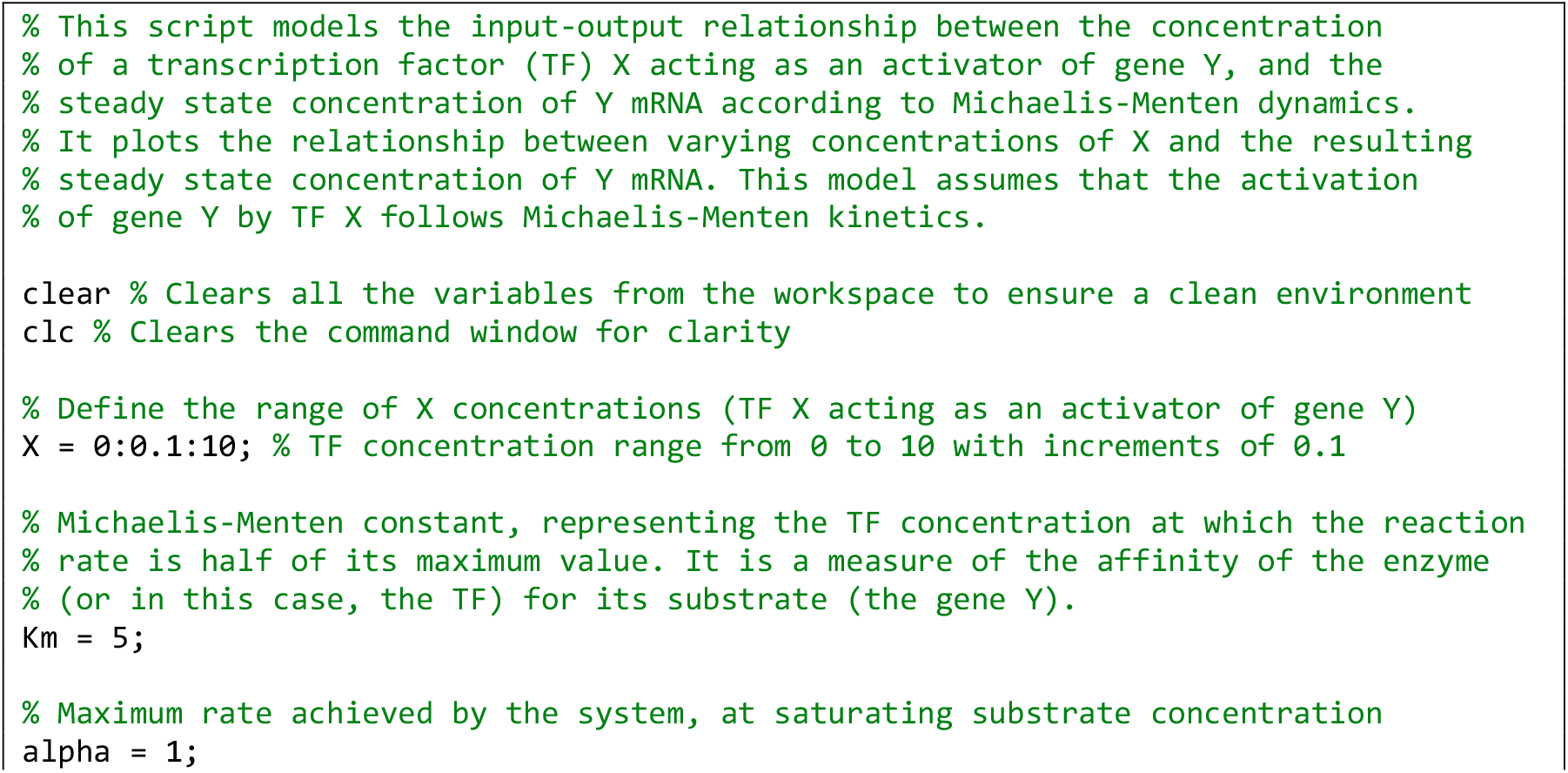
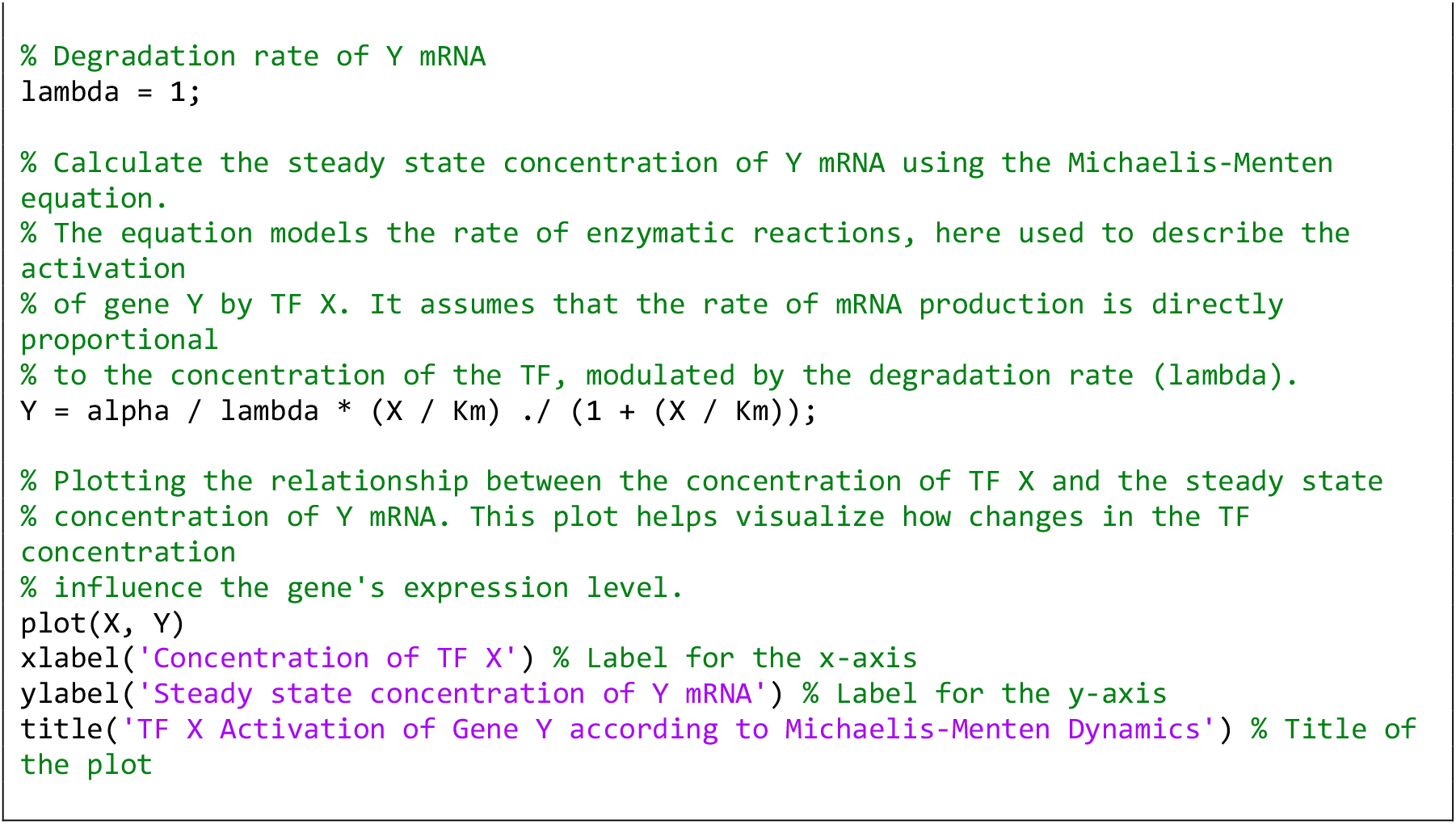

**Computer Simulation 6**. Input-Output Relationship Between Repressor Concentration and Gene Expression.

The results of this computer simulation are shown in Figure 1D.

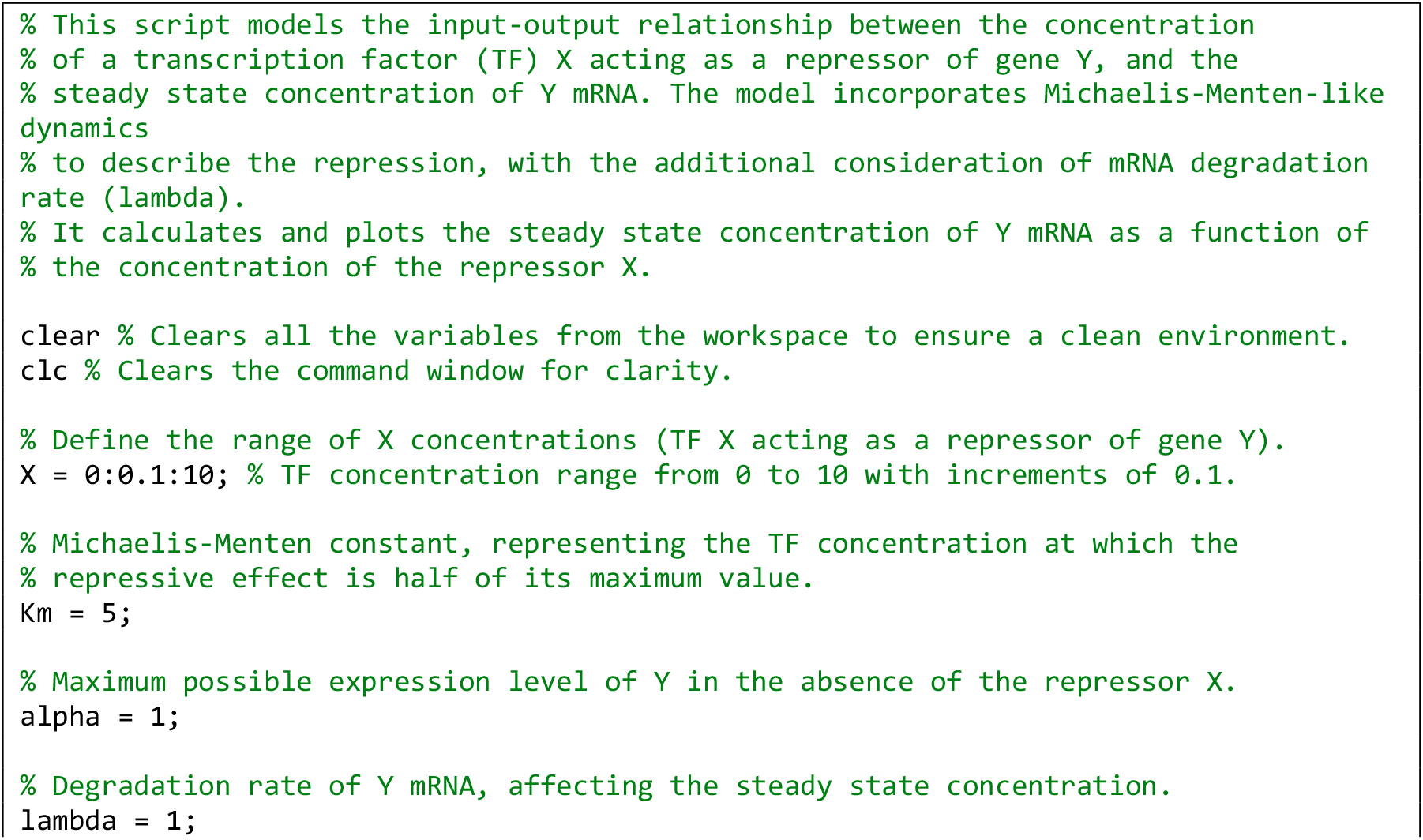
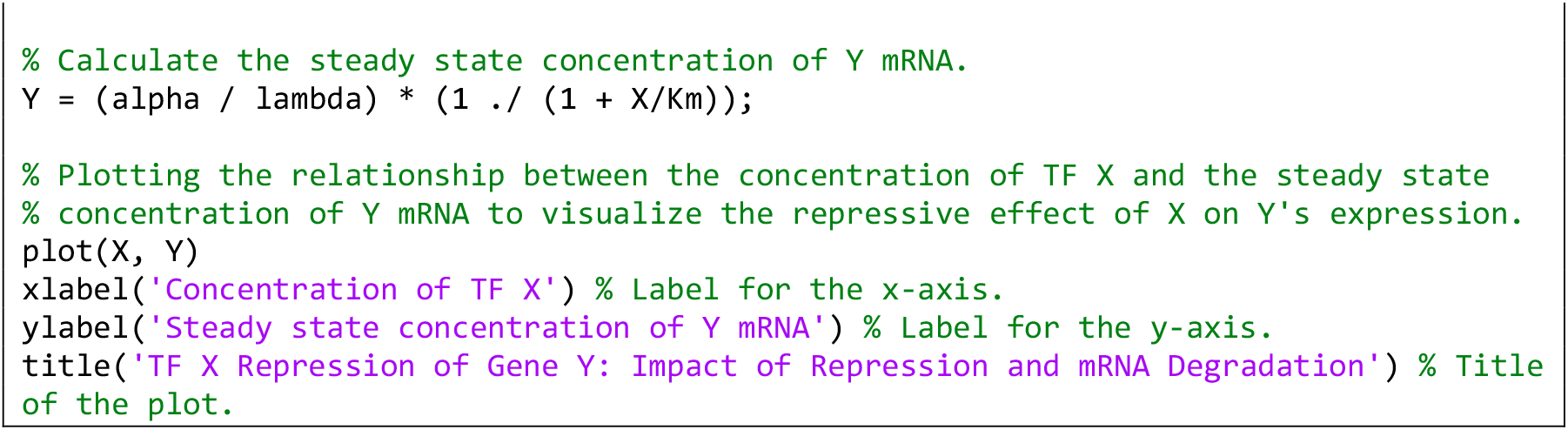

**Computer Simulation 7**. Effect of Cooperative Binding on Gene Activation Using the Hill Function.

The results of this computer simulation are shown in Fig. 1F (*n*=2) and Fig. 1G (*n*=5).

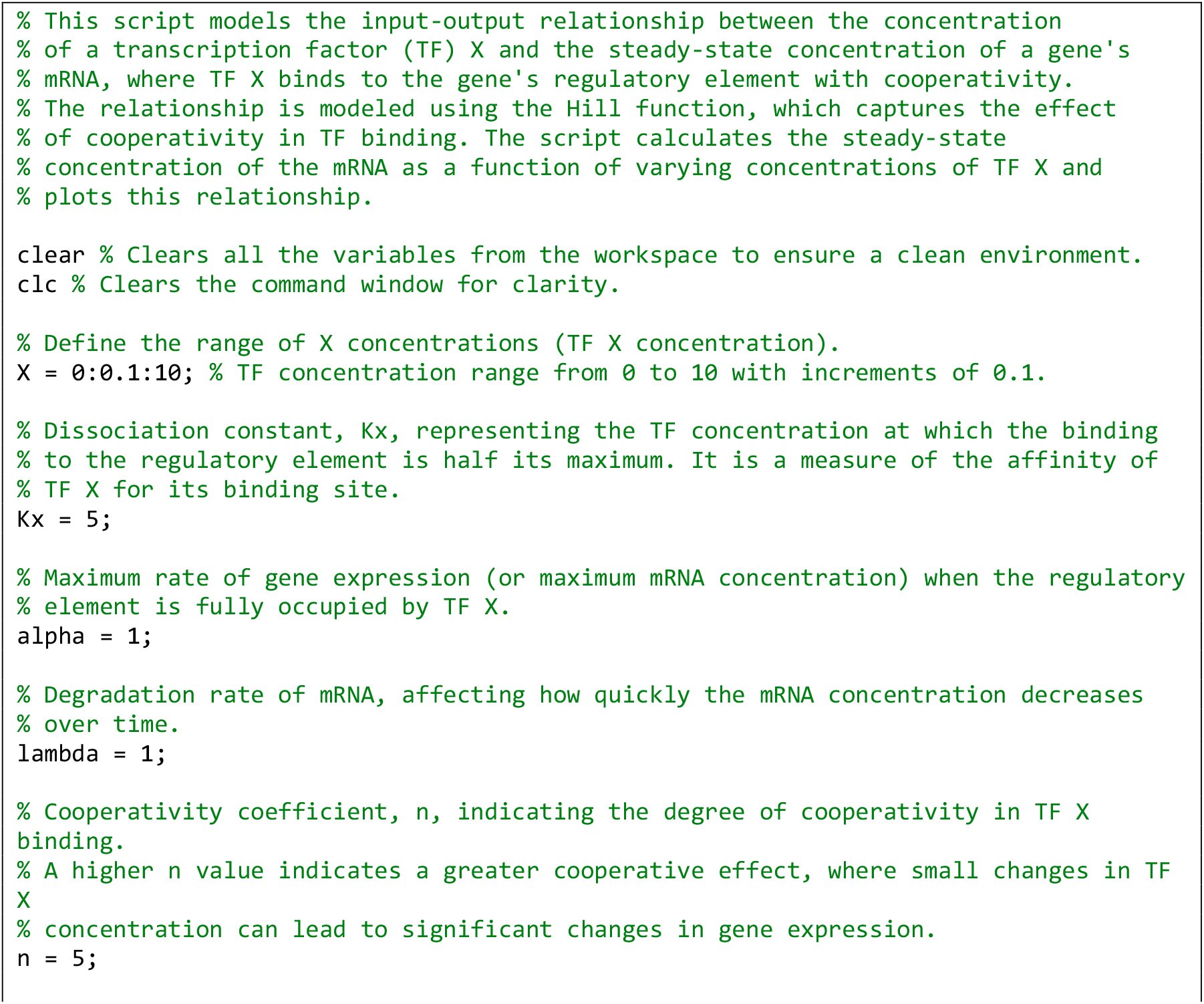
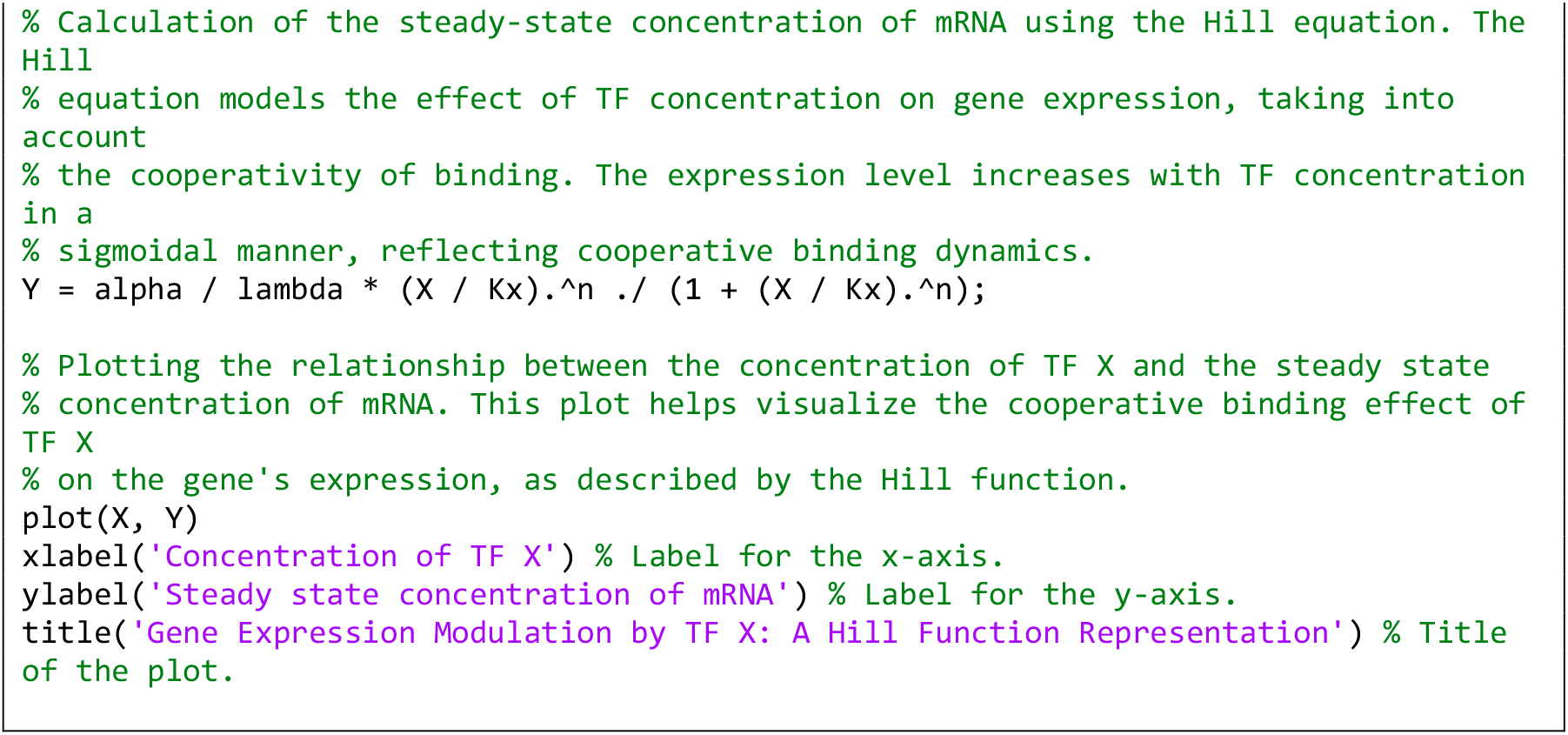

**Computer Simulation 8**. Modeling Gene Activation by Two ORed Activators Using Hill Functions.

The results of this computer simulation are shown in Fig. 2B.

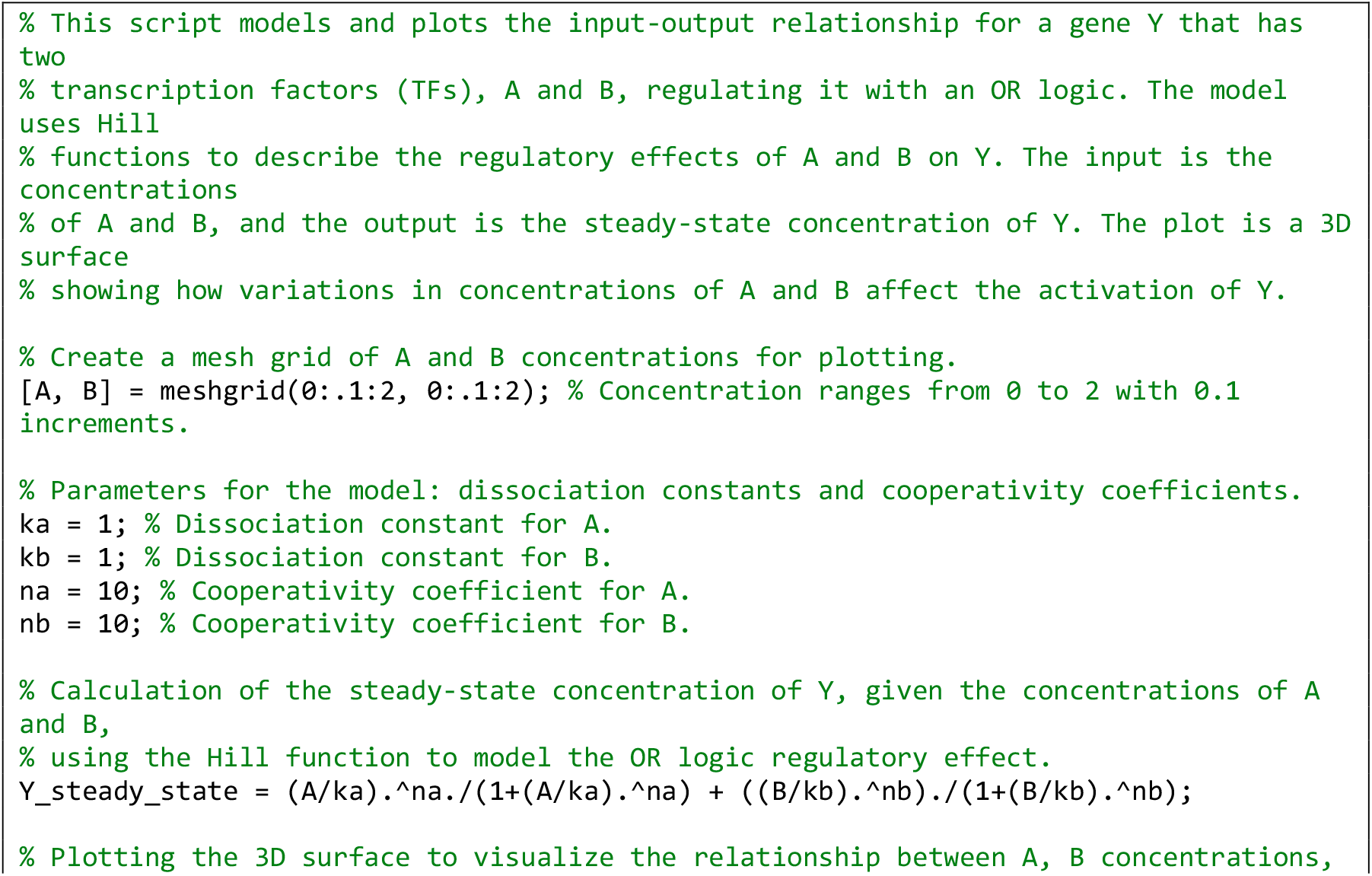
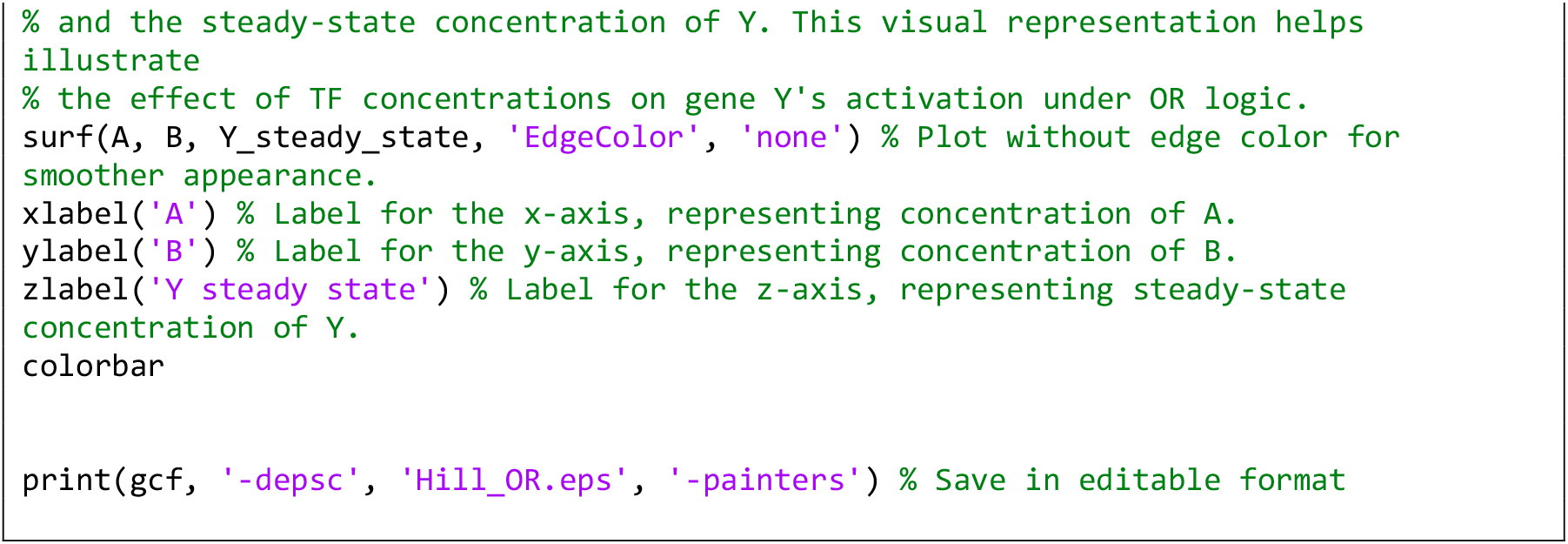

**Computer Simulation 9**. Modeling Gene Activation by Two Competitive ORed Activators.

The results of this computer simulation are shown in Fig. 2D.

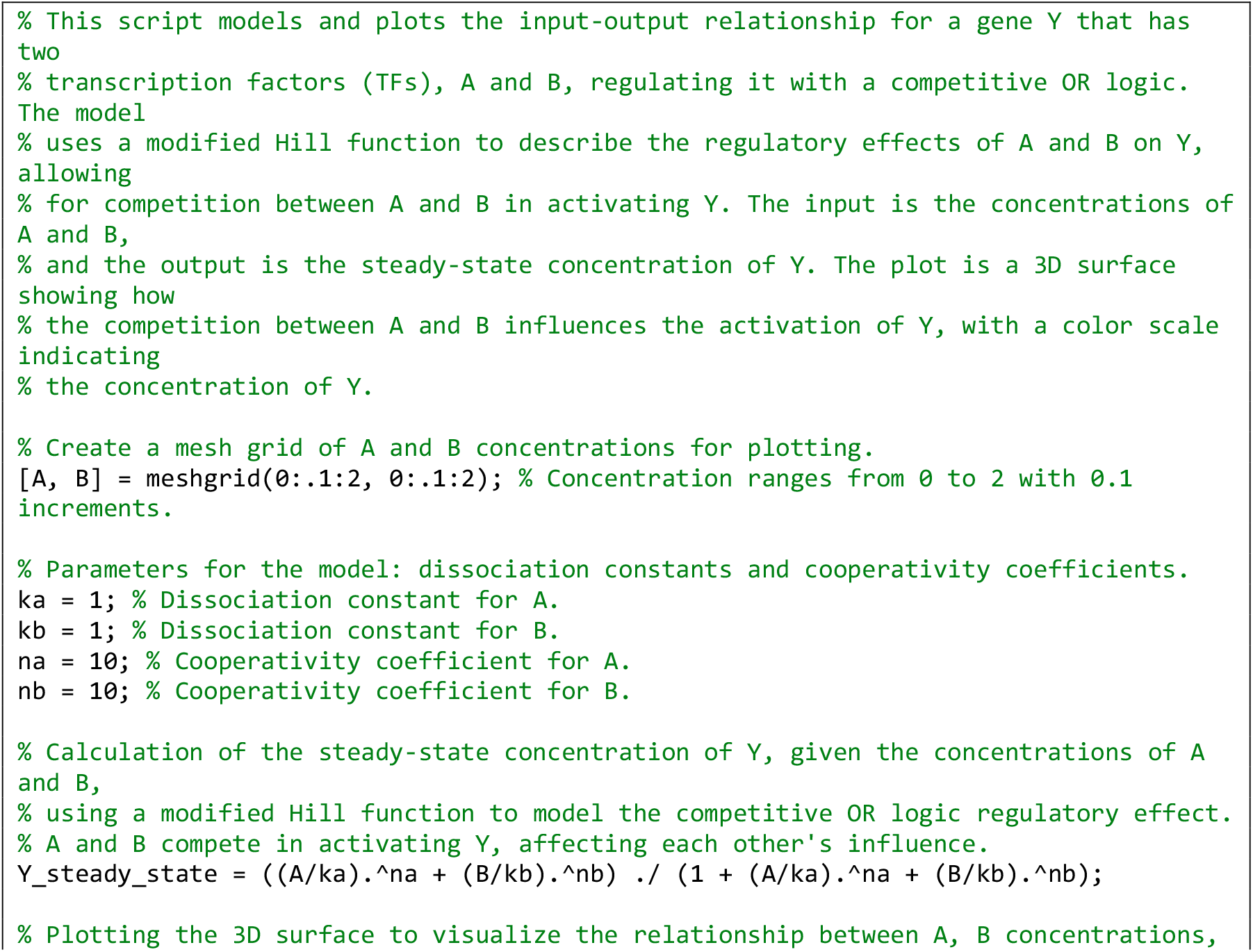
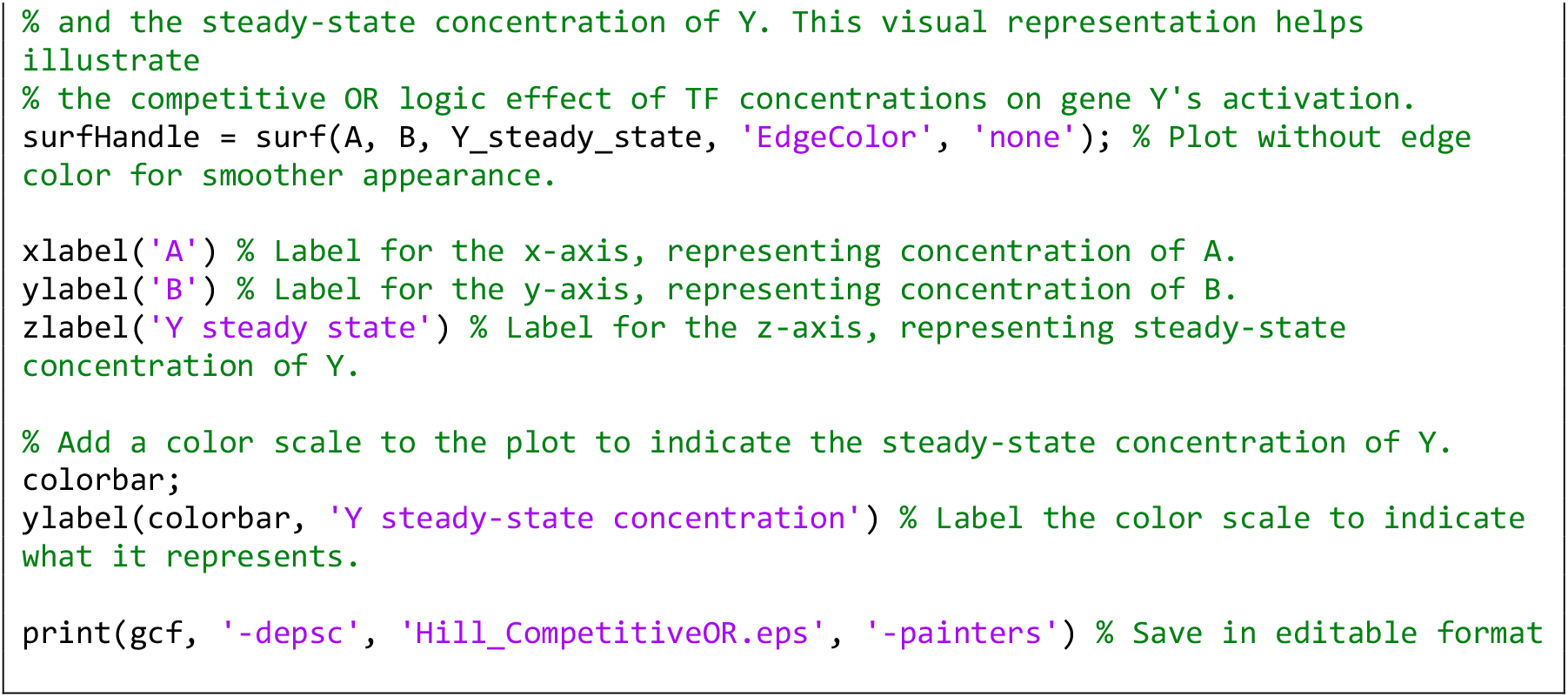

**Computer Simulation 10**. Modeling Gene Activation by Two ANDed Activators.

The results of this computer simulation are shown in Fig. 2F.

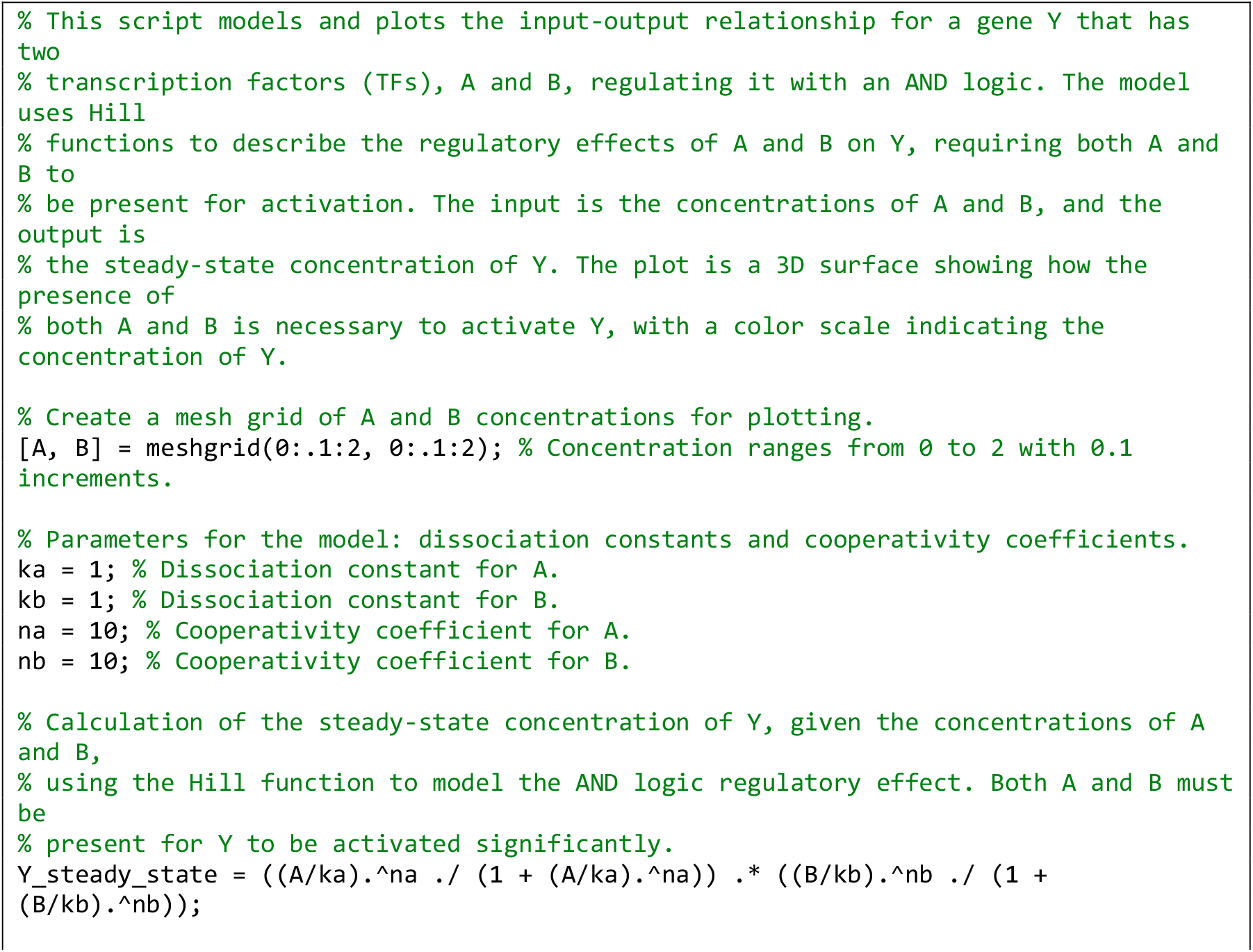
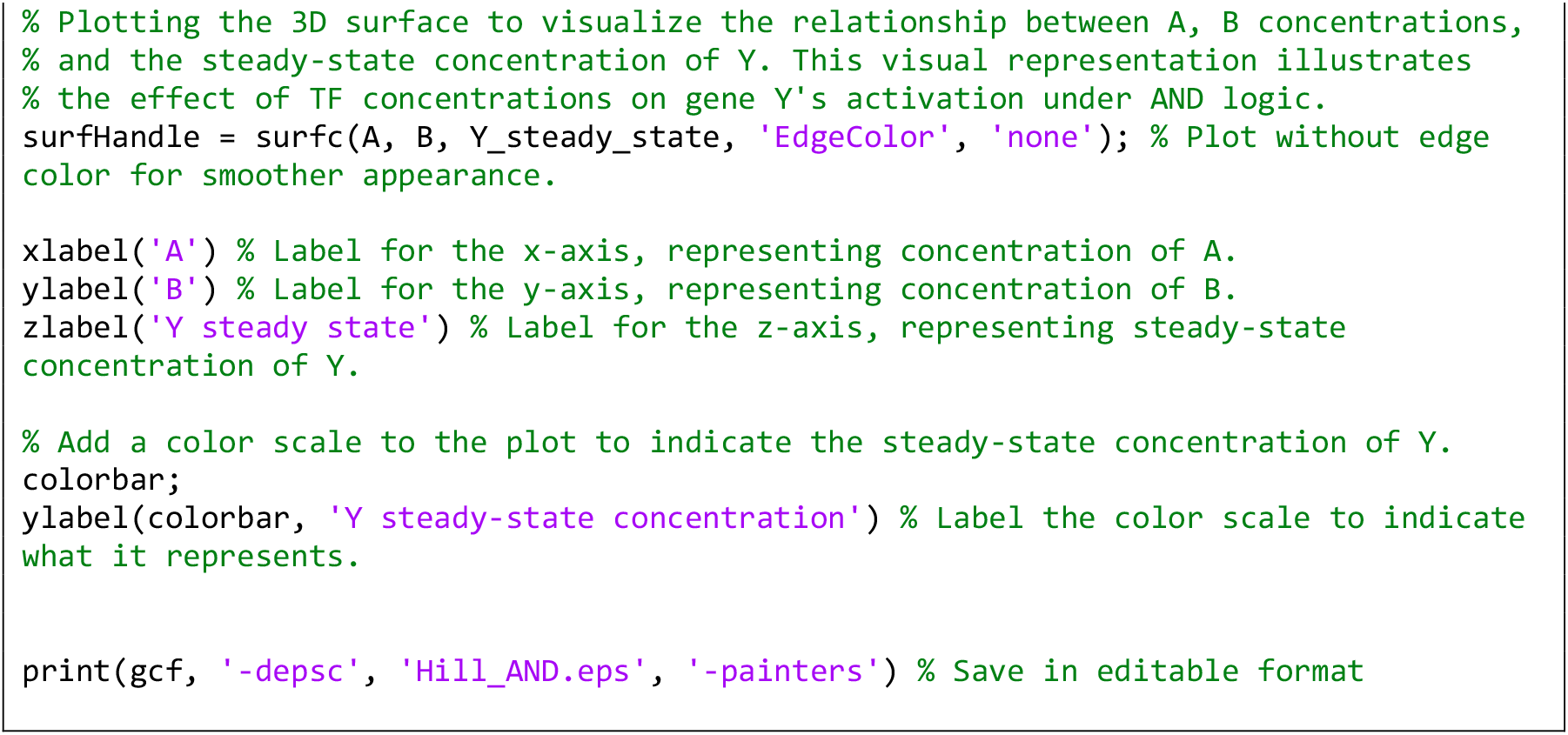

**Computer Simulation 11**. Modeling a Positive Feedback Loop Between Two Activator Genes.

The results of this computer simulation are shown in Fig. 3B’ (X0=0.5, Y0=0.8), and 3B’’ (X0=1.2, Y0=1.7).

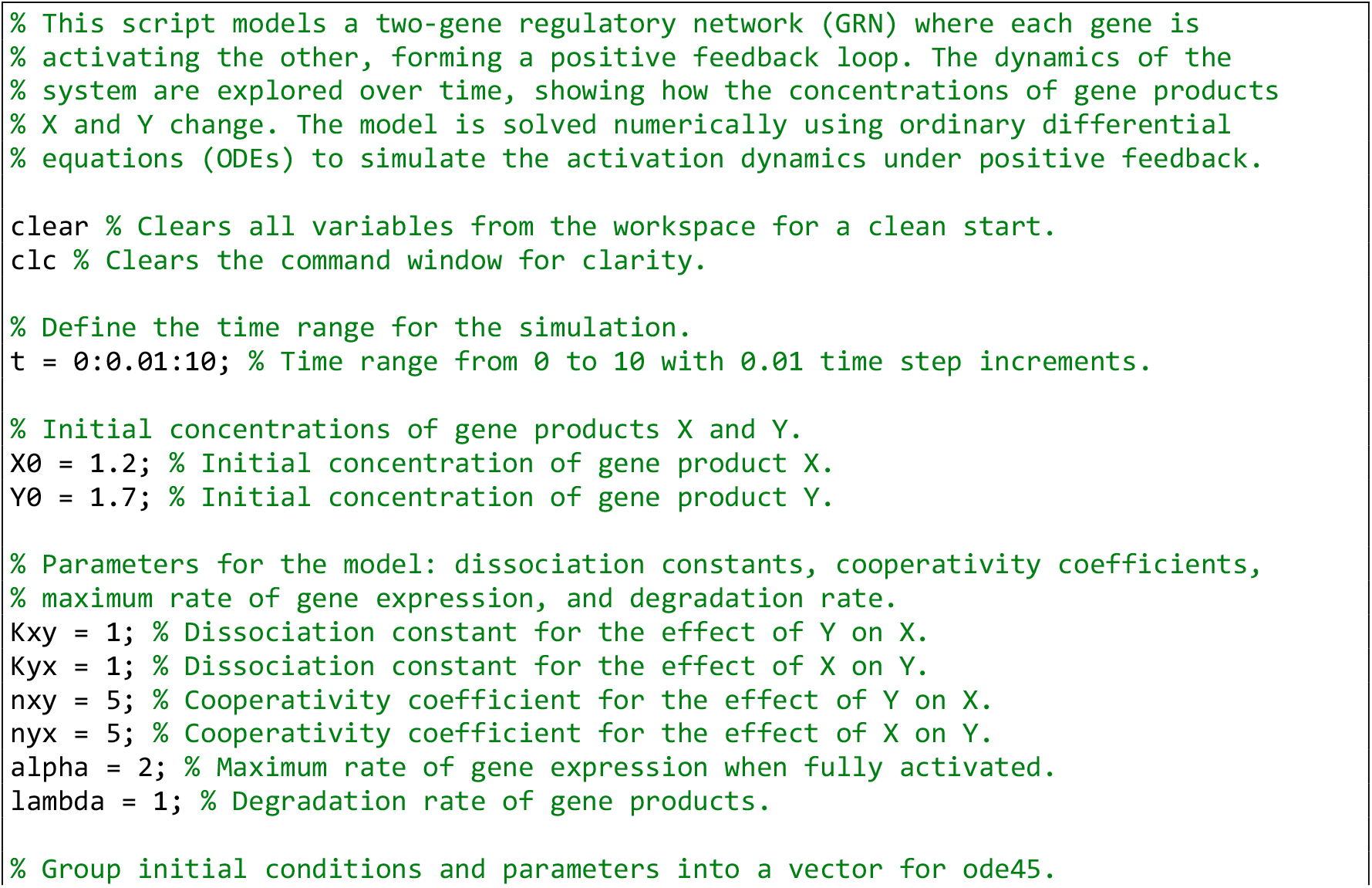
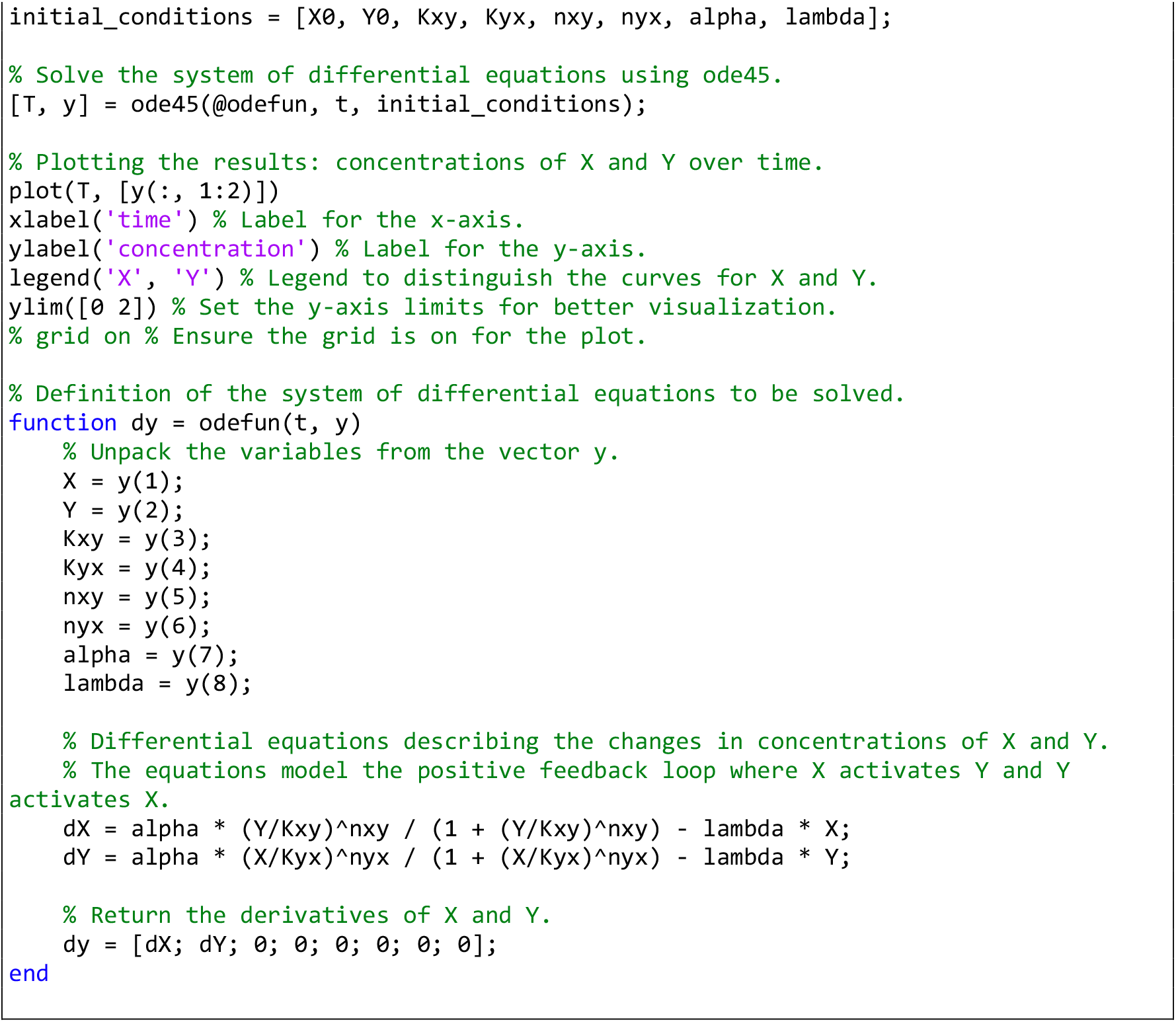

**Computer Simulation 12**. Modeling a Mutual Repression Loop Between Two Repressor Genes.

The results of this computer simulation are shown in Fig. 3C’ (X0=0.8, Y0=1.5), and 3C’’ (X0=1.5, Y0=0.8).

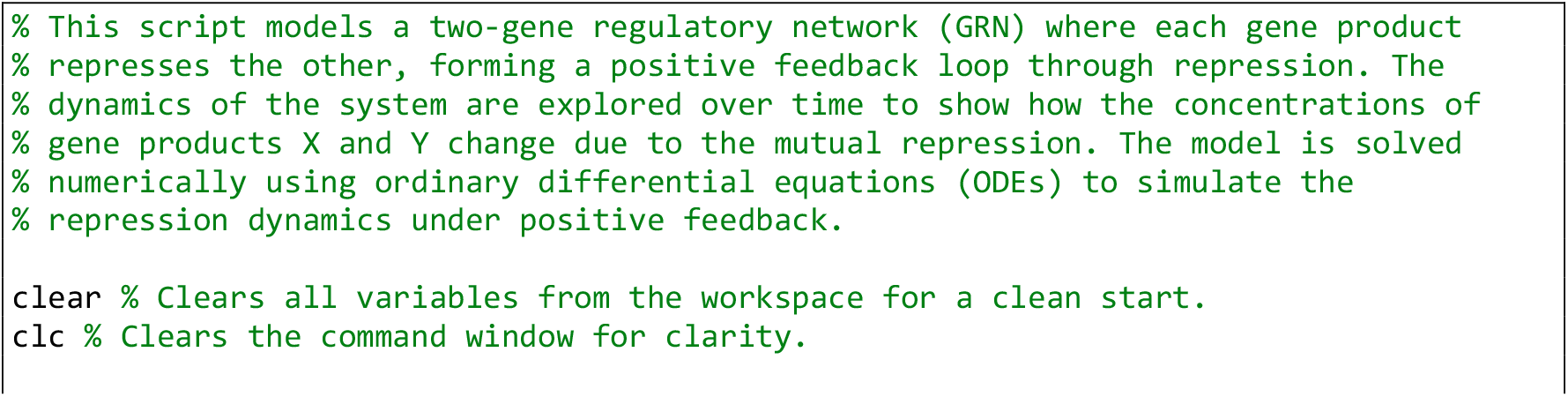
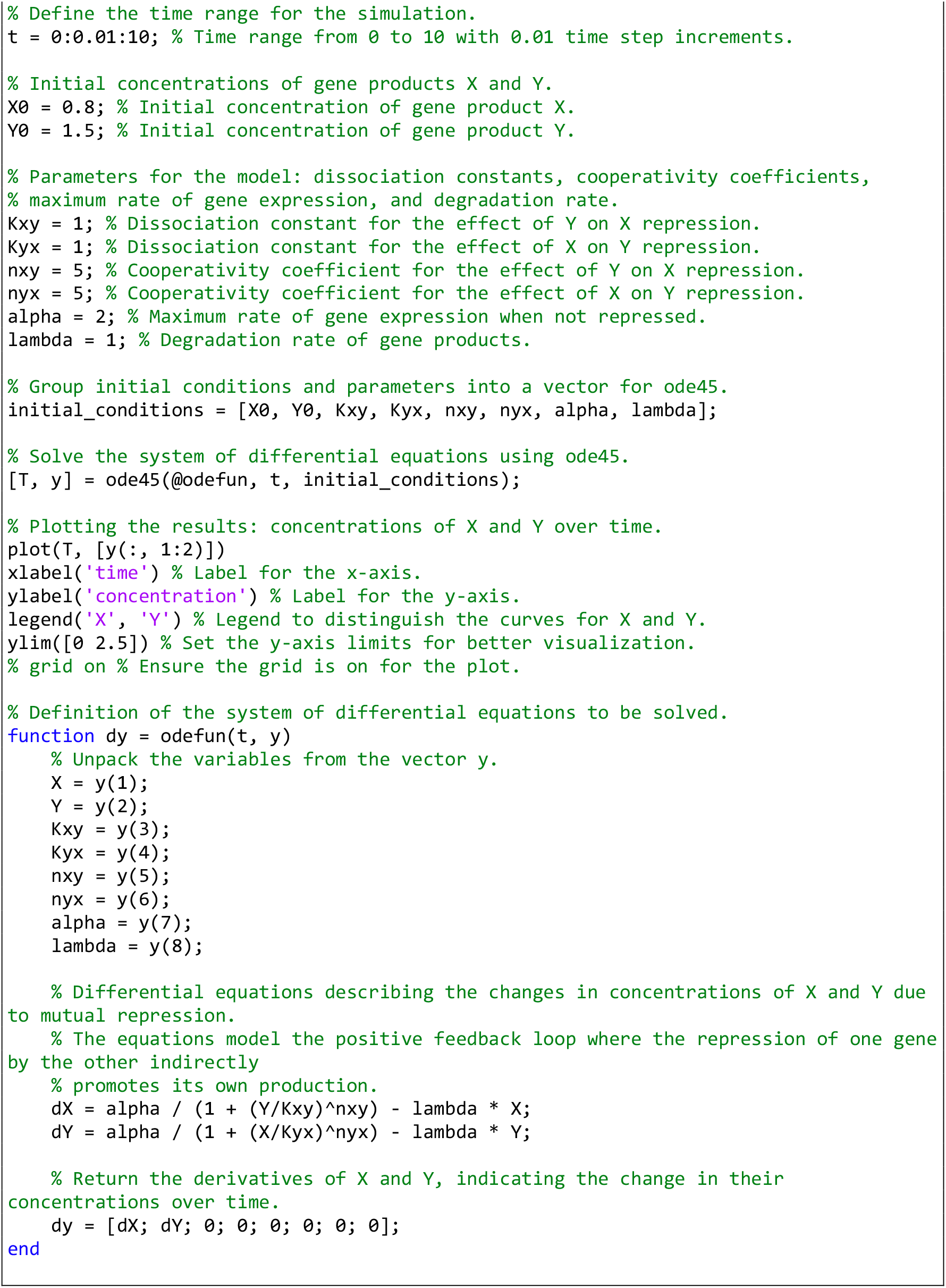

**Computer Simulation 13**. Modeling a Negative Feedback Loop Between an Activator and a Repressor Gene.

The results of this computer simulation are shown in Fig. 3D’.

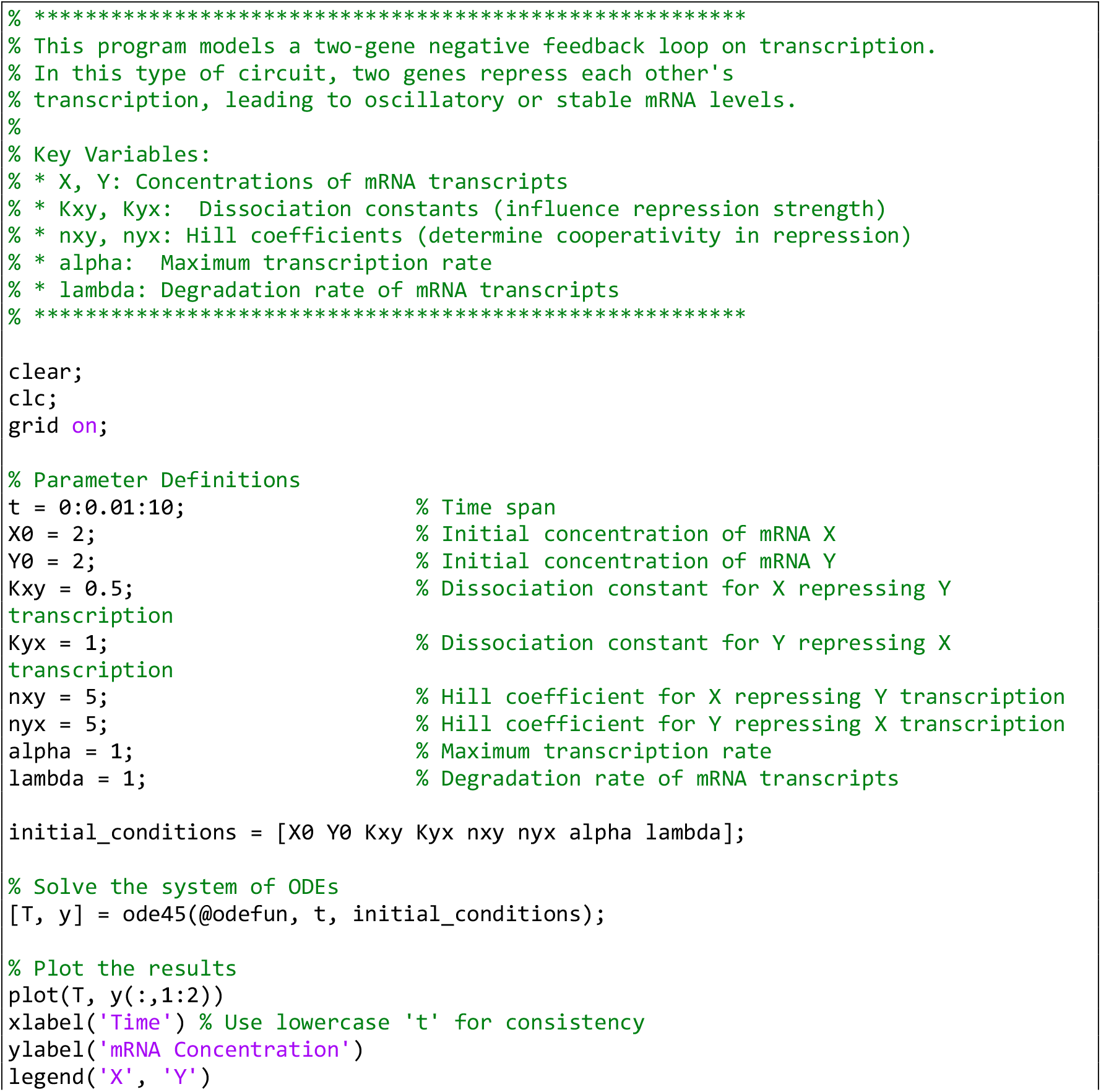
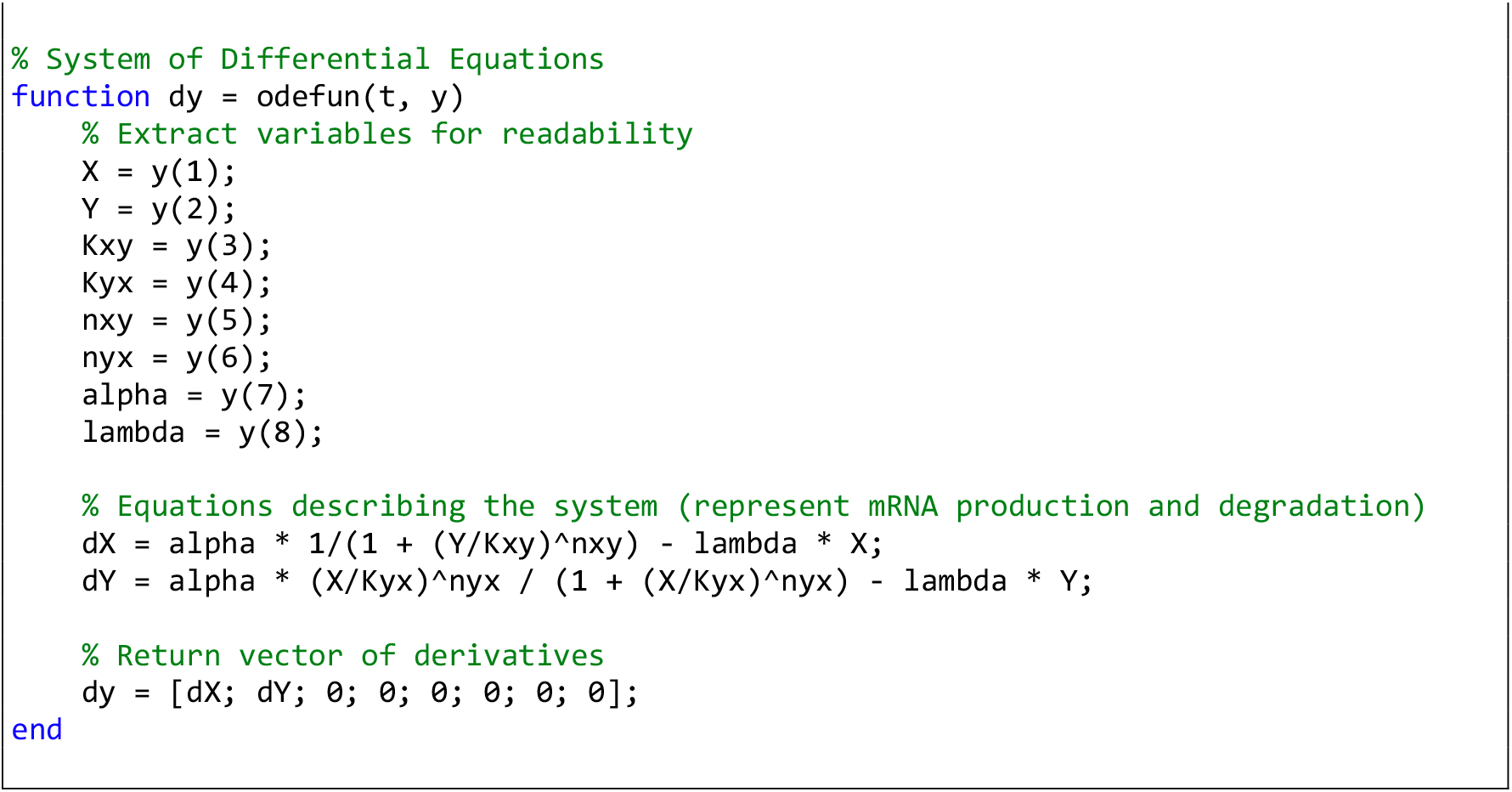

**Computer Simulation 14**. Modeling an Oscillatory Gene Network with Two Activators and One Repressor.

The results of this computer simulation are shown in Fig. 3E’.

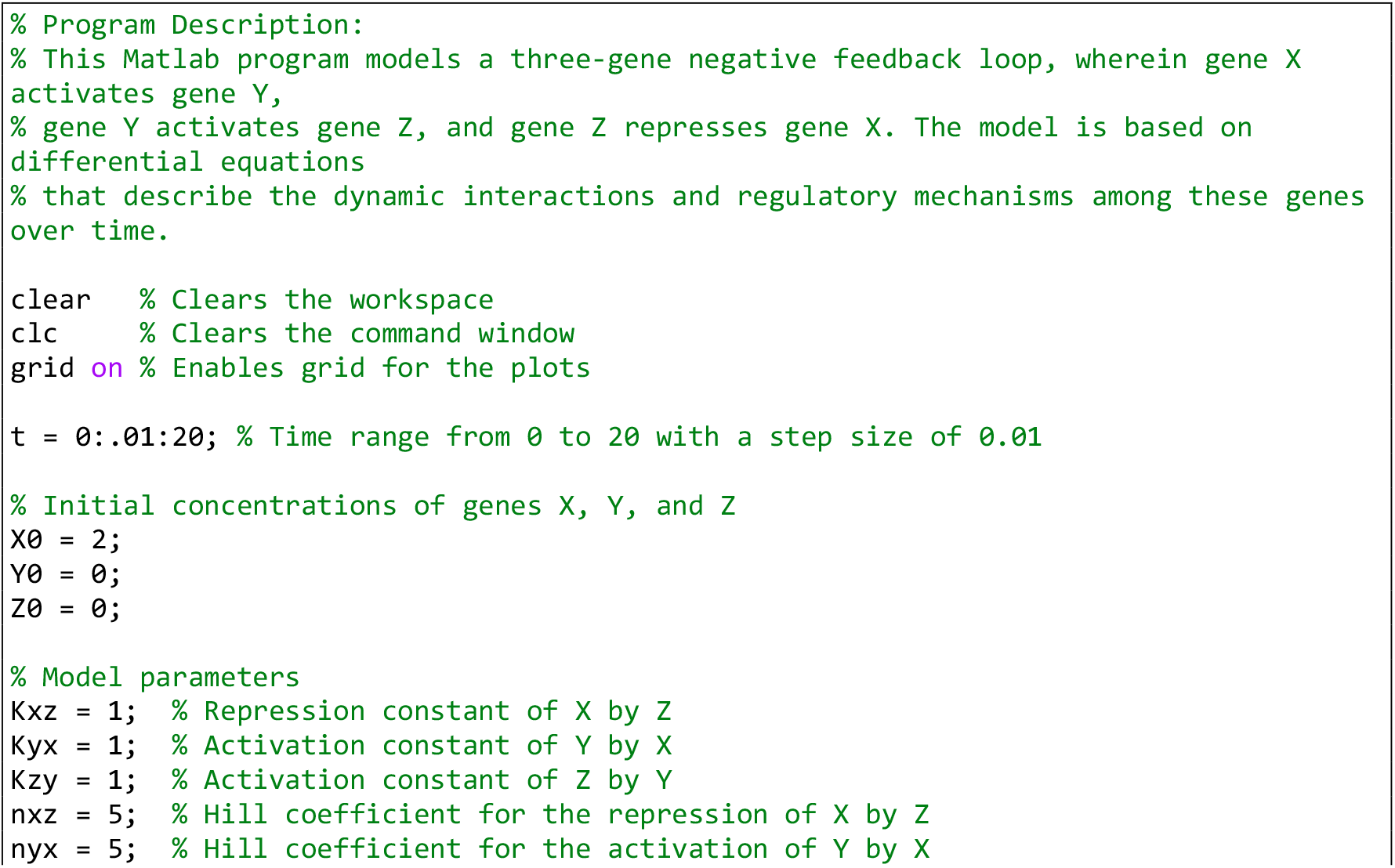
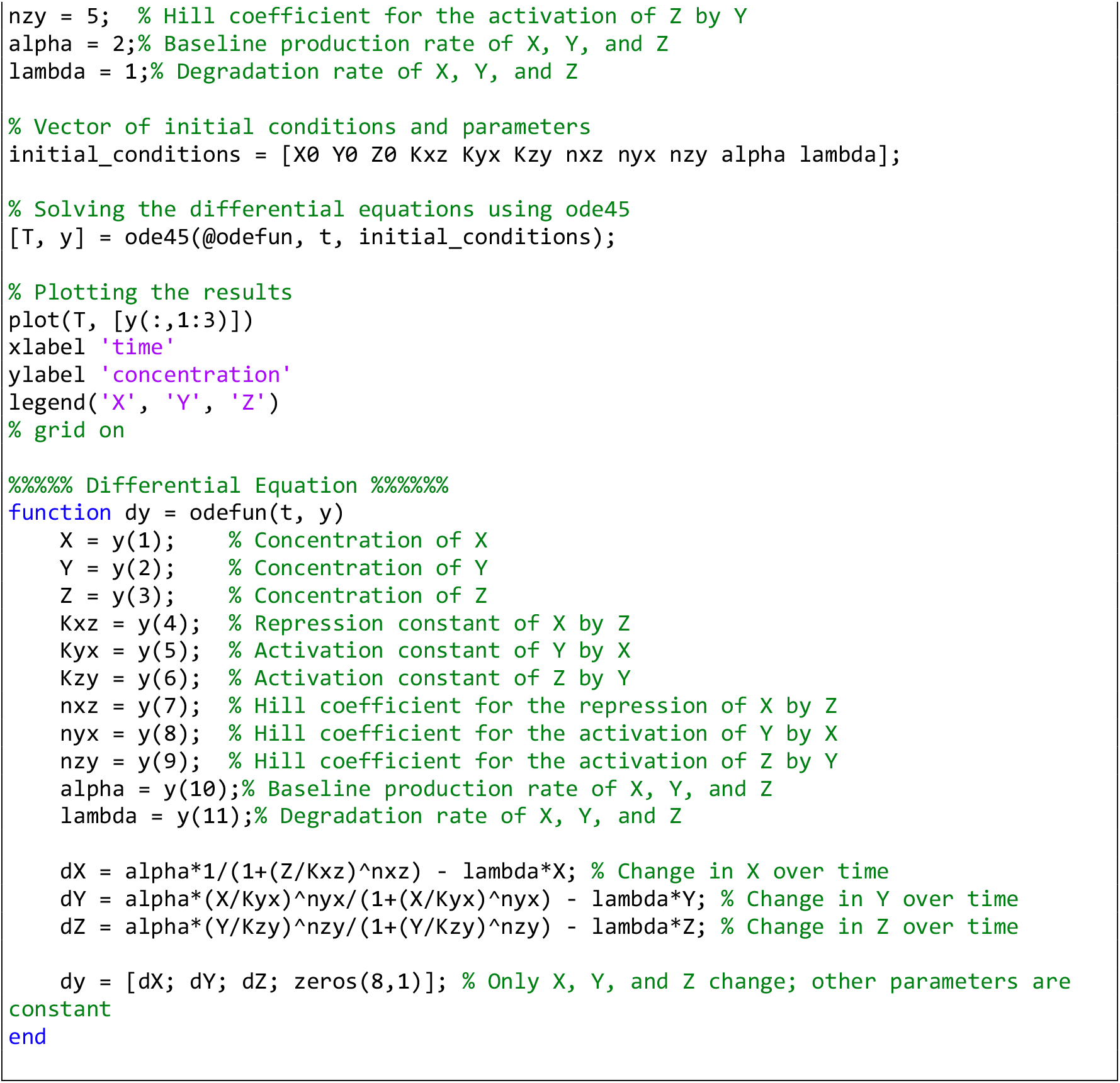

**Computer Simulation 15**. Modeling the Repressilator Network Composed of Three Repressor Genes.

The results of this computer simulation are shown in Fig. 3F’.

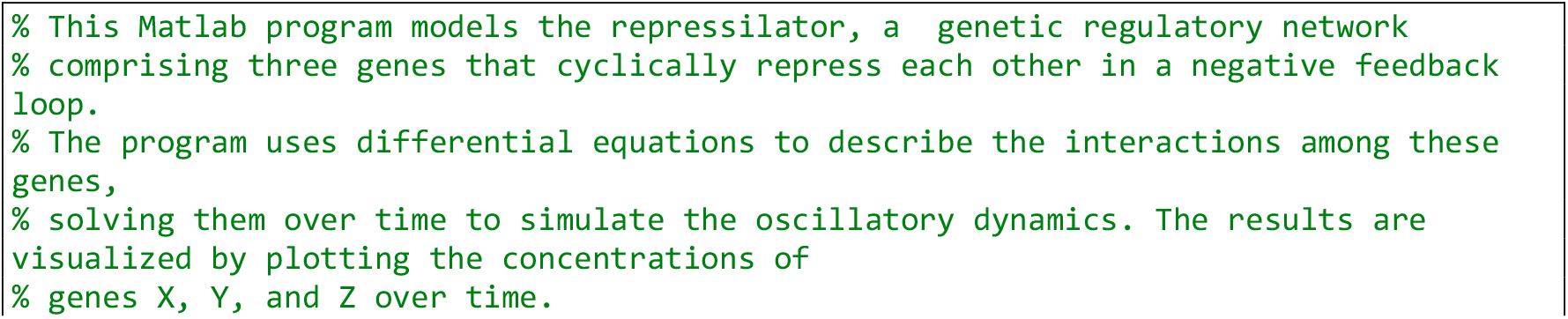
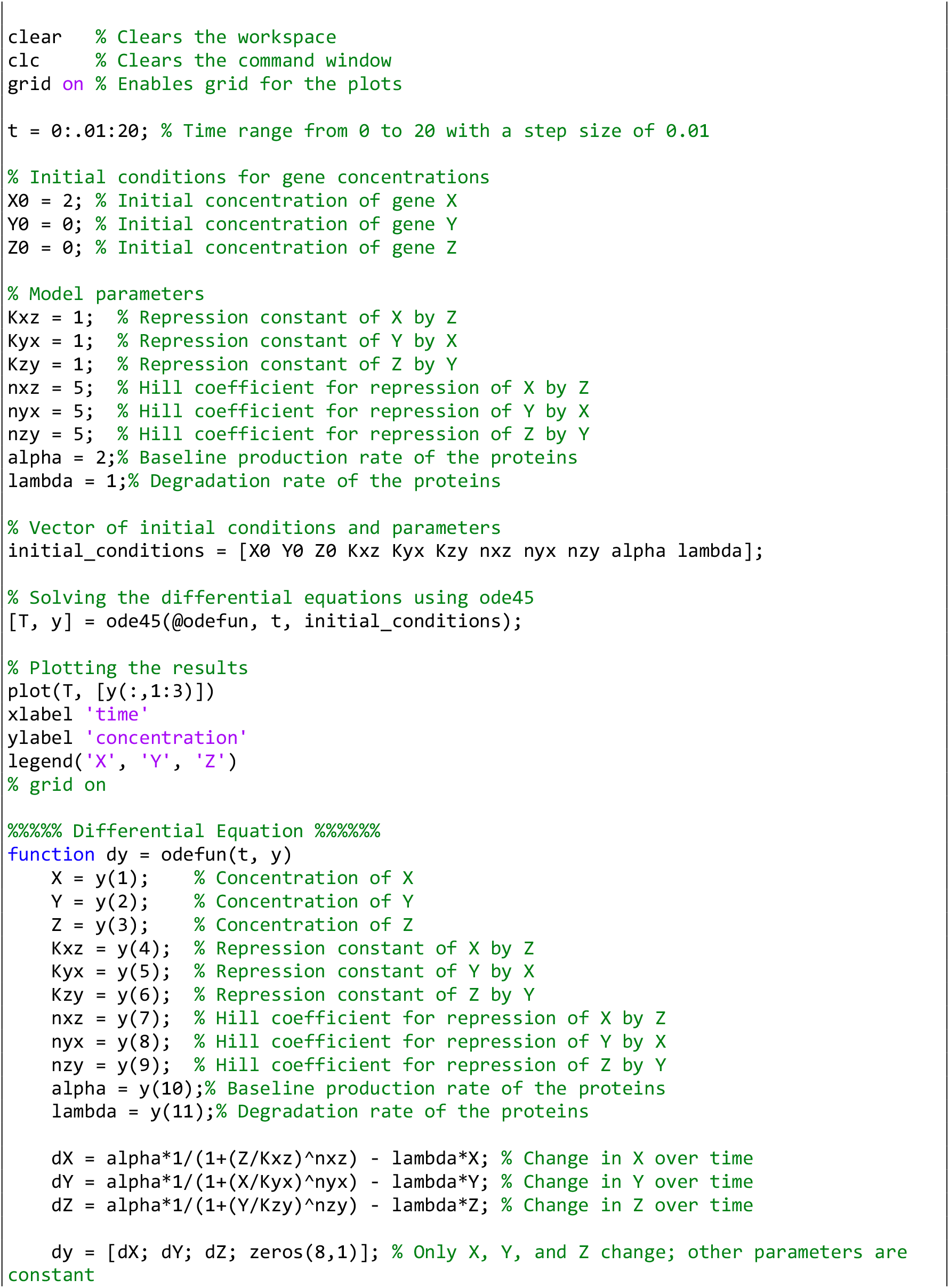
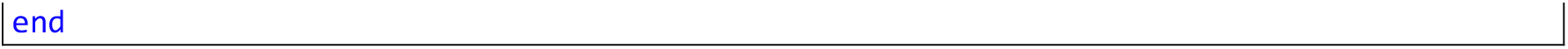

**Computer Simulation 16**. Modeling a Multi-Stable Gene Regulatory Network with Mutually Repressing Genes.

The results of this computer simulation are shown in Fig. 3G’.

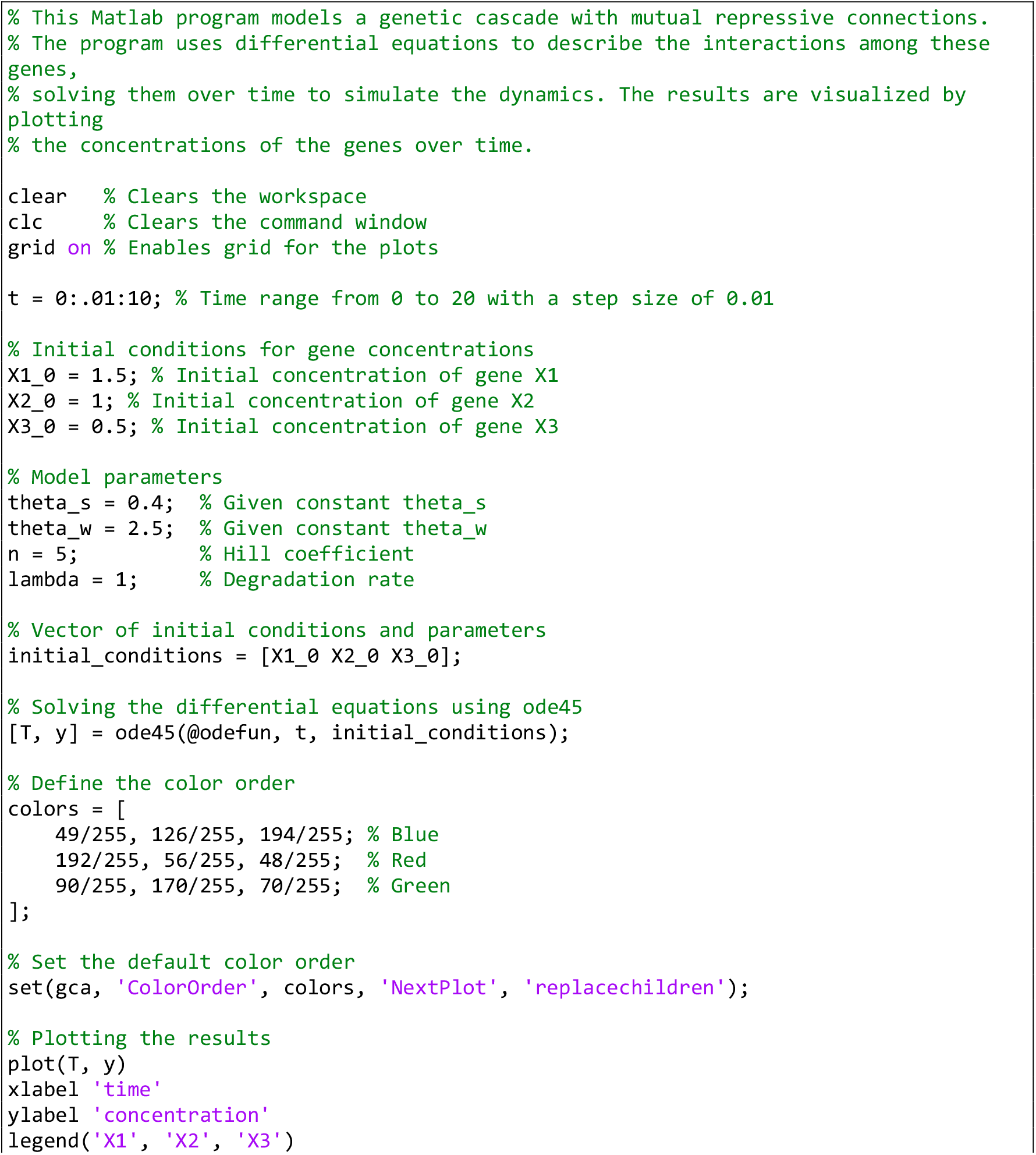
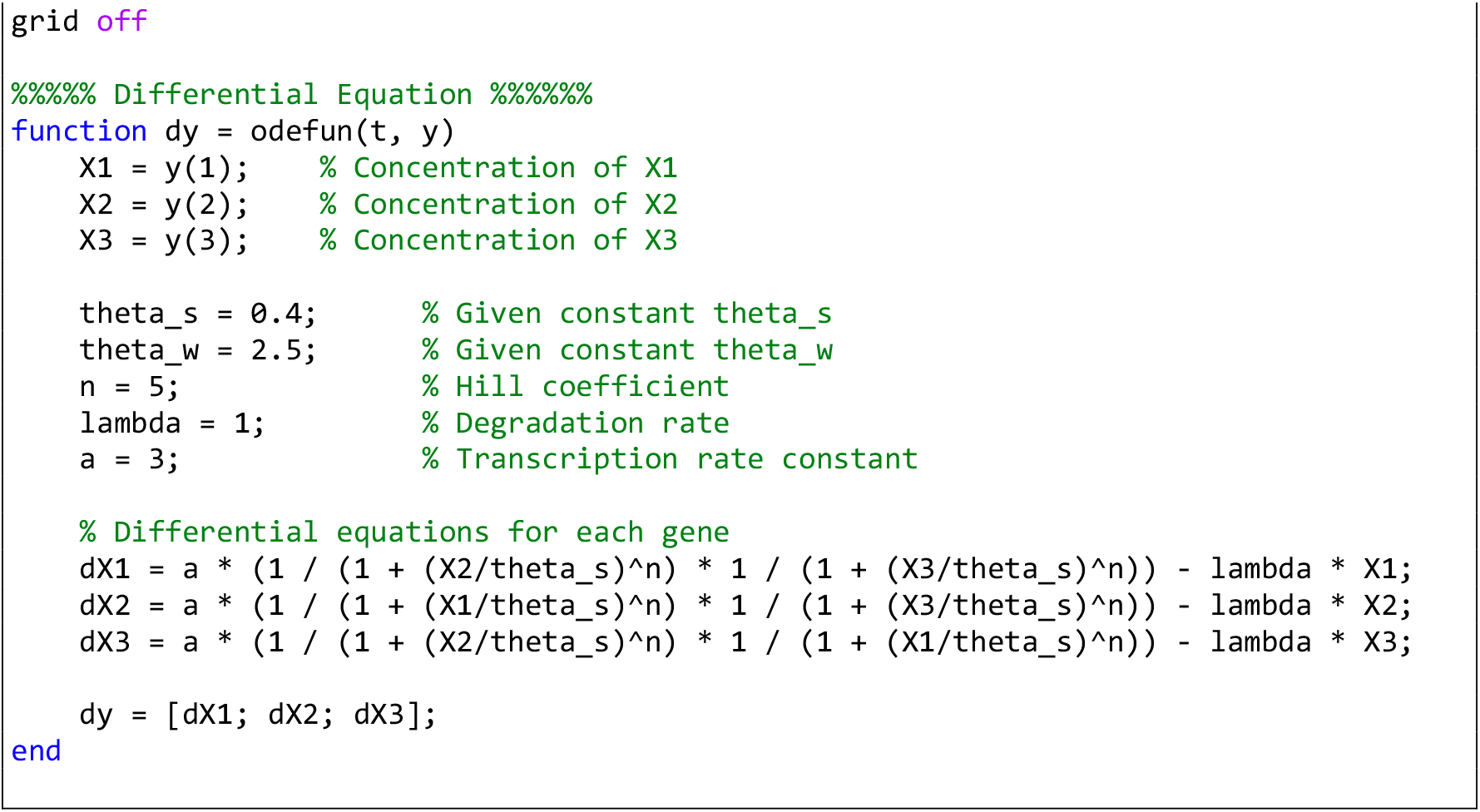

**Computer Simulation 17**. Modeling a Genetic Cascade Illustrating Sequential Gene Activation.

The results of this computer simulation are shown in Fig. 3H’.

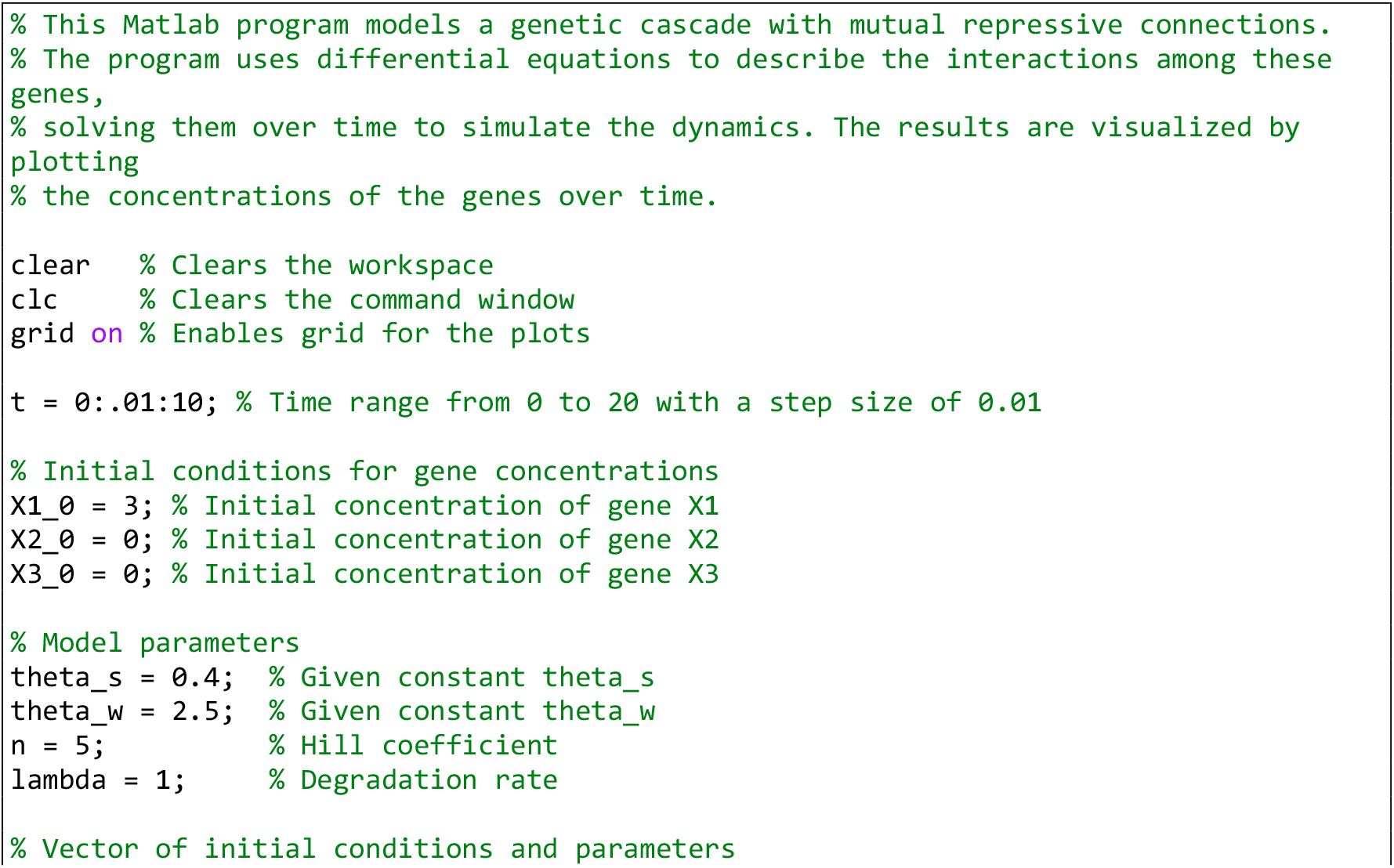
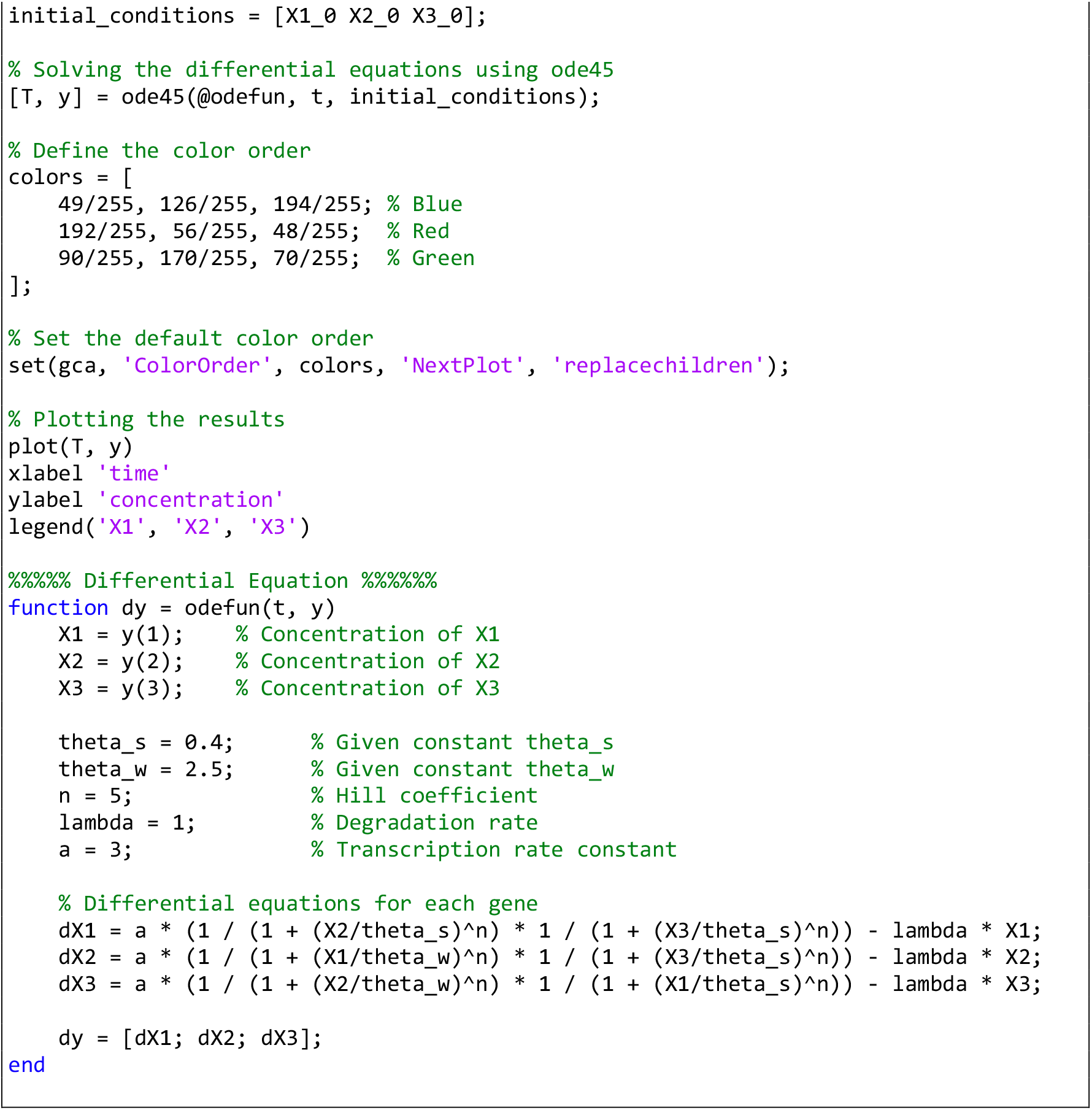

**Computer Simulation 18**. Modeling an Oscillatory Gene Network with Mutual Repression.

The results of this computer simulation are shown in Fig. 3I’.

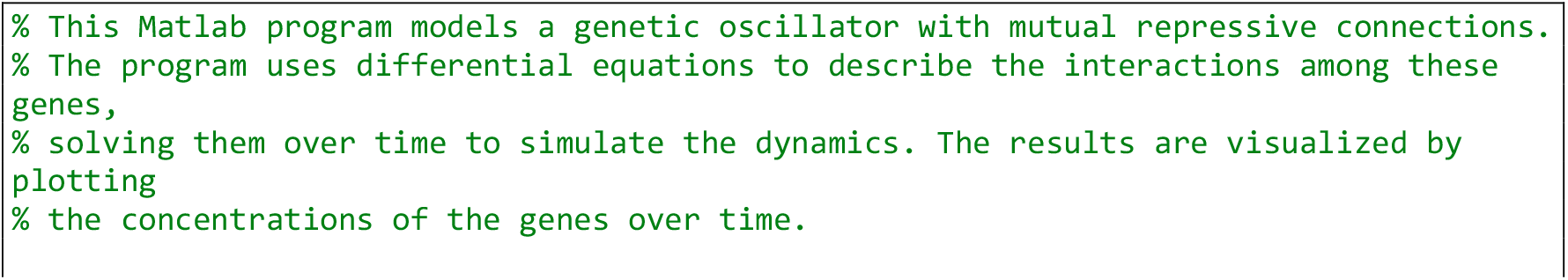
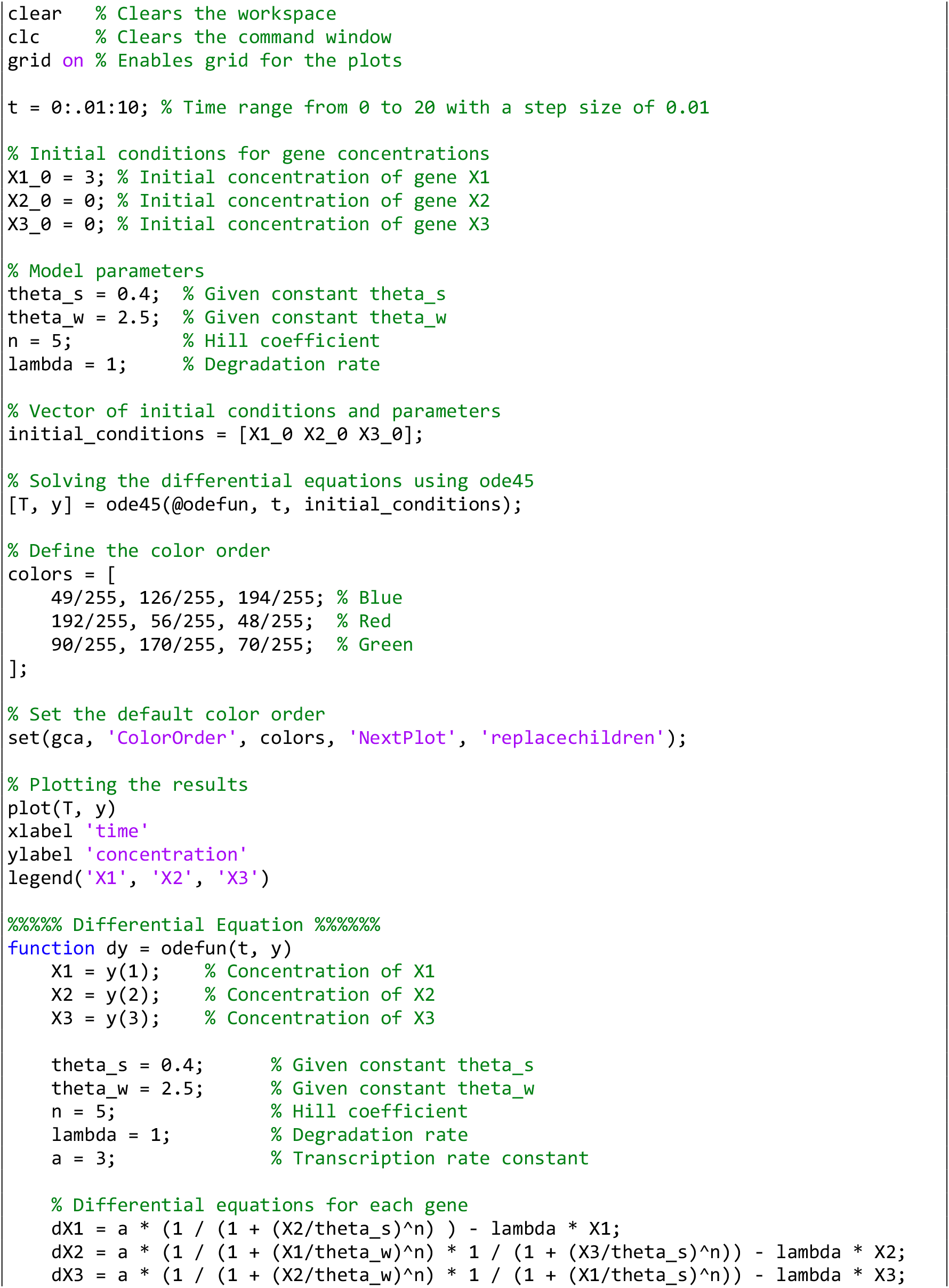
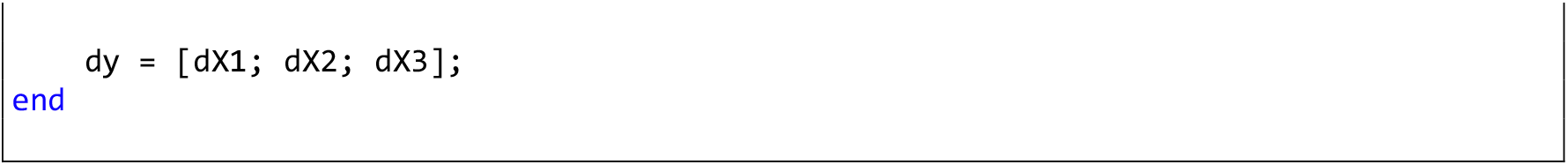

**Computer Simulation 19**. Gene Regulatory Network Realization of the French Flag Model in a Non-Elongating Tissue.

The results of this computer simulation are shown in Fig. 5C’.

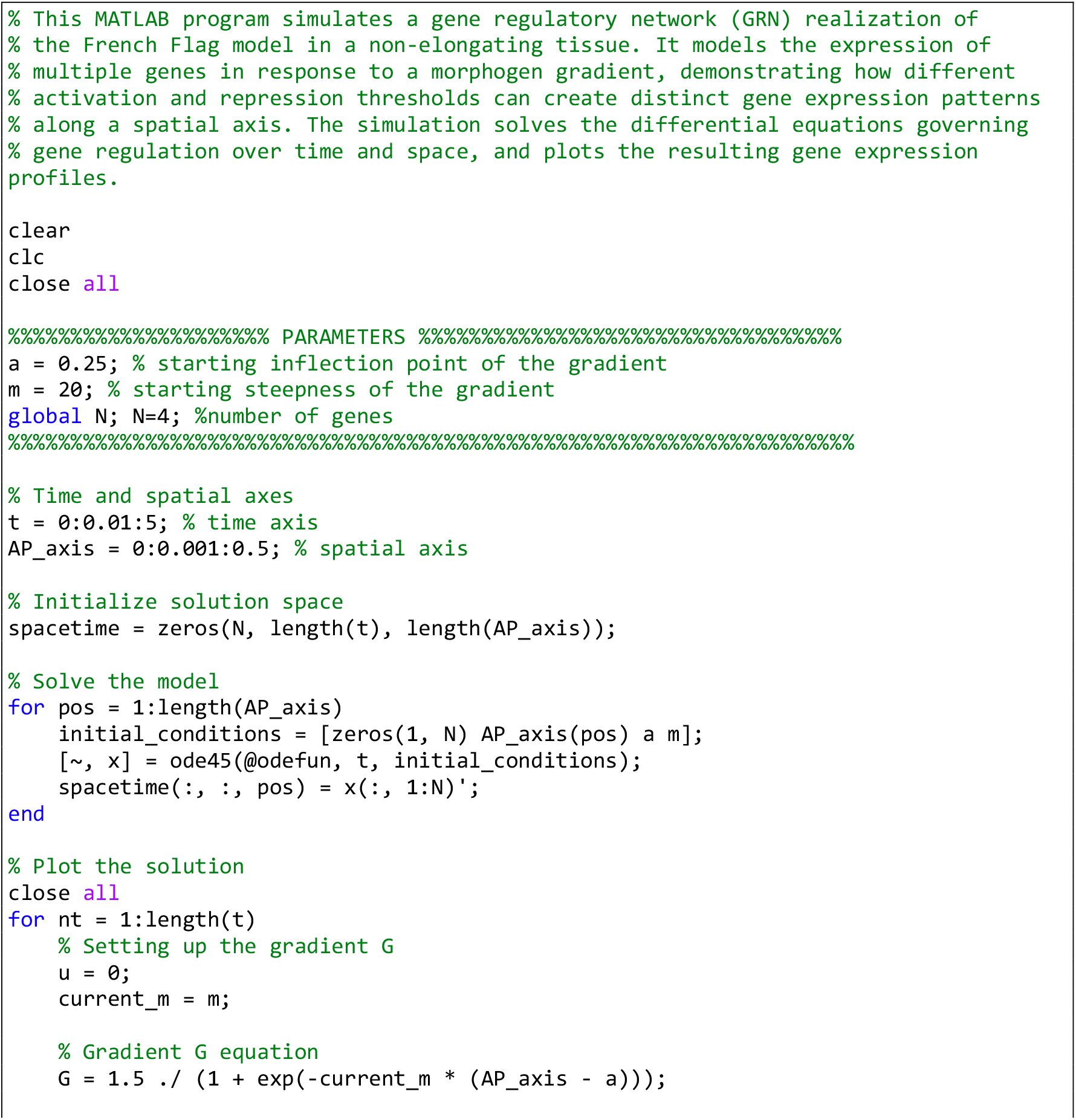
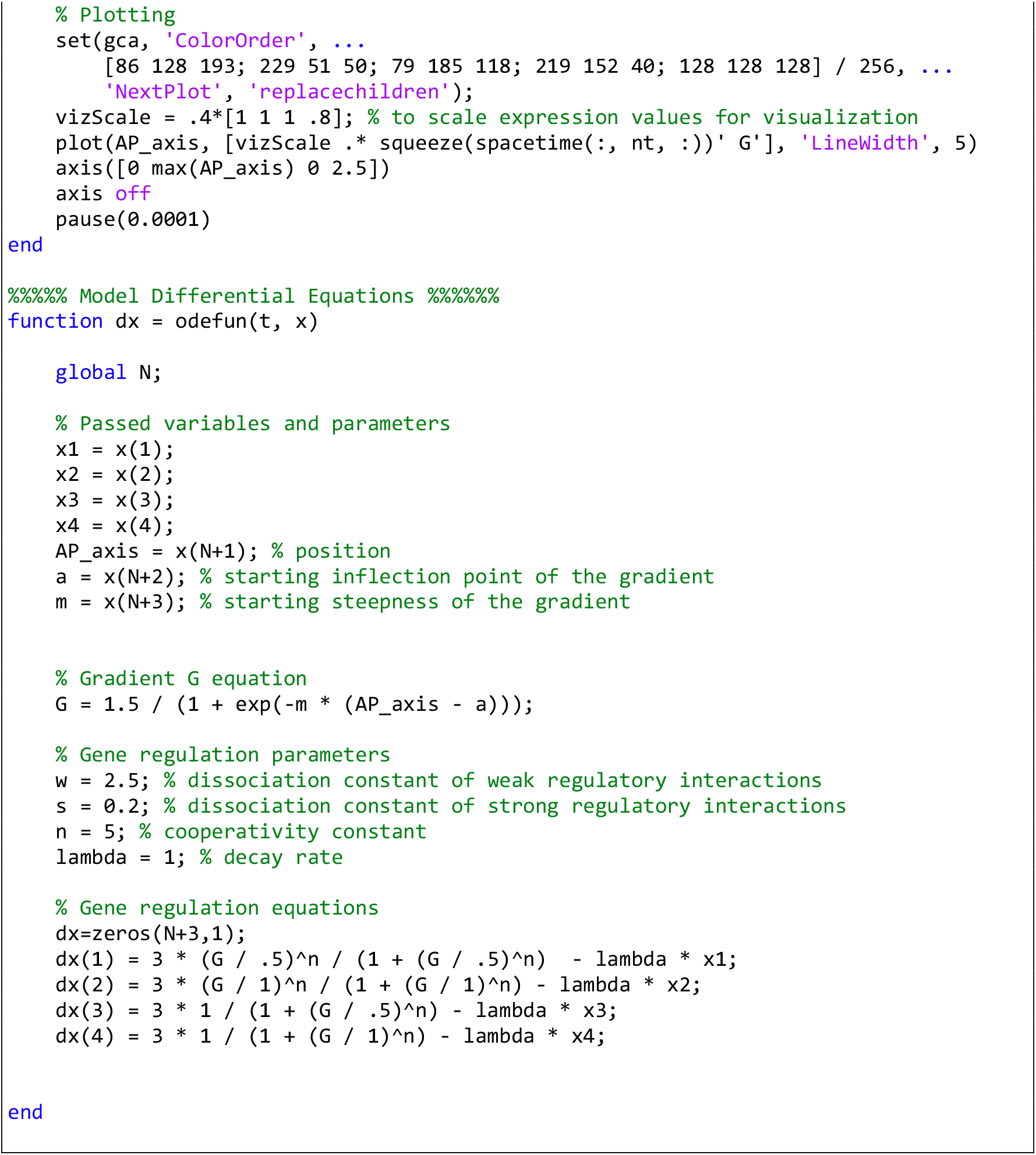

**Computer Simulation 20**. Defining Gene Expression Boundaries Using Two Morphogen Gradients.

The results of this computer simulation are shown in Fig. 5D’.

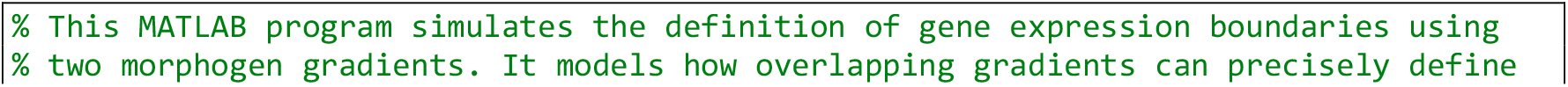
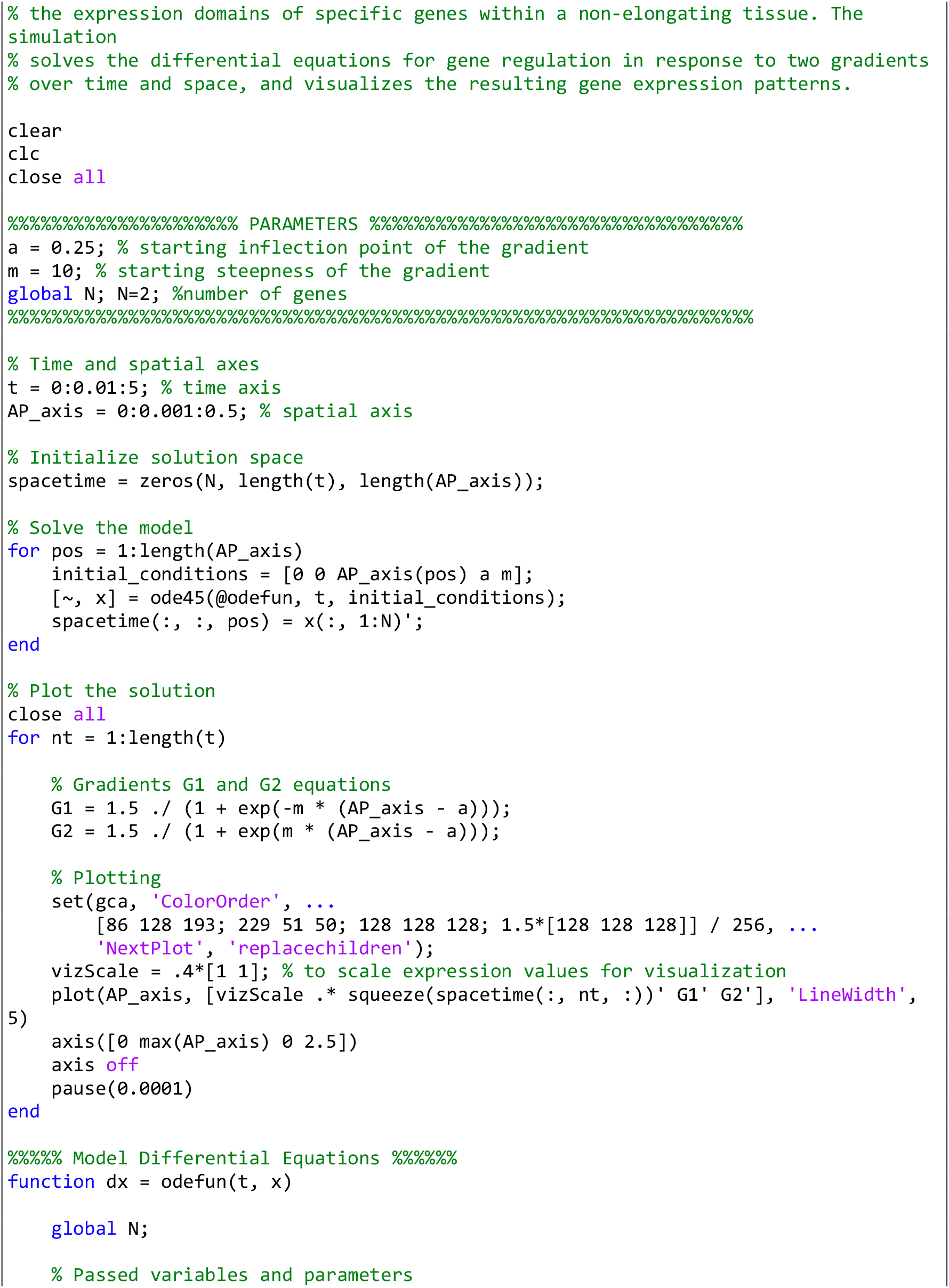
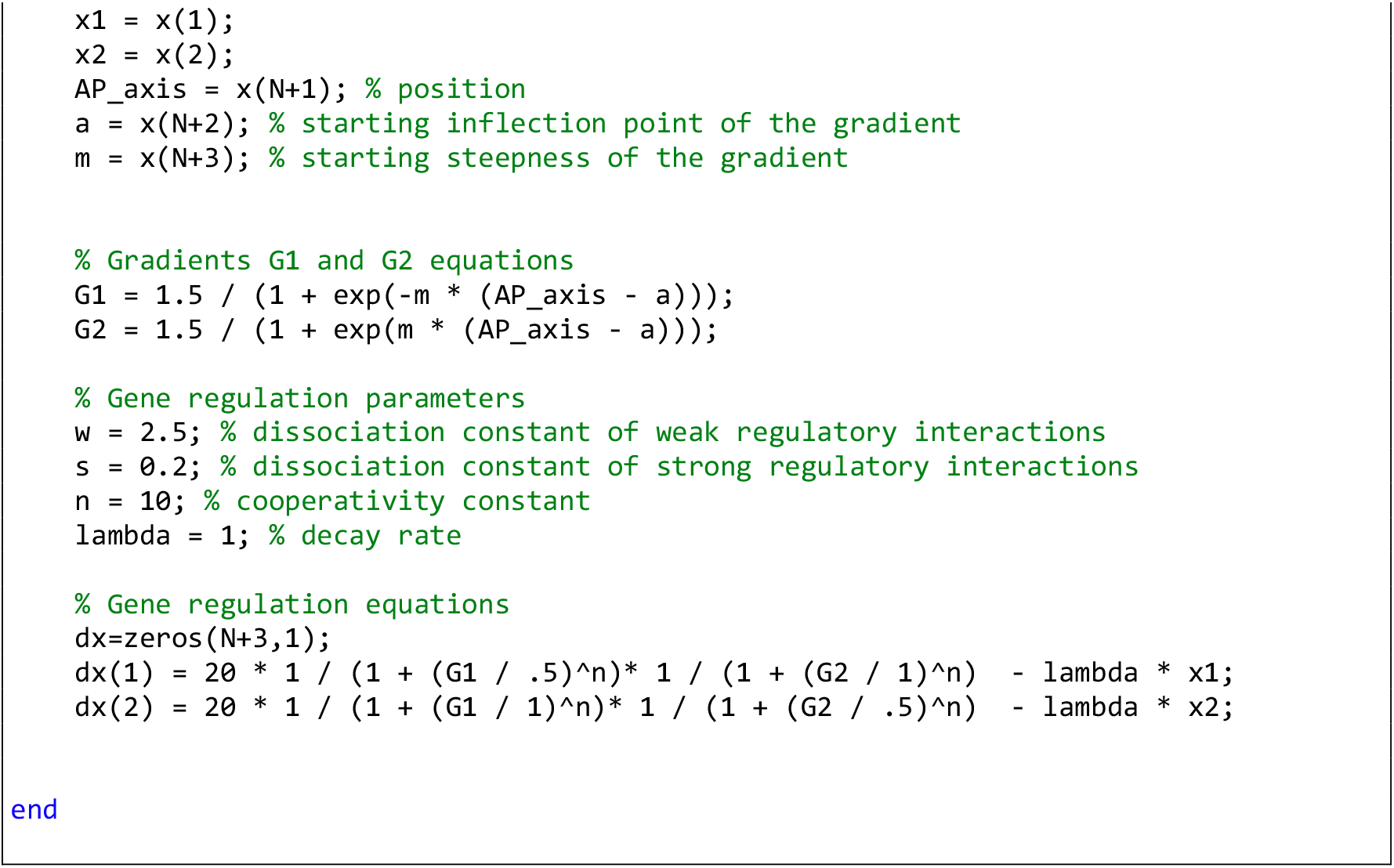

**Computer Simulation 21**. Gene Regulatory Network Configuration Involving Cross-Regulatory Interactions.

The results of this computer simulation are shown in Fig. 5E’.

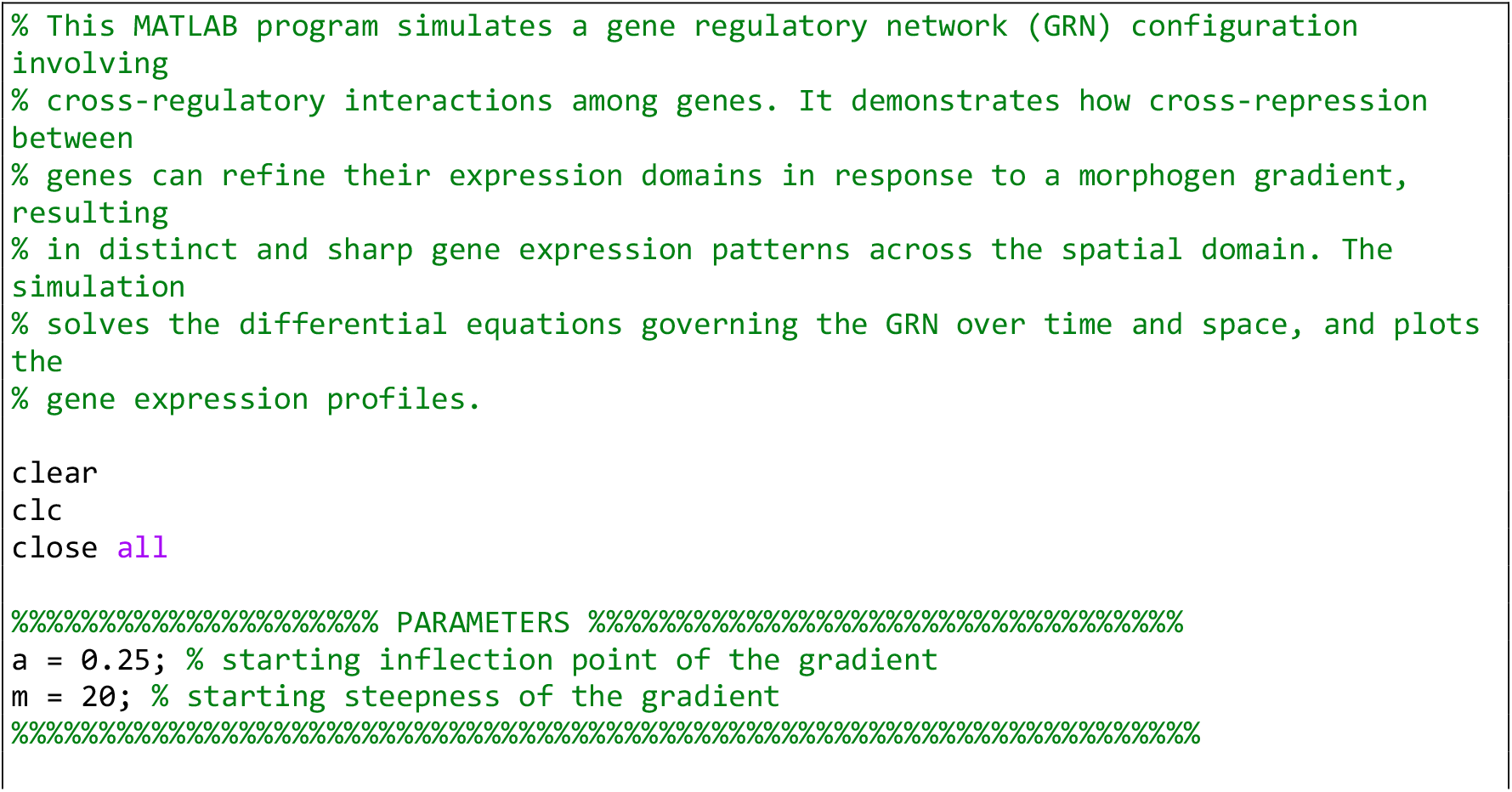
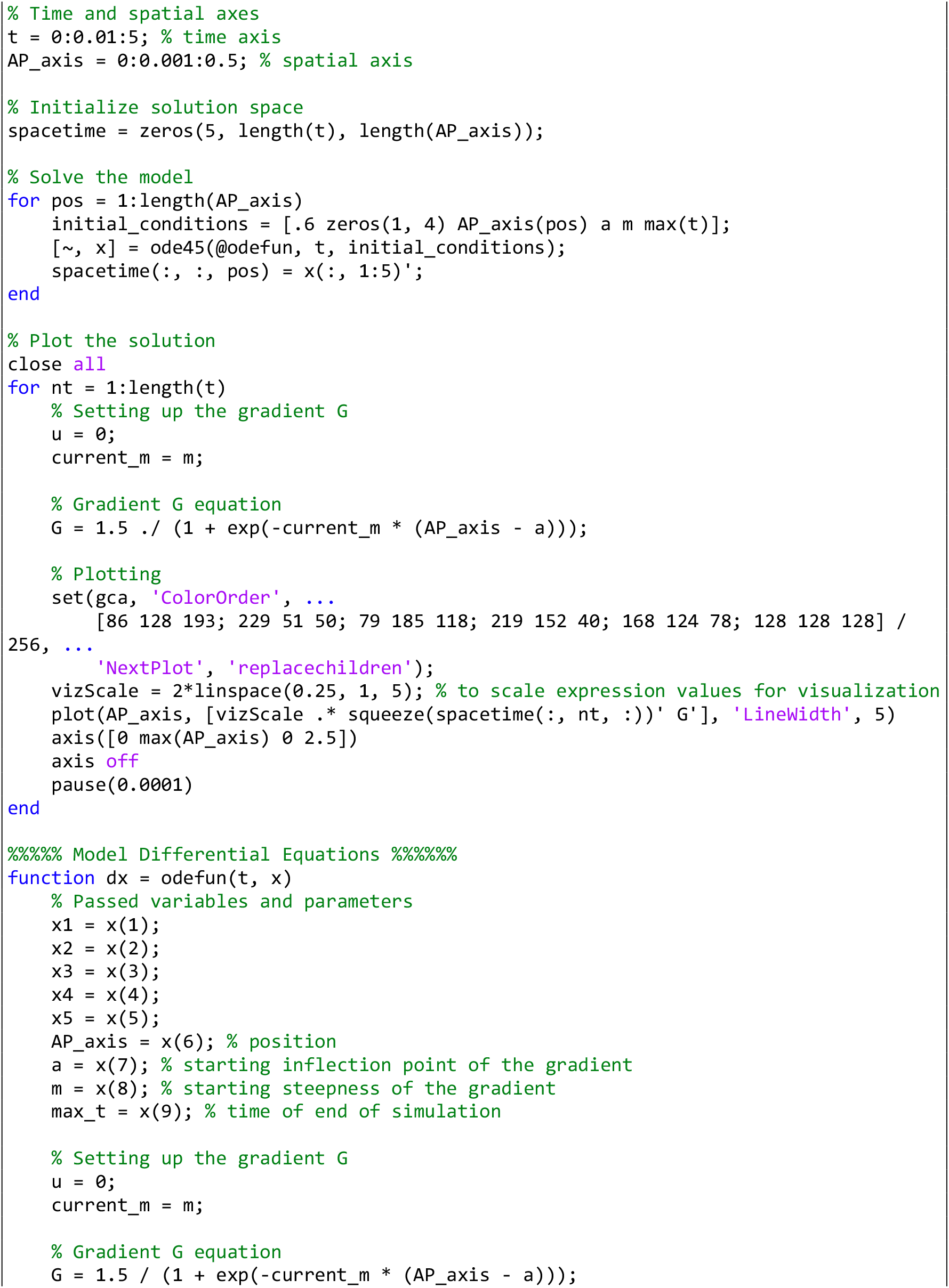
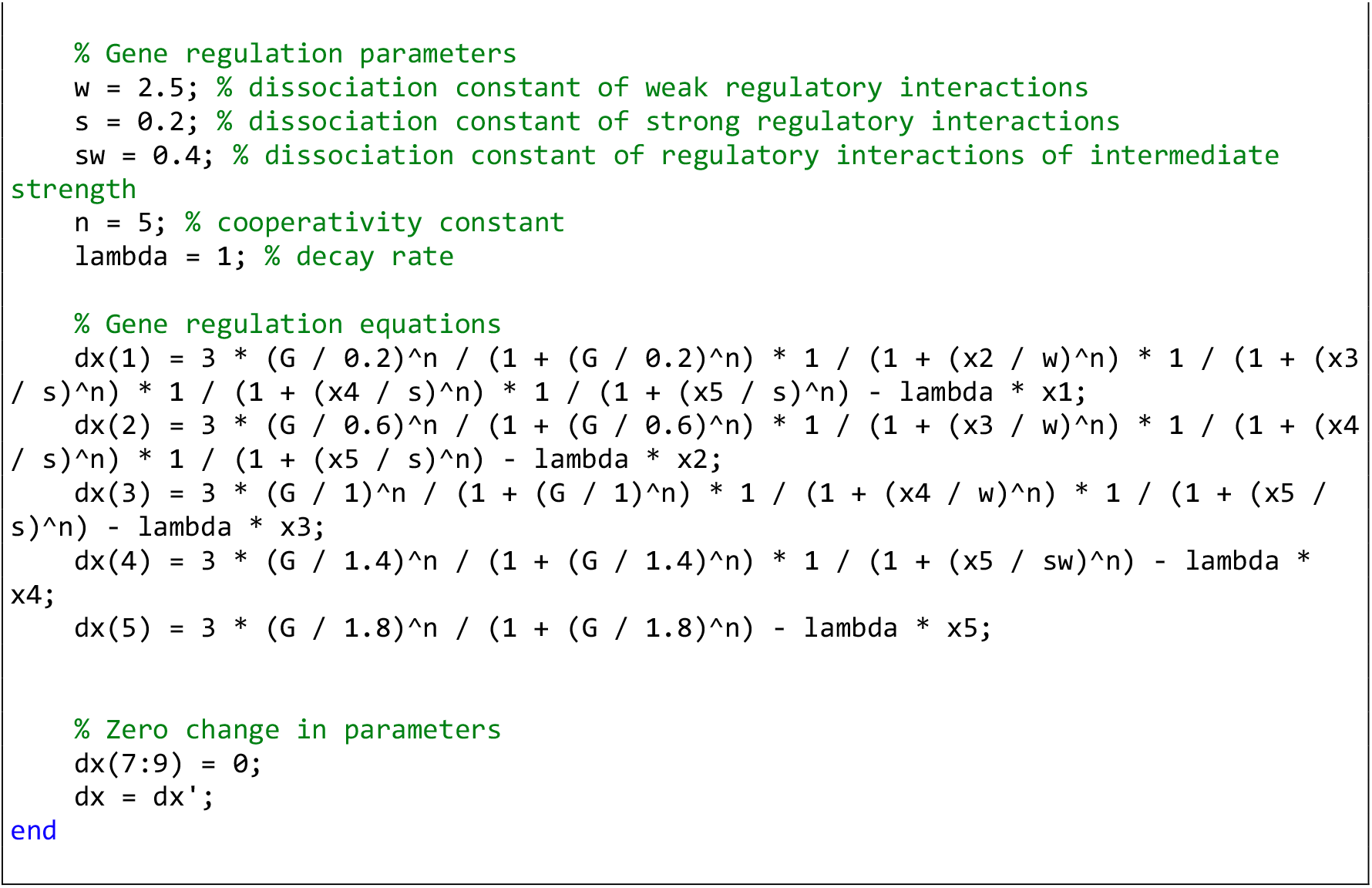

**Computer Simulation 22**. Additive Effect of Separate Enhancers on Gene Expression to Form Periodic Patterns.

The results of this computer simulation are shown in Fig. 5H’.

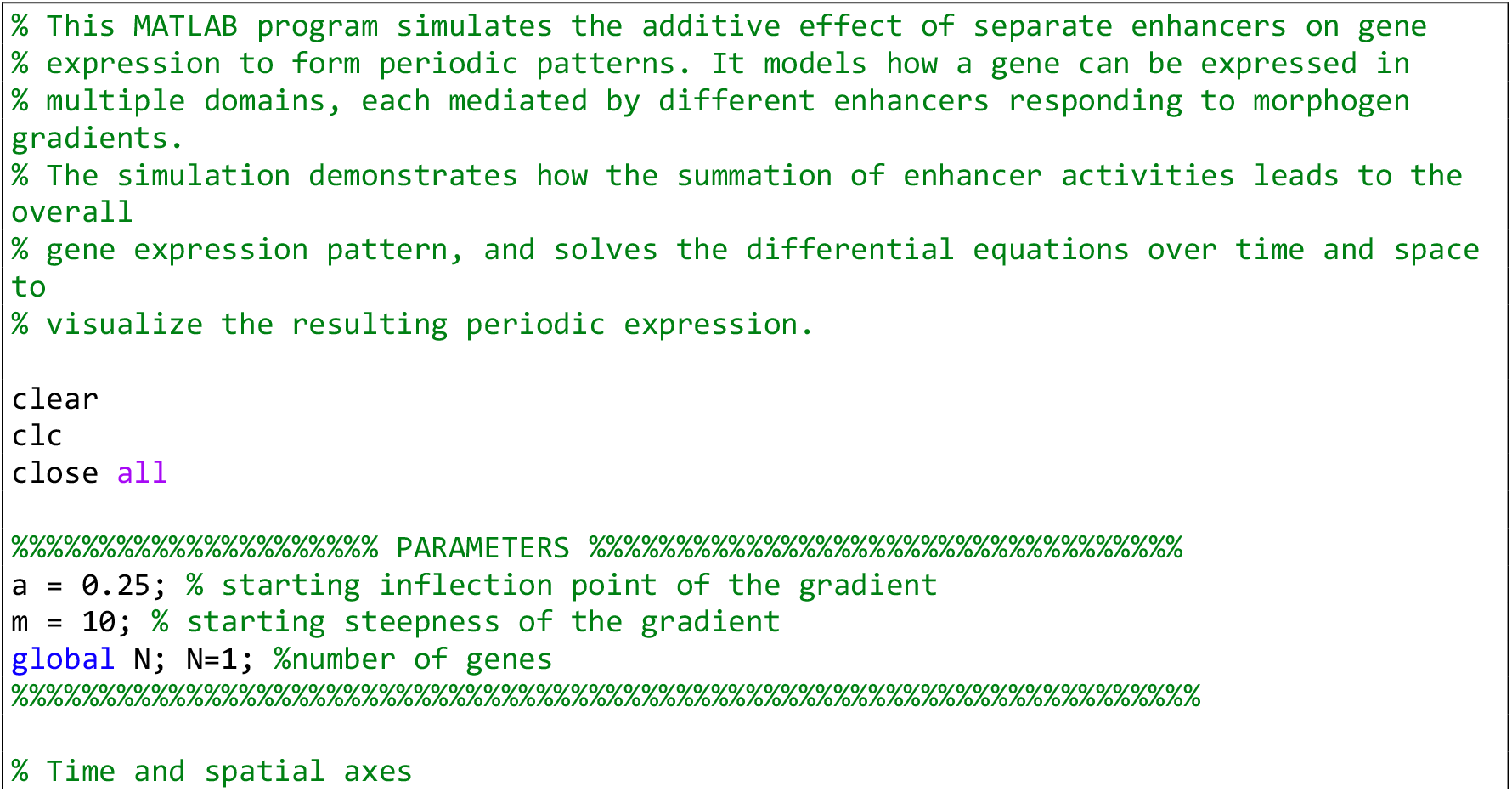
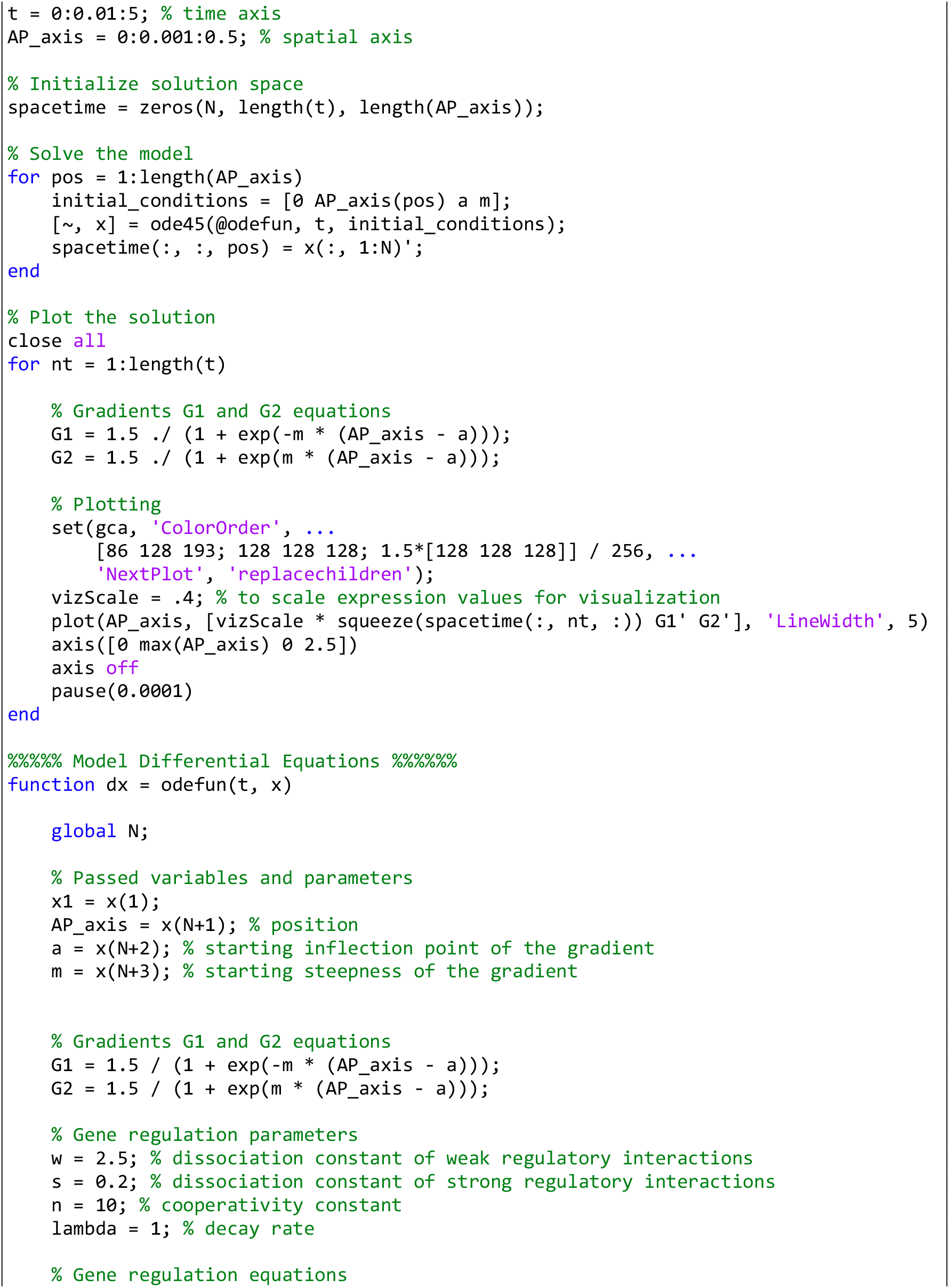
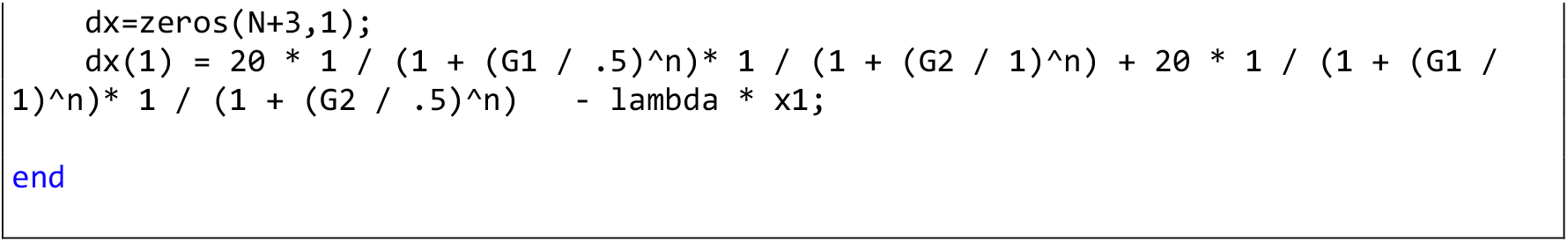

**Computer Simulation 23**. Realization of the Speed Regulation Model by Modulating Transcription and Decay Rates.

The results of this computer simulation are shown in Fig. 6B’.

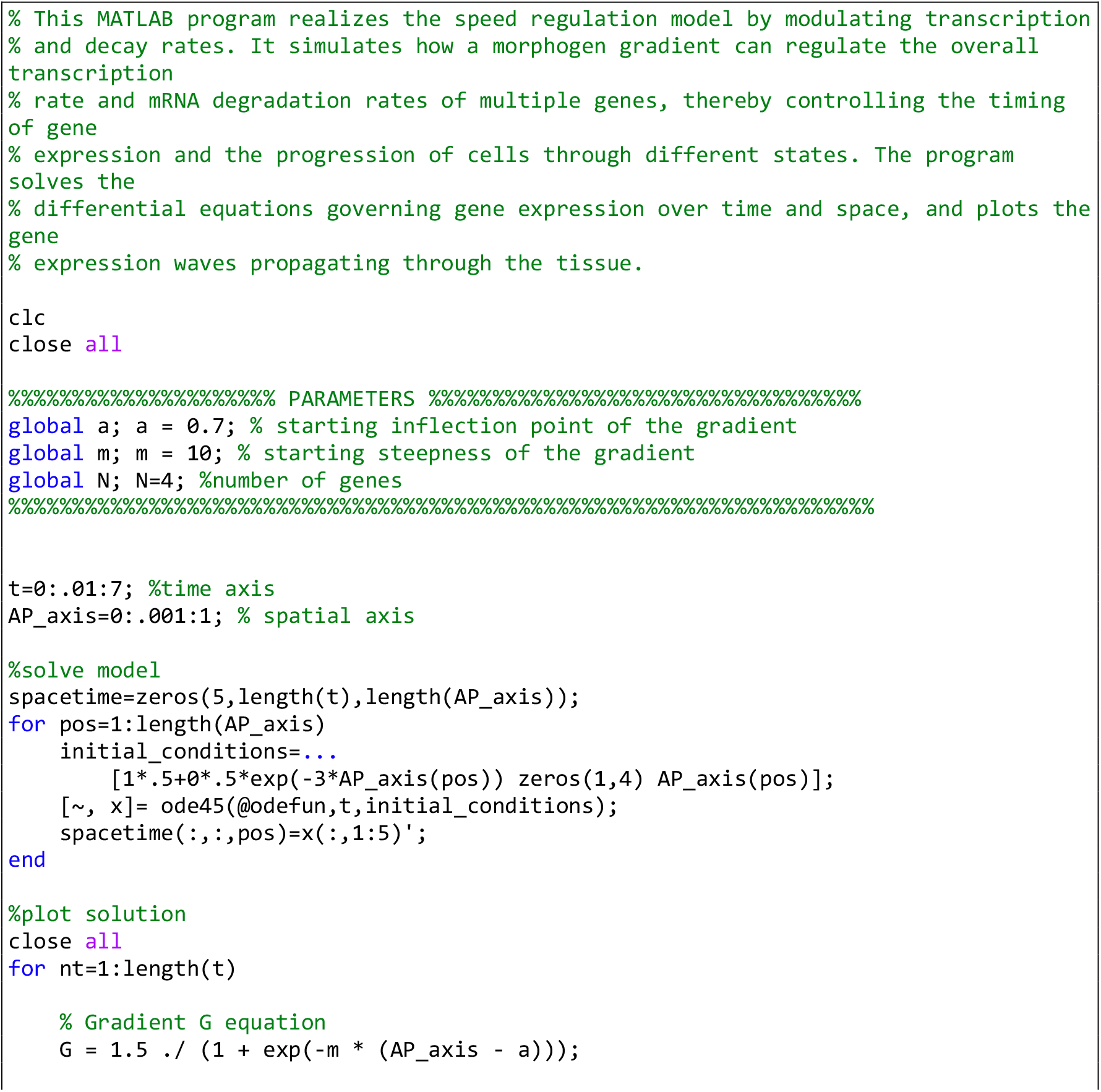
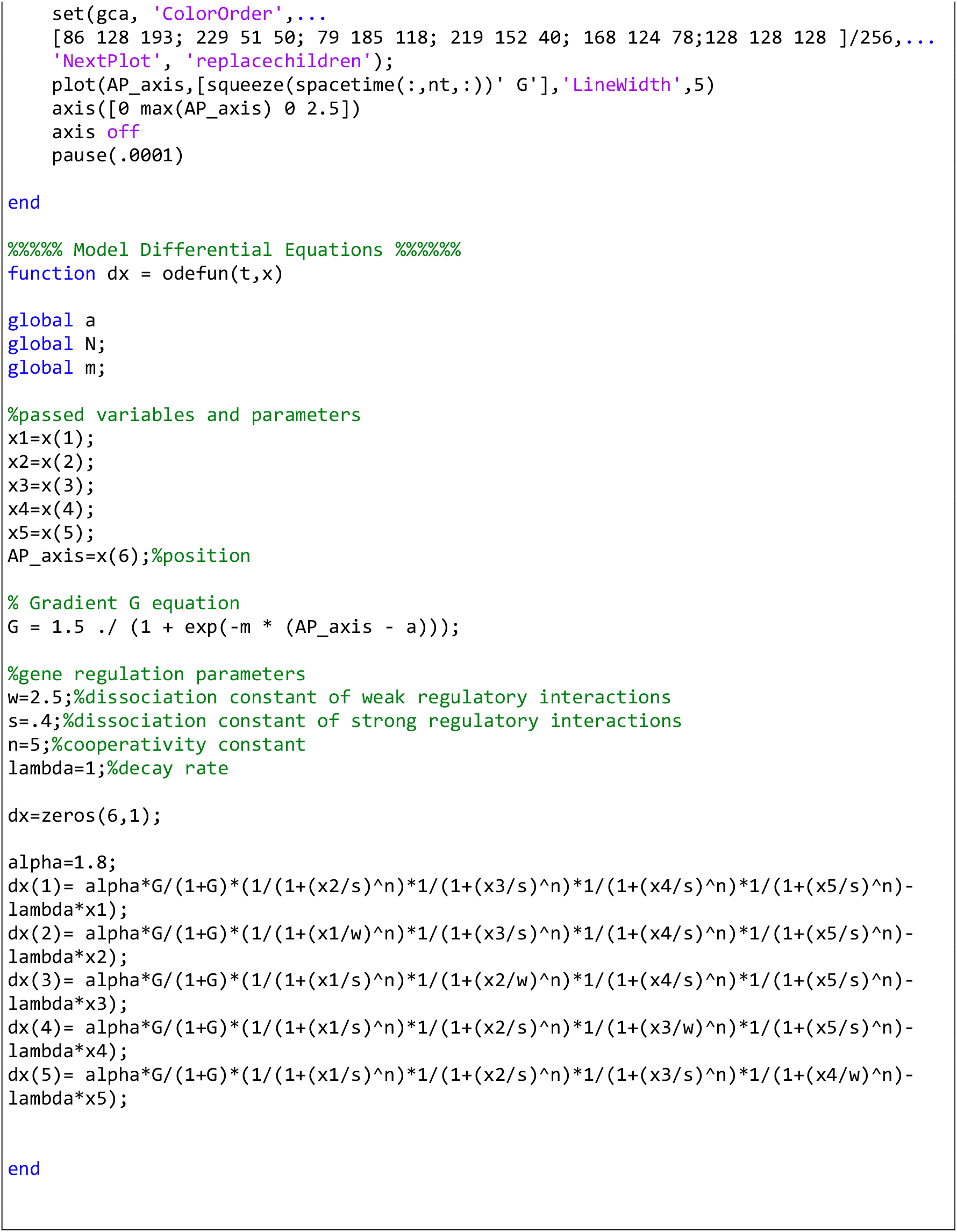

**Computer Simulation 24**. Enhancer Switching Model as a Gene Regulatory Network Realization of the Speed Regulation Model.

The results of this computer simulation are shown in Fig. 6C’.

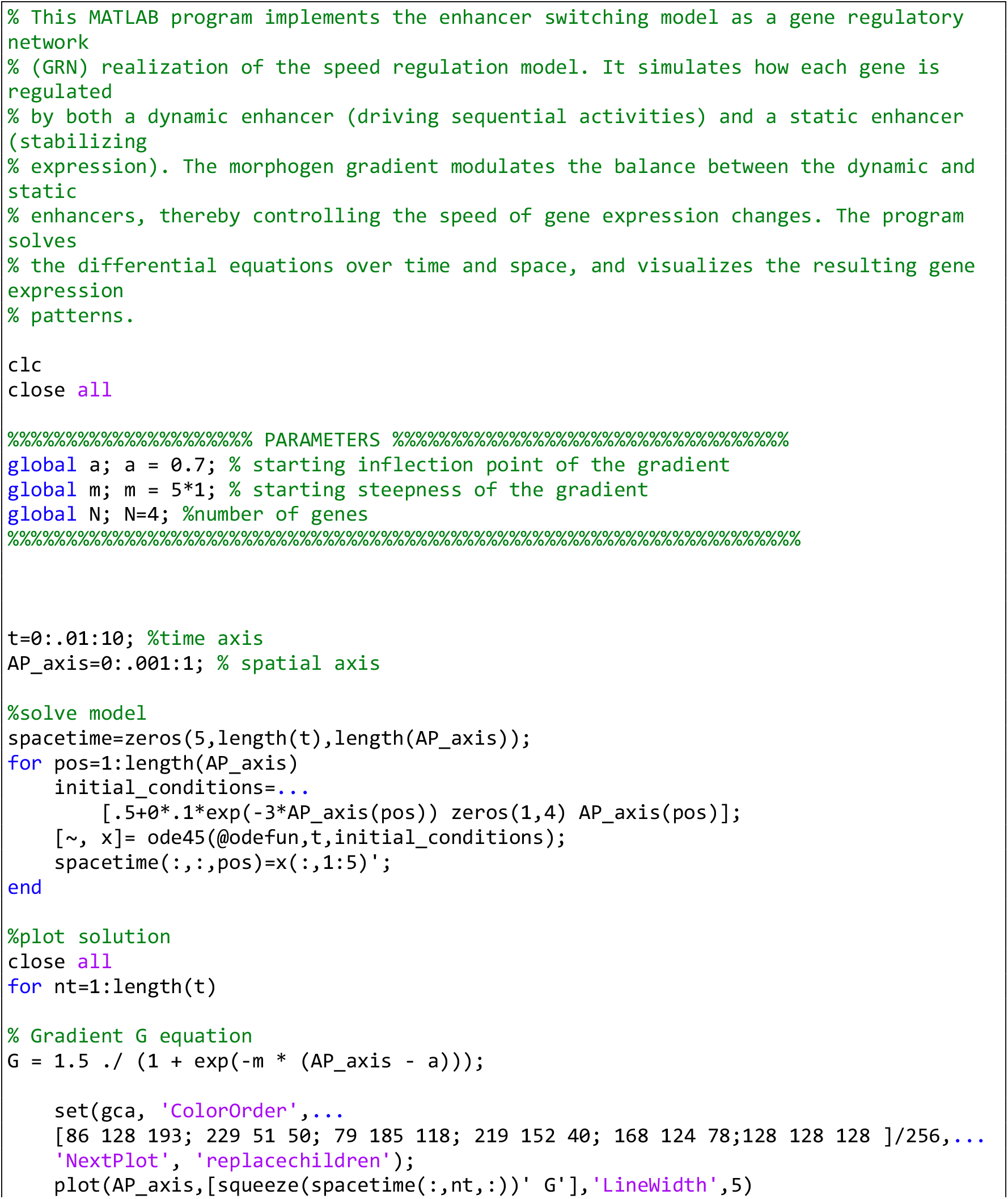
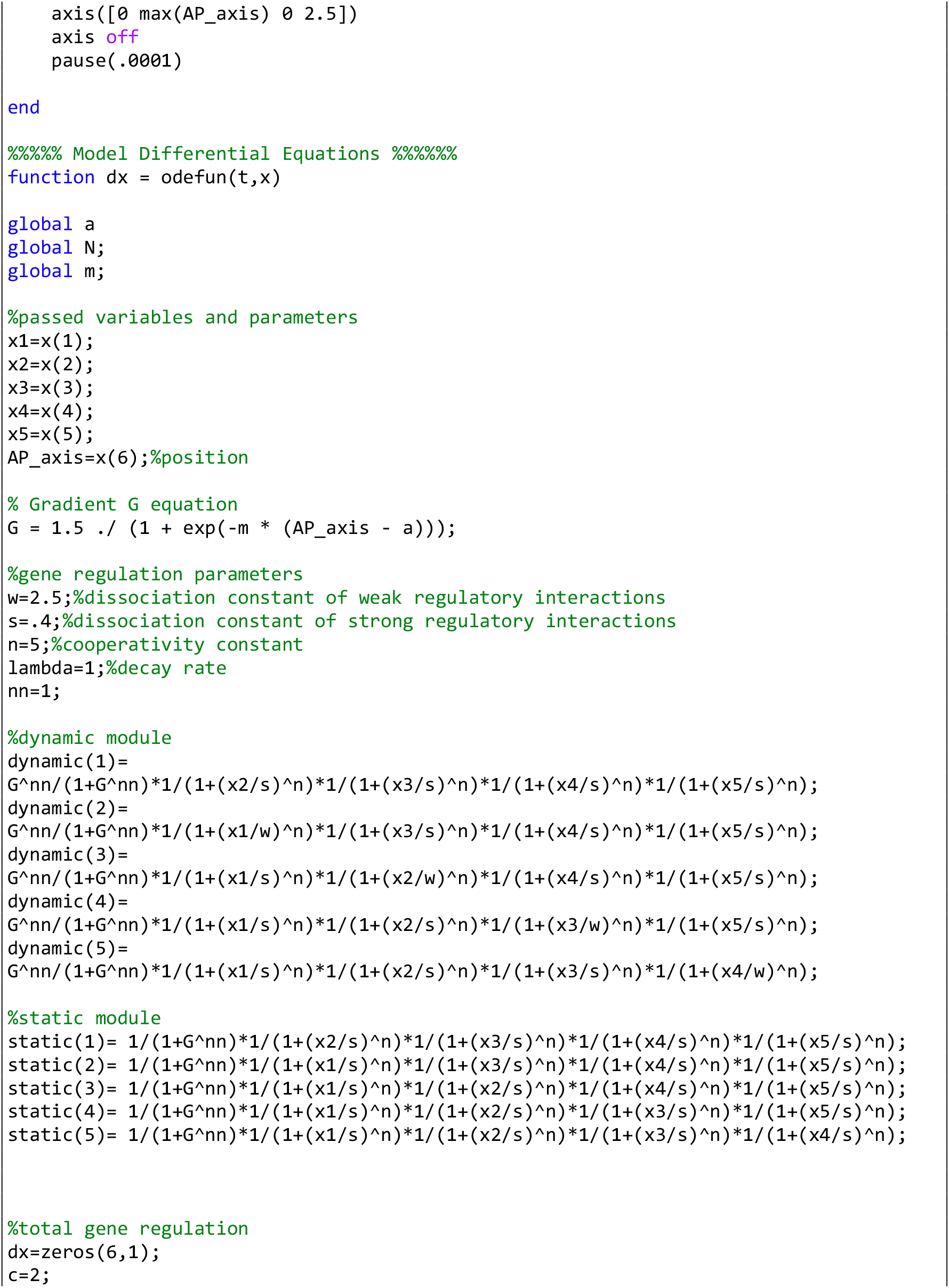
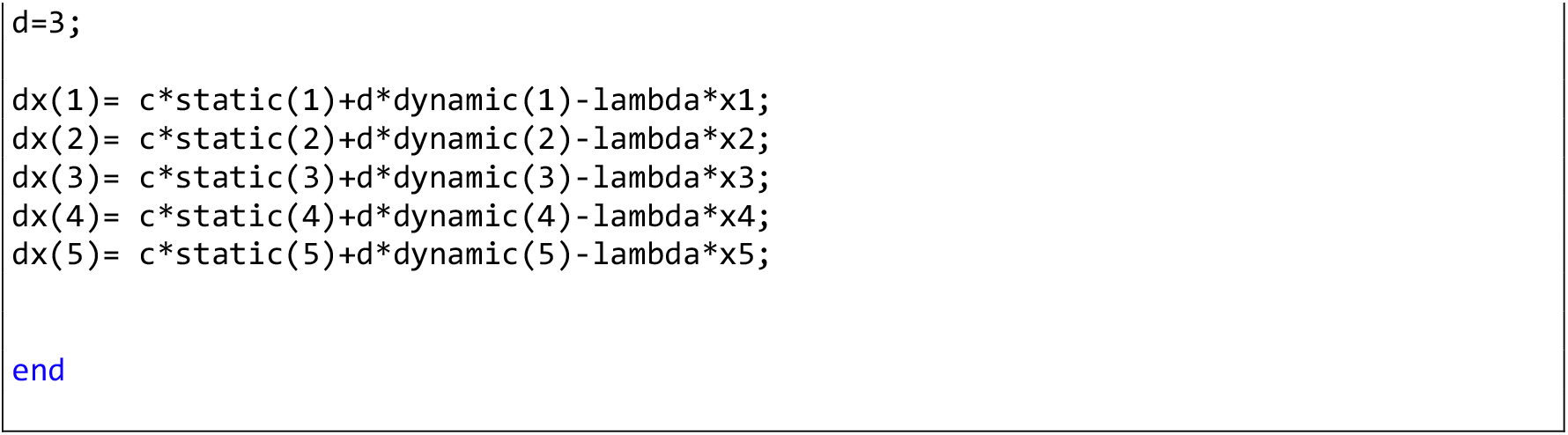

**Computer Simulation 25**. Impact of an Overactive Enhancer in the Additive Integration Model.

The results of this computer simulation are shown in Fig. 7A’ (WT; parameter mut_flag=0 in the program) and 7A’’ (overactive dynamic enhancer of Gene 3; parameter mut_flag=1 in the program).

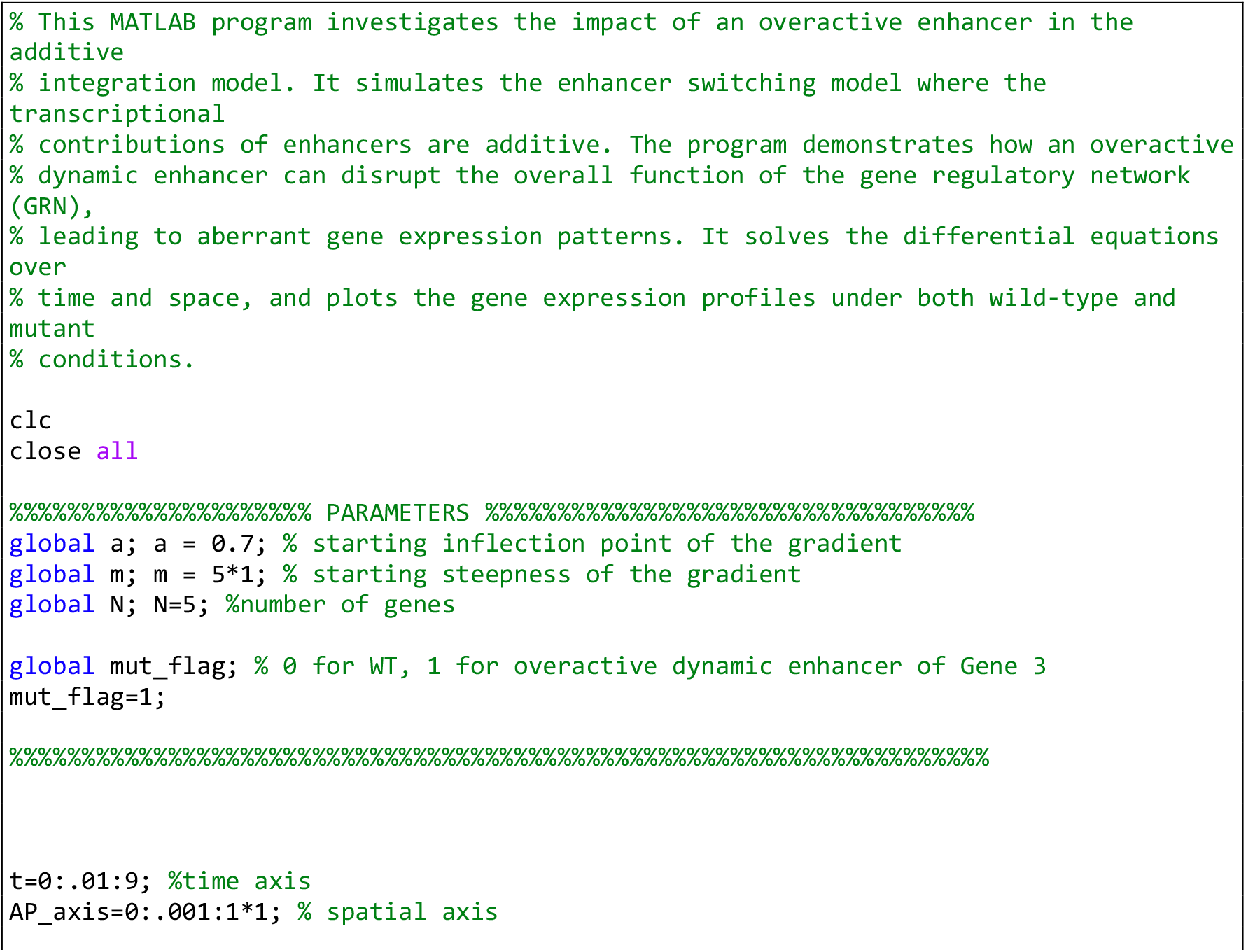
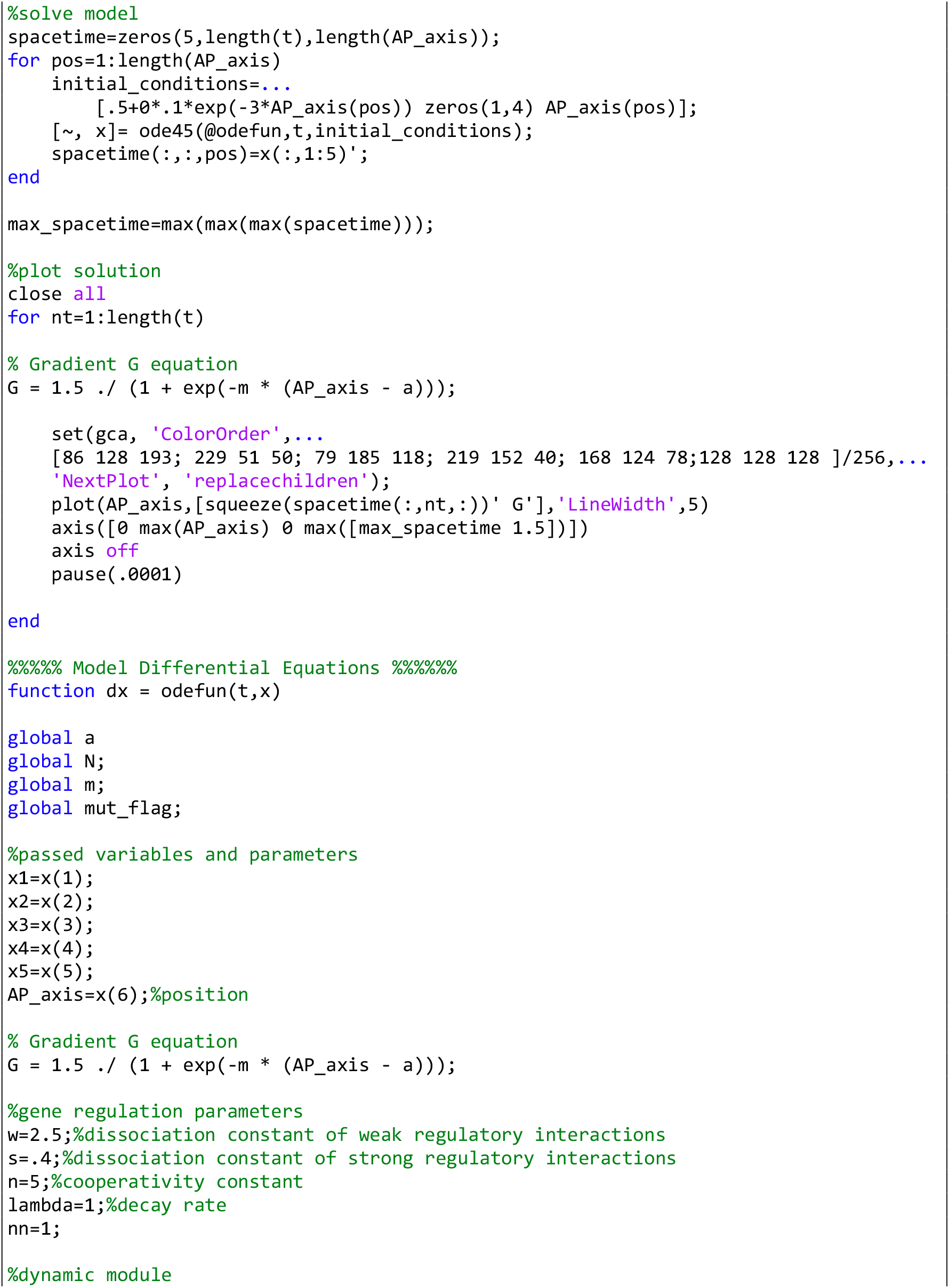
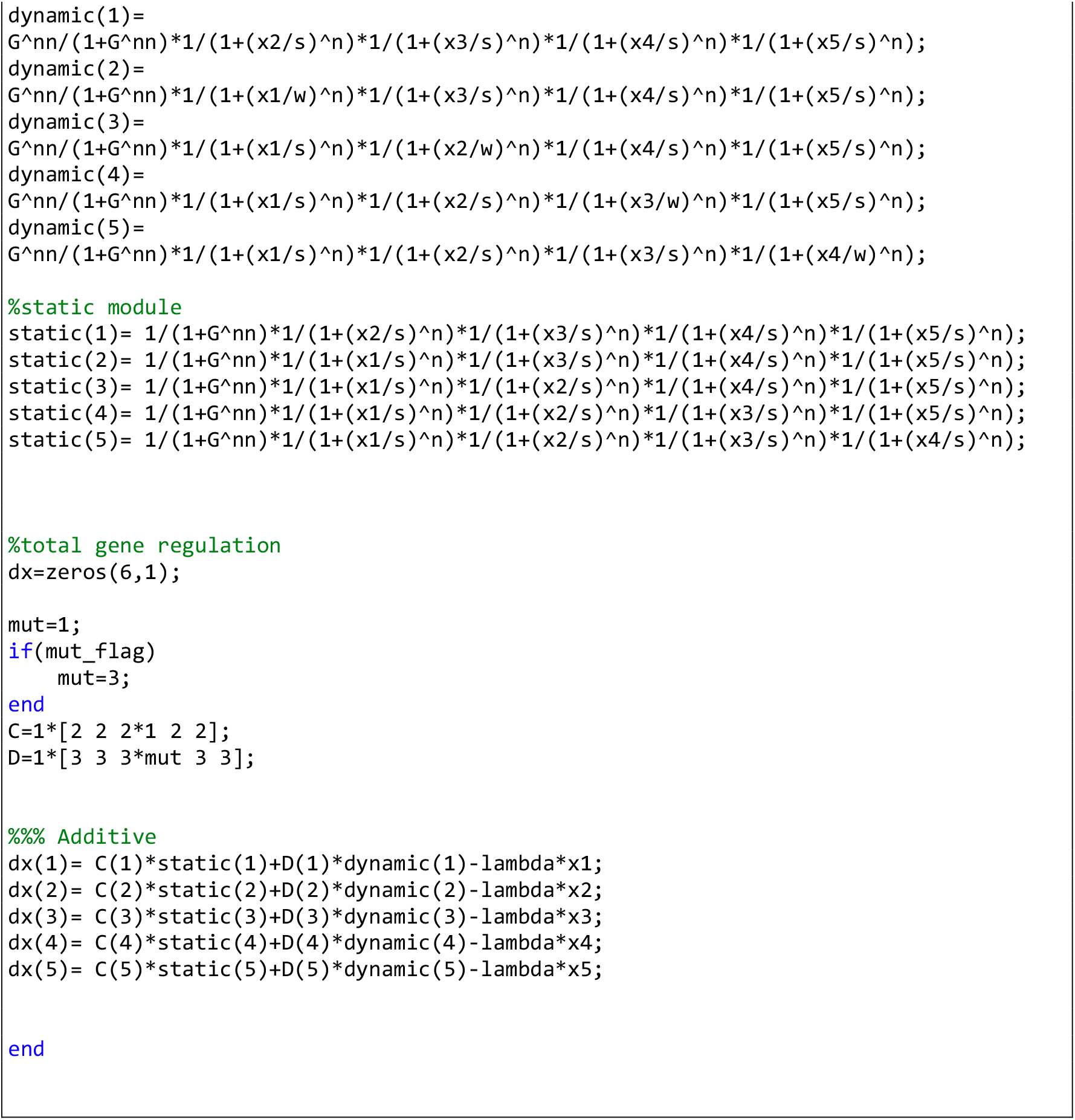

**Computer Simulation 26**. Mitigation of Overactive Enhancer Effects Using Enhancer Competition Integration.

The results of this computer simulation are shown in Fig. 7B’ (WT; parameter mut_flag=0 in the program) and 7B’’ (overactive dynamic enhancer of Gene 3; parameter mut_flag=1 in the program).

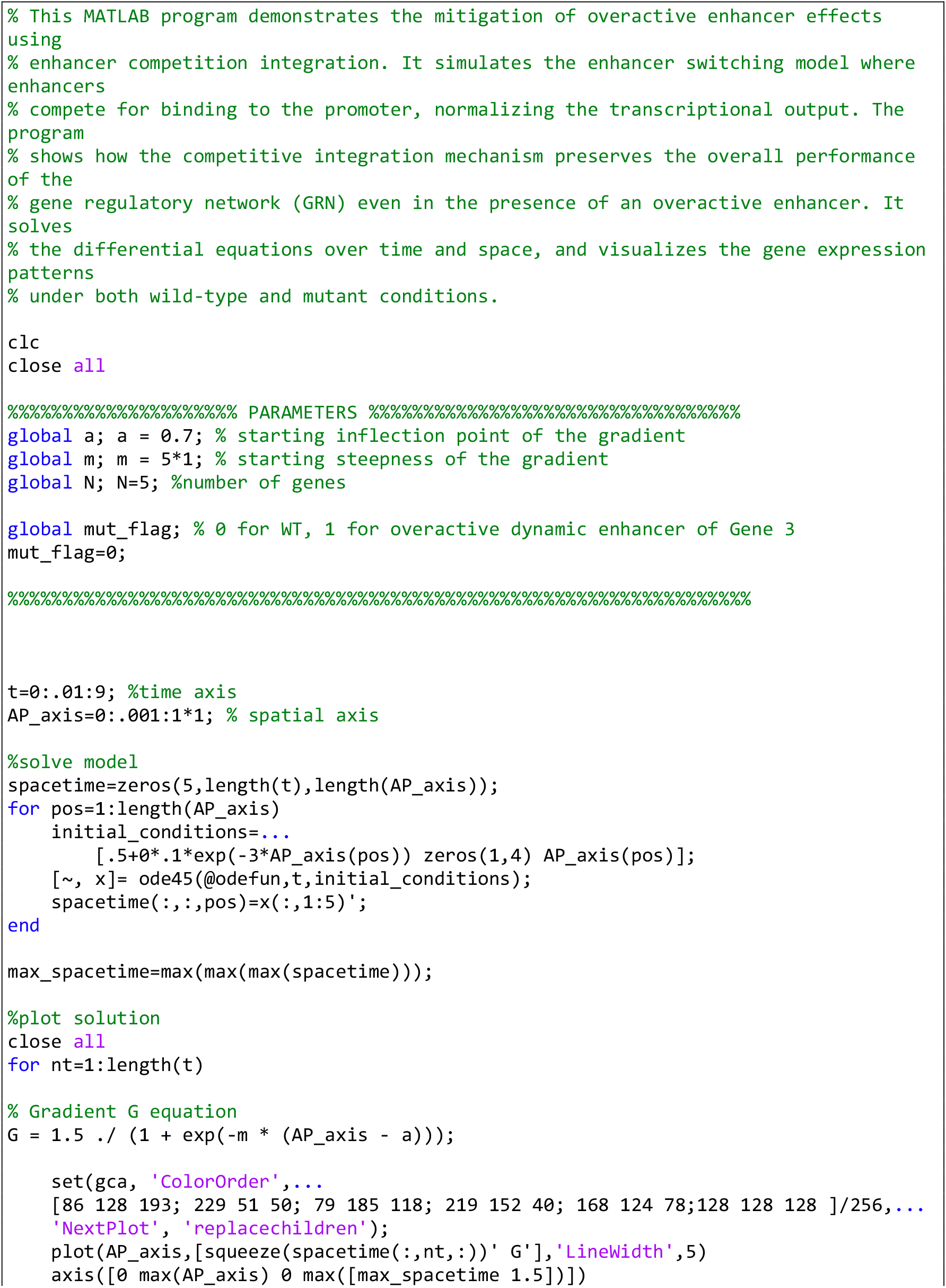
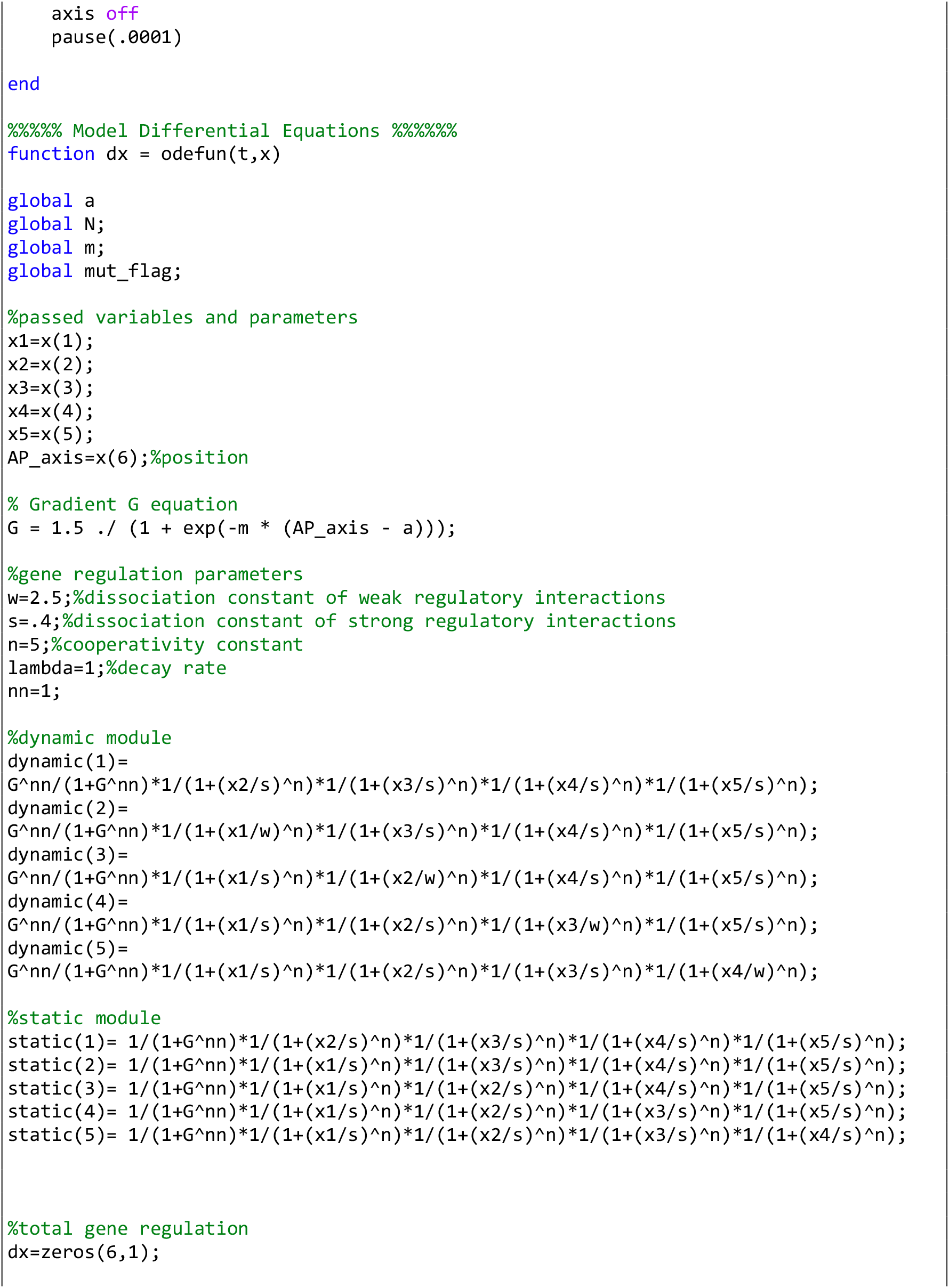
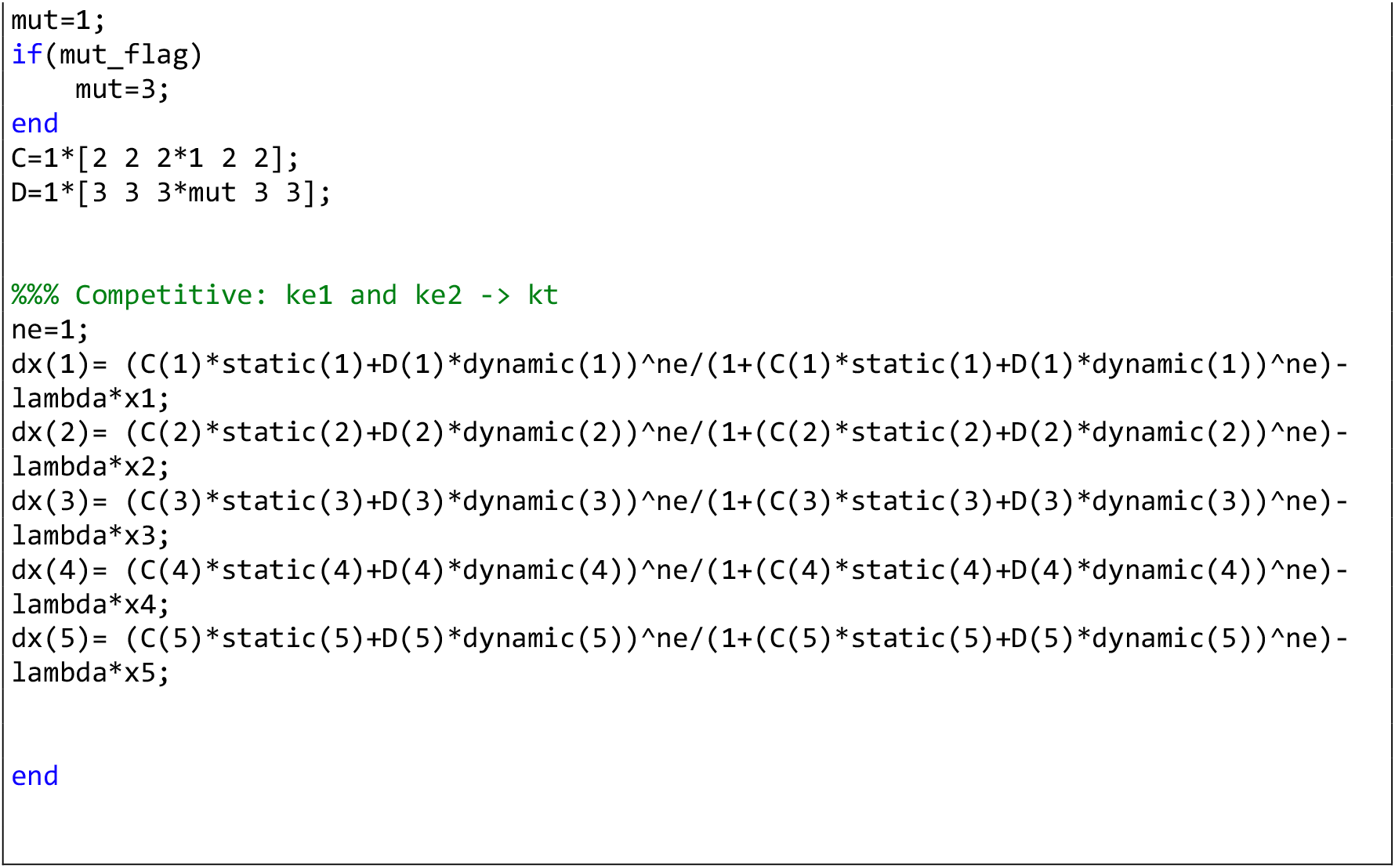

## Notes

### Competing Interest Statement

The authors have declared no competing interest.

### Summary of Updates

- Codes for computer simulations added - General Editing

## References

1. Negrete J, Oates AC. Towards a physical understanding of developmental patterning. Nat Rev Genet. 2021 Aug;22(8):518–31.

2. Wolpert L. Positional information and the spatial pattern of cellular differentiation. J Theor Biol. 1969 Oct;25(1):1–47.

3. Green JBA, Sharpe J. Positional information and reaction-diffusion: two big ideas in developmental biology combine. Development. 2015 Apr 1;142(7):1203–11.

4. Müller P, El-Sherif E. A systems-level view of pattern formation mechanisms in development. Dev Biol. 2020 Apr 1;460(1):1.

5. Wolpert L. Positional information revisited. Development. 1989;107 Suppl:3–12.

6. Landge AN, Jordan BM, Diego X, Müller P. Pattern formation mechanisms of self-organizing reaction-diffusion systems. Dev Biol. 2020 Apr 1;460(1):2–11.

7. Meinhardt H. Models of biological pattern formation and their application to the early development of drosophila. In: Mosekilde E, Mosekilde L, editors. Complexity, chaos, and biological evolution. New York, NY: Sprimger US; 1991. p. 303–22.

8. Roth S. Mathematics and biology: a Kantian view on the history of pattern formation theory. Dev Genes Evol. 2011 Dec 1;221(5–6):255–79.

9. Briscoe J, Small S. Morphogen rules: design principles of gradient-mediated embryo patterning. Development. 2015 Dec 1;142(23):3996–4009.

10. Spitz F, Furlong EEM. Transcription factors: from enhancer binding to developmental control. Nat Rev Genet. 2012 Sep;13(9):613–26.

11. Anderson PW. More is different. Science. 1972 Aug 4;177(4047):393–6.

12. Davidson EH. Emerging properties of animal gene regulatory networks. Nature. 2010 Dec 16;468(7326):911–20.

13. Levine M, Davidson EH. Gene regulatory networks for development. Proc Natl Acad Sci USA. 2005 Apr 5;102(14):4936–42.

14. Fuda NJ, Ardehali MB, Lis JT. Defining mechanisms that regulate RNA polymerase II transcription in vivo. Nature. 2009 Sep 10;461(7261):186–92.

15. Alon U. An Introduction to Systems Biology: Design Principles of Biological Circuits. illustrated ed. CRC Press; 2006.

16. Alon U. Network motifs: theory and experimental approaches. Nat Rev Genet. 2007 Jun;8(6):450–61.

17. Cramer P. Organization and regulation of gene transcription. Nature. 2019 Sep;573(7772):45– 54.

18. Furlong EEM, Levine M. Developmental enhancers and chromosome topology. Science. 2018 Sep 28;361(6409):1341–5.

19. Fukaya T, Lim B, Levine M. Enhancer control of transcriptional bursting. Cell. 2016 Jul 14;166(2):358–68.

20. Kalir S, Mangan S, Alon U. A coherent feed-forward loop with a SUM input function prolongs flagella expression in Escherichia coli. Mol Syst Biol. 2005 Mar 29;1:2005.0006.

21. Bintu L, Buchler NE, Garcia HG, Gerland U, Hwa T, Kondev J, et al. Transcriptional regulation by the numbers: models. Curr Opin Genet Dev. 2005 Apr;15(2):116–24.

22. Bintu L, Buchler NE, Garcia HG, Gerland U, Hwa T, Kondev J, et al. Transcriptional regulation the numbers: applications. Curr Opin Genet Dev. 2005 Apr;15(2):125–35.

23. Shea MA, Ackers GK. The OR control system of bacteriophage lambda. A physical-chemical model for gene regulation. J Mol Biol. 1985 Jan 20;181(2):211–30.

24. Kim YJ, Rhee K, Liu J, Jeammet S, Turner MA, Small SJ, et al. Predictive modeling reveals that higher-order cooperativity drives transcriptional repression in a synthetic developmental enhancer. eLife. 2022 Dec 12;11.

25. Vincent BJ, Estrada J, DePace AH. The appeasement of Doug: a synthetic approach to enhancer biology. Integr Biol (Camb). 2016 Apr 18;8(4):475–84.

26. Samee MAH, Lydiard-Martin T, Biette KM, Vincent BJ, Bragdon MD, Eckenrode KB, et al. Quantitative Measurement and Thermodynamic Modeling of Fused Enhancers Support a Two-Tiered Mechanism for Interpreting Regulatory DNA. Cell Rep. 2017 Oct 3;21(1):236–45.

27. Park J, Estrada J, Johnson G, Vincent BJ, Ricci-Tam C, Bragdon MD, et al. Dissecting the sharp response of a canonical developmental enhancer reveals multiple sources of cooperativity. eLife. 2019 Jun 21;8.

28. Chen J, Boyaci H, Campbell EA. Diverse and unified mechanisms of transcription initiation in bacteria. Nat Rev Microbiol. 2021 Feb;19(2):95–109.

29. Yokoshi M, Fukaya T. Dynamics of transcriptional enhancers and chromosome topology in gene regulation. Dev Growth Differ. 2019 Jun;61(5):343–52.

30. Lammers NC, Kim YJ, Zhao J, Garcia HG. A matter of time: Using dynamics and theory to uncover mechanisms of transcriptional bursting. Curr Opin Cell Biol. 2020 Dec;67:147–57.

31. Adelman K, Lis JT. Promoter-proximal pausing of RNA polymerase II: emerging roles in metazoans. Nat Rev Genet. 2012 Oct 1;13(10):720–31.

32. Gaertner B, Zeitlinger J. RNA polymerase II pausing during development. Development. 2014 Mar;141(6):1179–83.

33. Green MR. Eukaryotic transcription activation: right on target. Mol Cell. 2005 May 13;18(4):399–402.

34. Nechaev S, Adelman K. Pol II waiting in the starting gates: Regulating the transition from transcription initiation into productive elongation. Biochim Biophys Acta. 2011 Jan;1809(1):34–45.

35. Scholes C, DePace AH, Sánchez Á. Combinatorial Gene Regulation through Kinetic Control of the Transcription Cycle. Cell Syst. 2017 Jan 25;4(1):97-108.e9.

36. Sherman MS, Cohen BA. Thermodynamic state ensemble models of cis-regulation. PLoS Comput Biol. 2012 Mar 29;8(3):e1002407.

37. Longabaugh WJR, Davidson EH, Bolouri H. Computational representation of developmental genetic regulatory networks. Dev Biol. 2005 Jul 1;283(1):1–16.

38. Elowitz MB, Leibler S. A synthetic oscillatory network of transcriptional regulators. Nature. 2000 Jan 20;403(6767):335–8.

39. Averbukh I, Lai S-L, Doe CQ, Barkai N. A repressor-decay timer for robust temporal patterning in embryonic Drosophila neuroblast lineages. eLife. 2018 Dec 10;7.

40. Tufcea DE, François P. Critical Timing without a Timer for Embryonic Development. Biophys J. 2015 Oct 20;109(8):1724–34.

41. Clark E, Peel AD, Akam M. Arthropod segmentation. Development. 2019 Sep 25;146(18).

42. Diaz-Cuadros M, Pourquié O, El-Sherif E. Patterning with clocks and genetic cascades: Segmentation and regionalization of vertebrate versus insect body plans. PLoS Genet. 2021 Oct 14;17(10):e1009812.

43. Moussian B, Roth S. Dorsoventral axis formation in the Drosophila embryo--shaping and transducing a morphogen gradient. Curr Biol. 2005 Nov 8;15(21):R887–99.

44. Zhu X, Rudolf H, Healey L, François P, Brown SJ, Klingler M, et al. Speed regulation of genetic cascades allows for evolvability in the body plan specification of insects. Proc Natl Acad Sci USA. 2017 Sep 25;114(41).

45. Jaeger J, Surkova S, Blagov M, Janssens H, Kosman D, Kozlov KN, et al. Dynamic control of positional information in the early Drosophila embryo. Nature. 2004 Jul 15;430(6997):368–71.

46. Setty Y, Mayo AE, Surette MG, Alon U. Detailed map of a cis-regulatory input function. Proc Natl Acad Sci USA. 2003 Jun 24;100(13):7702–7.

47. Mayo AE, Setty Y, Shavit S, Zaslaver A, Alon U. Plasticity of the cis-regulatory input function of a gene. PLoS Biol. 2006 Apr;4(4):e45.

48. Yuh CH, Bolouri H, Davidson EH. Genomic cis-regulatory logic: experimental and computational analysis of a sea urchin gene. Science. 1998 Mar 20;279(5358):1896–902.

49. Hanna-Rose W, Licht JD, Hansen U. Two evolutionarily conserved repression domains in the Drosophila Kruppel protein differ in activator specificity. Mol Cell Biol. 1997 Aug;17(8):4820– 9.

50. Zuo P, Stanojević D, Colgan J, Han K, Levine M, Manley JL. Activation and repression of transcription by the gap proteins hunchback and Krüppel in cultured Drosophila cells. Genes Dev. 1991 Feb;5(2):254–64.

51. Ilsley GR, Fisher J, Apweiler R, De Pace AH, Luscombe NM. Cellular resolution models for even skipped regulation in the entire Drosophila embryo. eLife. 2013 Aug 6;2:e00522.

52. Jaeger J, Monk N. Bioattractors: dynamical systems theory and the evolution of regulatory processes. J Physiol (Lond). 2014 Jun 1;592(11):2267–81.

53. Jutras-Dubé L, El-Sherif E, François P. Geometric models for robust encoding of dynamical information into embryonic patterns. eLife. 2020 Aug 10;9.

54. Corson F, Siggia ED. Geometry, epistasis, and developmental patterning. Proc Natl Acad Sci USA. 2012 Apr 10;109(15):5568–75.

55. Rogers KW, Schier AF. Morphogen gradients: from generation to interpretation. Annu Rev Cell Dev Biol. 2011 Jul 29;27:377–407.

56. Simsek MF, Özbudak EM. Patterning principles of morphogen gradients. Open Biol. 2022 Oct 19;12(10):220224.

57. Zhu Y, Qiu Y, Chen W, Nie Q, Lander AD. Scaling a Dpp Morphogen Gradient through Feedback Control of Receptors and Co-receptors. Dev Cell. 2020 Jun 22;53(6):724-739.e14.

58. Bollenbach T, Pantazis P, Kicheva A, Bökel C, González-Gaitán M, Jülicher F. Precision of the Dpp gradient. Development. 2008 Mar;135(6):1137–46.

59. Stapornwongkul KS, Vincent J-P. Generation of extracellular morphogen gradients: the case for diffusion. Nat Rev Genet. 2021 Jun;22(6):393–411.

60. Lord ND, Carte AN, Abitua PB, Schier AF. The pattern of nodal morphogen signaling is shaped by co-receptor expression. eLife. 2021 May 26;10.

61. Boos A, Distler J, Rudolf H, Klingler M, El-Sherif E. A re-inducible gap gene cascade patterns the anterior-posterior axis of insects in a threshold-free fashion. eLife. 2018 Dec 20;7.

62. Rudolf H, Zellner C, El-Sherif E. Speeding up anterior-posterior patterning of insects by differential initialization of the gap gene cascade. Dev Biol. 2020 Apr 1;460(1):20–31.

63. El-Sherif E, Averof M, Brown SJ. A segmentation clock operating in blastoderm and germband stages of Tribolium development. Development. 2012 Dec 1;139(23):4341–6.

64. El-Sherif E, Zhu X, Fu J, Brown SJ. Caudal regulates the spatiotemporal dynamics of pair-rule waves in Tribolium. PLoS Genet. 2014 Oct 16;10(10):e1004677.

65. Ebisuya M, Briscoe J. What does time mean in development? Development. 2018 Jun 26;145(12).

66. Dessaud E, Yang LL, Hill K, Cox B, Ulloa F, Ribeiro A, et al. Interpretation of the sonic hedgehog morphogen gradient by a temporal adaptation mechanism. Nature. 2007 Nov 29;450(7170):717–20.

67. Balaskas N, Ribeiro A, Panovska J, Dessaud E, Sasai N, Page KM, et al. Gene regulatory logic for reading the Sonic Hedgehog signaling gradient in the vertebrate neural tube. Cell. 2012 Jan 20;148(1–2):273–84.

68. Perez-Carrasco R, Barnes CP, Schaerli Y, Isalan M, Briscoe J, Page KM. Combining a Toggle Switch and a Repressilator within the AC-DC Circuit Generates Distinct Dynamical Behaviors. Cell Syst. 2018 Apr 25;6(4):521-530.e3.

69. Panovska-Griffiths J, Page KM, Briscoe J. A gene regulatory motif that generates oscillatory or multiway switch outputs. J R Soc Interface. 2013 Feb;10(79):20120826.

70. Schroeder MD, Greer C, Gaul U. How to make stripes: deciphering the transition from non-periodic to periodic patterns in Drosophila segmentation. Development. 2011 Jul;138(14):3067–78.

71. Goto T, Macdonald P, Maniatis T. Early and late periodic patterns of even skipped expression are controlled by distinct regulatory elements that respond to different spatial cues. Cell. 1989 May 5;57(3):413–22.

72. Akam M. Drosophila development: making stripes inelegantly. Nature. 1989 Sep 28;341(6240):282–3.

73. Lynch JA, El-Sherif E, Brown SJ. Comparisons of the embryonic development of Drosophila, Nasonia, and Tribolium. Wiley Interdiscip Rev Dev Biol. 2012 Feb;1(1):16–39.

74. Clyde DE, Corado MSG, Wu X, Paré A, Papatsenko D, Small S. A self-organizing system of repressor gradients establishes segmental complexity in Drosophila. Nature. 2003 Dec 18;426(6968):849–53.

75. Durston A, Wacker S, Bardine N, Jansen H. Time space translation: a hox mechanism for vertebrate a-p patterning. Curr Genomics. 2012 Jun;13(4):300–7.

76. Palmeirim I, Henrique D, Ish-Horowicz D, Pourquié O. Avian hairy gene expression identifies a molecular clock linked to vertebrate segmentation and somitogenesis. Cell. 1997 Nov 28;91(5):639–48.

77. Kohwi M, Doe CQ. Temporal fate specification and neural progenitor competence during development. Nat Rev Neurosci. 2013 Nov 20;14(12):823–38.

78. Naidu VG, Zhang Y, Lowe S, Ray A, Zhu H, Li X. Temporal progression of Drosophila medulla neuroblasts generates the transcription factor combination to control T1 neuron morphogenesis. Dev Biol. 2020 May 19;

79. Filippopoulou K, Konstantinides N. Evolution of patterning. FEBS J. 2024 Feb;291(4):663–71.

80. Sarrazin AF, Peel AD, Averof M. A segmentation clock with two-segment periodicity in insects. Science. 2012 Apr 20;336(6079):338–41.

81. Oates AC, Morelli LG, Ares S. Patterning embryos with oscillations: structure, function and dynamics of the vertebrate segmentation clock. Development. 2012 Feb;139(4):625–39.

82. Pourquié O. The segmentation clock: converting embryonic time into spatial pattern. Science. 2003 Jul 18;301(5631):328–30.

83. McQueen C, Towers M. Establishing the pattern of the vertebrate limb. Development. 2020 Sep 11;147(17).

84. Roensch K, Tazaki A, Chara O, Tanaka EM. Progressive specification rather than intercalation of segments during limb regeneration. Science. 2013 Dec 13;342(6164):1375–9.

85. Beck MT, Váradi ZB. One, Two and Three-dimensional Spatially Periodic Chemical Reactions. Nature. 1972 Jan 3;

86. Rohde LA, Bercowsky-Rama A, Valentin G, Naganathan SR, Desai RA, Strnad P, et al. Cell-autonomous timing drives the vertebrate segmentation clock’s wave pattern. 2024 Feb 13;

87. Mau C, Rudolf H, Strobl F, Schmid B, Regensburger T, Palmisano R, et al. How enhancers regulate wavelike gene expression patterns. eLife. 2023 Jul 11;12.

88. El-Sherif E, Levine M. Shadow enhancers mediate dynamic shifts of gap gene expression in the drosophila embryo. Curr Biol. 2016 May 9;26(9):1164–9.

89. Clark E, Akam M. Odd-paired controls frequency doubling in Drosophila segmentation by altering the pair-rule gene regulatory network. eLife. 2016 Aug 15;5.

90. Koromila T, Gao F, Iwasaki Y, He P, Pachter L, Gergen JP, et al. Odd-paired is a pioneer-like factor that coordinates with Zelda to control gene expression in embryos. eLife. 2020 Jul 23;9.

91. Soluri IV, Zumerling LM, Payan Parra OA, Clark EG, Blythe SA. Zygotic pioneer factor activity of Odd-paired/Zic is necessary for late function of the Drosophila segmentation network. eLife. 2020 Apr 29;9.

92. Bothma JP, Garcia HG, Ng S, Perry MW, Gregor T, Levine M. Enhancer additivity and non-additivity are determined by enhancer strength in the Drosophila embryo. eLife. 2015 Aug 12;4.

93. Verd B, Clark E, Wotton KR, Janssens H, Jiménez-Guri E, Crombach A, et al. A damped oscillator imposes temporal order on posterior gap gene expression in Drosophila. PLoS Biol. 2018 Feb 16;16(2):e2003174.

94. Manu, Surkova S, Spirov AV, Gursky VV, Janssens H, Kim A-R, et al. Canalization of gene expression and domain shifts in the Drosophila blastoderm by dynamical attractors. PLoS Comput Biol. 2009 Mar 13;5(3):e1000303.

95. François P, Hakim V, Siggia ED. Deriving structure from evolution: metazoan segmentation. Mol Syst Biol. 2007 Dec 18;3:154.

96. François P, Siggia ED. Predicting embryonic patterning using mutual entropy fitness and in silico evolution. Development. 2010 Jul;137(14):2385–95.

97. Dar RD, Razooky BS, Singh A, Trimeloni TV, McCollum JM, Cox CD, et al. Transcriptional burst frequency and burst size are equally modulated across the human genome. Proc Natl Acad Sci USA. 2012 Oct 23;109(43):17454–9.

98. Cho W-K, Jayanth N, English BP, Inoue T, Andrews JO, Conway W, et al. RNA Polymerase II cluster dynamics predict mRNA output in living cells. eLife. 2016 May 3;5.

99. Sabari BR, Dall’Agnese A, Boija A, Klein IA, Coffey EL, Shrinivas K, et al. Coactivator condensation at super-enhancers links phase separation and gene control. Science. 2018 Jul 27;361(6400):eaar3958.

100. Hamamoto K, Fukaya T. Molecular architecture of enhancer-promoter interaction. Curr Opin Cell Biol. 2022 Feb 12;74:62–70.

